# Whole genome phylogenies reflect long-tailed distributions of recombination rates in many bacterial species

**DOI:** 10.1101/601914

**Authors:** Thomas Sakoparnig, Chris Field, Erik van Nimwegen

## Abstract

Although homologous recombination is accepted to be common in bacteria, so far it has been challenging to accurately quantify its impact on genome evolution within bacterial species. We here introduce methods that use the statistics of single-nucleotide polymorphism (SNP) splits in the core genome alignment of a set of strains to show that, for many bacterial species, recombination dominates genome evolution. Each genomic locus has been overwritten so many times by recombination that it is impossible to reconstruct the clonal phylogeny and, instead of a consensus phylogeny, the phylogeny typically changes many thousands of times along the core genome alignment.

We also show how SNP splits can be used to quantify the relative rates with which different subsets of strains have recombined in the past. We find that virtually every strain has a unique pattern of frequencies with which its lineages have recombined with those of other strains, and that the relative rates with which different subsets of strains share SNPs follow long-tailed distributions. Our findings show that bacterial populations are neither clonal nor freely recombining, but structured such that recombination rates between different lineages vary along a continuum spanning several orders of magnitude, with a unique pattern of rates for each lineage. Thus, rather than reflecting clonal ancestry, whole genome phylogenies reflect these long-tailed distributions of recombination rates.

## Introduction

The only illustration that appears in Darwin’s Origin of Species [1] is of a phylogenetic tree. Indeed, the tree has become the archetypical concept representing biological evolution. Since every biological cell that has ever lived was the result of a cell division, all cells are connected through cell divisions in a giant tree that stretches all the way back to the earliest cells that existed on earth. Thus, the study of biological evolution in some sense corresponds to the study of the structure of this giant cell-division tree.

It is therefore natural that the first step in the analysis of a set of related biological sequences is to reconstruct the phylogenetic tree that reflects the cell division history of the sequences, i.e their ‘ancestral phylogeny’. Once the ancestral relationships between the sequences are known, the evolution of the sequences can then be modeled along the branches of this tree. Indeed, virtually all models of evolutionary dynamics are formulated as occurring along the branches of a tree, and many mathematical and computational methods have been developed for their inference, see e.g. [2,3]. This strategy has been employed from the earliest days of sequence analysis [4] and is almost invariably applied in the analysis of microbial genome sequences, which is the main topic of this work.

A second key concept in models of evolutionary dynamics is the idea of a ‘population’ of organisms that are mutually competing for resources and that, for purposes of mathematical modeling, can be considered exchangeable in the sense that they are subjected to the same environment. Indeed, populations of exchangeable individuals form the basis of almost all mathematical population genetics models (see e.g. [5]), including coalescent models for phylogenies [6]. Although it is of course well recognized that, in the real world, populations are structured into sub-populations with varying degrees of interaction between them, population genetics models almost by definition assume that at some level there are sub-populations of exchangeable individuals sharing a common environment.

In this paper we present evidence that we believe challenges the usefulness of applying these two concepts for describing genome evolution in prokaryotes. First, we find that for most bacterial species recombination is so frequent that, within an alignment of strains, each genomic locus has been overwritten by recombination many times and the phylogeny typically changes tens of thousands of times along the genome. Moreover, for most pairs of strains, none of the loci in their pairwise alignment derives from their ancestor in the ancestral phylogeny, and the vast majority of genomic differences result from recombination events, even for very close pairs. Consequently, the ancestral phylogeny cannot be reconstructed from the genome sequences using currently available methods and, more generally, the strategy of modeling microbial genome evolution as occurring along the branches of an ancestral phylogeny breaks down.

Second, we show that the structure represented in whole genome phylogenies of microbial strains does not reflect ancestry, but instead the relative rates with which different lineages have recombined in the past. Whereas almost every short genomic segment follows a different phylogeny, these phylogenies are not uniformly randomly sampled from all possible phylogenies, but sampled from highly biased distributions. In particular, the relative frequencies with which particular sub-clades of strains occur in the phylogenies at different loci follow roughly power-law distributions and each strain has a distinct distribution of co-occurrence frequency with the other strains. Since almost every strain has a unique ‘finger print’ of recombination rates with the lineages of other strains, the assumption that at some level the strains can be considered as exchangeable members of a population, also fundamentally breaks down.

The structure of the paper is as follows. To present our analyses, we will focus on a collection of 91 wild *E. coli* strains that were isolated over a short period from a common habitat [7]. Using these strains, we introduce the main puzzle of bacterial whole genome phylogeny: although the phylogenies of individual genomic loci are all distinct, the phylogeny inferred from any large collection of genomic loci converges to a common structure, e.g. [8–10]. This convergence has led to many approaches where it is assumed that the phylogeny reconstructed from genome-wide alignments or pairwise divergences must represent the clonal phylogeny, and that recombination can be detected and quantified by measuring deviations from this global phylogeny, e.g. [11–15]. However, as far as we are aware, there are no rigorous justifications for assuming that this global phylogeny corresponds to the clonal phylogeny. Given that it is currently unclear what this global phylogeny represents, it is also unclear whether deviations from this global phylogeny can be used to characterize recombination. One main goal of this work is to introduce methods for quantifying the role of recombination without having to rely on assumptions that have not been substantiated.

We first study recombination by studying pairs of strains, extending a recent approach by Dixit et al. [16] to model each pairwise alignment as a mixture of ancestrally inherited and recombined regions. We show that, as the distance to the pair’s common ancestor increases, the fraction of the genome covered by recombined segments increases, and at some pairwise distance all clonally inherited DNA disappears. Importantly, this distance is far below the typical divergence of pairs of strains such that for the vast majority of pairs, none of the DNA in their genome alignment stems from their common ancestor.

Much of the new analysis methodology that we introduce is based on bi-allelic SNPs (which constitute almost all SNPs in the core alignment). Although bi-allelic SNPs have been studied to estimate the number of recombinations along alignments of sexually reproducing species [17], they have received relatively little attention in the study of prokaryotic genomes. We show that almost every bi-allelic SNP corresponds to single nucleotide substitution in the history at the corresponding genomic position, so that the subset of individuals sharing a common nucleotide must form a clade in the phylogeny at that position. We show various ways in which these bi-allelic SNPs can be used to investigate which SNPs are consistent with given phylogenies, and each other, and use them to quantify the amount of phylogenetic variation along the alignment. We use these SNPs to show there is no consensus phylogeny, that the phylogeny changes every few SNPs along the core phylogeny, and to estimate a lower bound on the ratio of recombination to mutation events in a genome alignment.

To obtain an additional assessment of the validity of our methods, we not only apply our methods to the real data from the *E. coli* strains but also to data from simulations of a simple evolutionary dynamics in which genomes of a well-mixed population of fixed size *N* evolve under reproduction, (neutral) mutation at a fixed rate *μ* per base per generation, and homologous recombination at a rate *ρ* per base per generation. By comparing the known ground truth for the simulated data with the results of our methods, we confirm the accuracy of our methods on the simulated data. We also use results of our methods on simulated data to help elucidate the precise meaning of several statistics that we calculate.

Applying the methods and statistics that we developed for *E. coli* to a set of other bacterial species including: *B. subtilis, H. pylori, M. tuberculosis, S. enterica*, and *S. aureus*, we show that, with the exception of *M. tuberculosis* where all strains are very closely related and no pair has yet been fully recombined, all other species follow the same general behavior as *E. coli.* Thus, for all but one of these bacterial species, there is no common or consensus phylogeny, but many thousands of different phylogenies along the core genome.

To explain how a robust core genome phylogeny can emerge in spite of the fact that a very large number of different, phylogenies occurs along the genome, we use data from human genomes as an illustration. We show that phylogenies reconstructed from large numbers of loci from human genomes also converge to a robust core phylogeny. Moreover, this phylogeny reflects the known human population structure, i.e. the relative rates with which different human lineages have recombined, and we propose that bacterial whole genome phylogenies similarly reflect the rates with which different lineages have recombined. To support this interpretation, we show how bi-allelic SNPs can be used to quantify the relative rates with which different bacterial lineages share mutations. We find that these rates follow approximately scale free distributions, indicating that there is ‘population structure’ on every scale. Finally, we define entropy profiles of the phylogenetic variability of each strain and show that these entropy profiles provide a unique phylogenetic fingerprint of almost every strain. Comparison of the various statistics we calculate between the real data and data from simulations also illustrates that simple evolutionary models of fully mixed populations cannot reproduce the statistics we observe for real genomic data. We feel that our observations necessitate a new way of thinking about how to model genome evolution in prokaryotes.

## Results

To illustrate our methods we focus on the SC1 collection of wild *E. coli* isolates that were collected in 2003 — 2004 near the shore of St. Louis river in Duluth, Minnesota [7,18]. We sequenced 91 strains from this collection together with the K12 MG1655 lab strain as a reference. In a companion paper [19] we discuss this collection in more detail and extensively analyze the evolution of gene content and phenotypes of this collection. Here we focus on sequence evolution in the core genome of these strains. Although the SC1 strains were collected from a common habitat over a short period of time, they show a remarkable diversity, with no two identical strains, all known major groups of *E. coli* represented, as well as an ‘out group’ of 9 strains that are more than 6% diverged at the nucleotide level from other *E. coli* strains (see Suppl. Fig. S1 for a phylogenetic tree constructed using maximum likelihood on the joint core genome of the SC1 strains and 189 reference strains, [19]).

### Phylogenies of individual loci disagree with the phylogeny of the core genome

To construct a core genome alignment of the SC1 strains and K12 MG1655 we used the Realphy software [20] (see Materials and Methods), resulting in a multiple alignment across all 92 strains of 2’756’541 base pairs long, with 299’077 (10.8%) of the positions exhibiting polymorphism. Although all strains are unique when their full genomes are considered, there are some groups of strains that are so close that they are identical in their core genomes, so that there are only 82 unique core genomes in total. Realphy used PhyML [21] to reconstruct a phylogeny from the core genome alignment and we will refer to this tree as the *core tree* from here on (Fig. 1).

**Figure 1.**
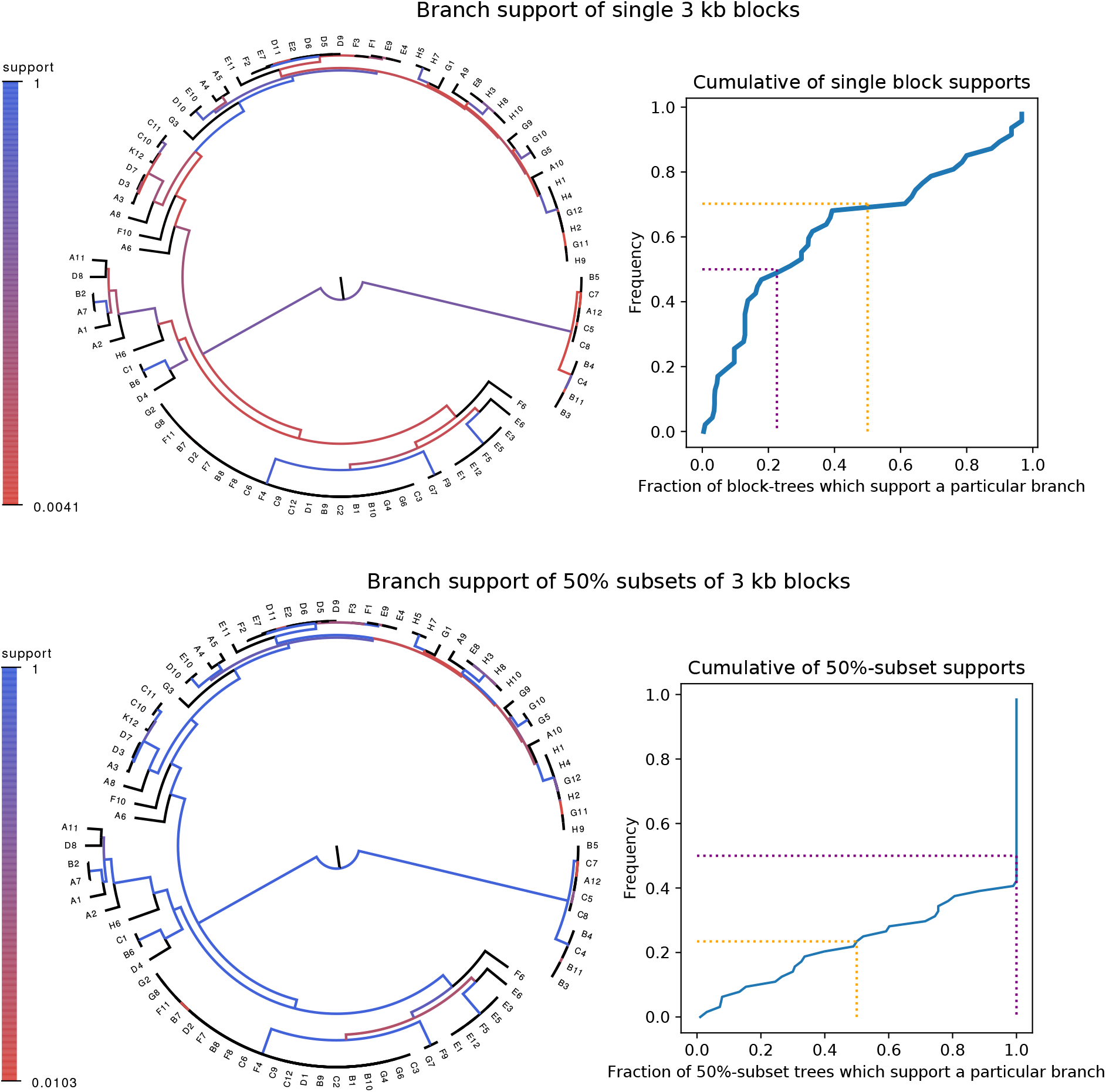
Whereas phylogenies of individual alignment blocks differ substantially from the core tree, phylogenies reconstructed from a large number of blocks are highly similar to the core tree. **Top left:** For each split (i.e. branch) in the core tree, the color indicates what fraction of the phylogenies of 3 Kb blocks support that bi-partition of the strains. **Top right:** Cumulative distribution of branch support, i.e. fraction of 3 Kb blocks supporting each branch. The dotted lines indicated show the fraction of branches that have less than 50% support (yellow) and the median support per branch (purple). **Bottom left and bottom right:** As in the top row, but now based on phylogenies reconstructed from random subsets of 50% of all 3 Kb blocks as opposed to individual blocks.

We first checked to what extent the alignments of individual genomic loci are statistically consistent with the core tree. For each 3 kilobase block of the core alignment we used PhyML to reconstruct a phylogeny and then compared its log-likelihood with the log-likelihood that can be obtained when the phylogeny is constrained to have the topology of the core tree. We find that essentially all 3 Kb alignment blocks reject the core tree topology (Suppl. Fig. S2, left panel). Moreover, each alignment block rejects the topologies that were constructed from all other alignment blocks (Suppl. Fig. S2, right panel).

Although it thus appears that the phylogeny at each genomic locus is statistically significantly distinct, it is still possible that all these phylogenies are highly similar. In order to quantify the differences between the core tree and the phylogenies of 3Kb blocks we calculated, for each split in the core tree, the fraction of 3Kb blocks for which the same split occurred in the phylogeny reconstructed from that alignment block. As shown in the top row of Fig. 1, the phylogenies of individual blocks differ substantially from the core tree: roughly two-thirds of the splits in the core tree occur in less than half of 3Kb block phylogenies and half of the core tree splits occur in less than a quarter of all 3Kb block phylogenies. Especially the splits higher up in the core tree do not occur in the large majority of block phylogenies.

These observations are not particularly novel. There is by now a vast and sometimes contentious literature on the role of recombination in prokaryotic genome evolution which is beyond the scope of this article to review. We thus focus on a few key points that are central to the questions and methods we study here. First, systematic studies of complete microbial genomes have shown that horizontal gene transfer is relatively common and can significantly affect phylogenies of individual loci, e.g. [22,23], and many studies have observed that different genomic loci support different phylogenies, e.g. [24]. Such observations caused some researchers to question whether trees can be meaningfully used to describe genome evolution [25]. Interestingly, it has recently been shown that recombination can play a major role in genome evolution even within relatively short term laboratory evolution experiments [26].

In spite of this, many researchers in the field feel that a major phylogenetic backbone can still be extracted from genomic data. For example, it has been observed that, whenever a phylogeny is reconstructed from the alignments of a large number of genomic loci, one obtains the same or highly similar phylogenies, e.g. [8–10]. We also observe this behavior for our strains. Phylogenies reconstructed from a random sample of 50% of all 3Kb blocks look highly similar to the core tree, i.e. with two thirds of the core tree’s splits occurring in *all* phylogenies (Fig. 1, bottom row).

How should we interpret this convergence of phylogenies to the core phylogeny as increasing numbers of genomic loci are included? One interpretation that has been proposed, is that once a large number of genomic segments is considered, effects of horizontal transfer are effectively averaged out, and the phylogeny that emerges corresponds to the clonal ancestry of the strains, e.g. [8, 11]. Indeed, it has become quite common for researchers to detect and quantify recombination using methods that compare local phylogenetic patterns with an overall reference phylogeny constructed from the entire genome, e.g. [11–15, 27]. However, the validity of such approaches rests on the assumption that this reference phylogeny really represents the clonal phylogeny, and it is currently unclear whether this is justified.

Indeed, some recent studies have argued that recombination is so common in some bacterial species that it is impossible to meaningfully reconstruct the clonal ancestry from the genome sequences, and that these species should be considered freely recombining, e.g. [28]. However, if members of the species are freely recombining, one would expect the core tree to take on a star-like structure as opposed to the clear and consistent phylogenetic structure that phylogenies converge to as more genomic regions are included in the analysis. Addressing this puzzle is one of the topics of this work.

### Quantifying recombination through analysis of pairs of strains

As a first analysis of the impact of recombination, we follow an approach recently proposed by Dixit et al. based on the pairwise comparison of strains [16]. The simplest measure of the distance between a pair of strains is their nucleotide divergence, i.e. the fraction of mismatching nucleotides in the core genome alignment of the two strains. For pairs of strains with very low divergence, e.g. D6 and F2 with divergence 4 × 10 ^4^ (Fig. 2A), the effects of recombination are almost directly visible in the pattern of SNP density along the genome. While the SNP density is very low along most of the genome, i.e. 0 — 2 SNPs per kilobase, there are a few segments, typically tens of kilobases long, where the SNP density is much higher and similar to the typical SNP density between random pairs of *E. coli* strains, i.e. 10 — 30 SNPs per kilobase. These high SNP density regions almost certainly result from horizontal transfer events in which a segment of DNA from another *E. coli* strain, for example carried by a phage, made it into one of the ancestor cells of this pair, and was incorporated into the genome through homologous recombination. For pairs of increasing divergence, e.g. the pair C10-D7 with divergence 0.002 in Fig. 2B, the frequency of these recombined regions increases, until eventually most of the genome is covered by such regions (pair D6-H10 in Fig. 2C).

**Figure 2.**
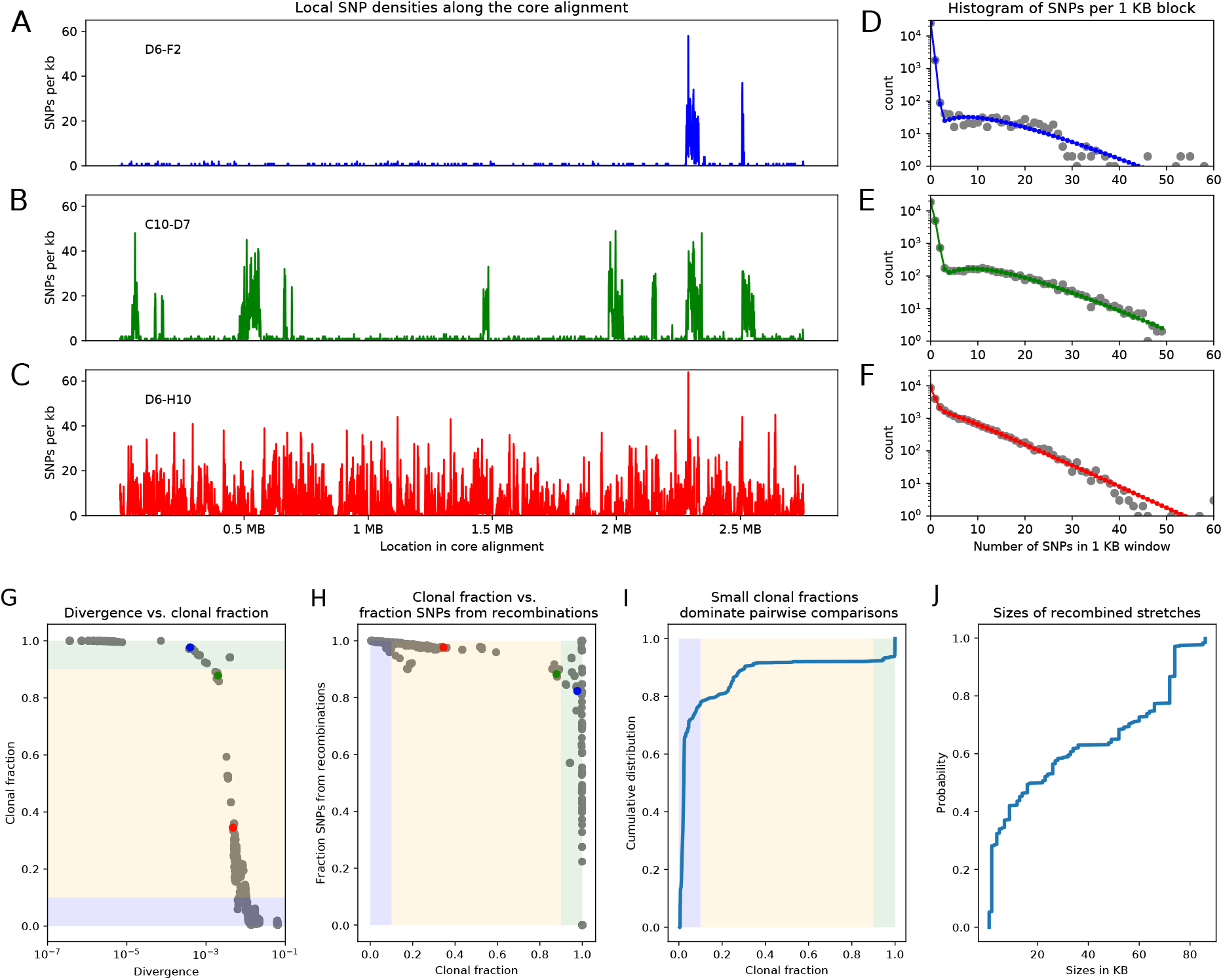
Summary statistics of pairwise analysis for the SC1 strains. **A-C**: SNP densities (SNPs per kilobase) along the core genome for three pairs of strains at overall nucleotide divergences of 4 × 10^-4^ (D6-F2), 0.002 (C10-D7), and 0.0048 (D6-H10). **D-F**: Corresponding histograms for the number of SNPs per kilobase (dots) together with fits of the mixture model for D6-F2 (blue), C10-D7 (green), and D6-H10 (red). Note the vertical axis is on a logarithmic scale. **G**: For each pair of strains (dots), the fraction of the genome that was inherited clonally is shown as a function of the nucleotide divergence of the pair, shown on a logarithmic scale. The three pairs that were shown in panels A-F are shown as the blue, green, and red dots. The light green, yellow, and blue segments show strains that are mostly clonal, a mixture of clonal and recombined, and fully recombined, respectively. **H**: Fraction of all SNPs that lie in recombined regions as a function of the clonally inherited fraction of the genome. **I**: Cumulative distribution of the clonal fractions of the pairs. **J**: Cumulative distribution of the lengths of recombined segments for pairs that are in the mostly clonal regime. The mean length of recombined regions is 31’197, with first quartile 2000, median 19’500, and third quartile 66’000.

For close pairs, the histograms of SNP densities also clearly separate into two components: a majority of clonally inherited regions with up to at most 3 SNPs per kilobase, and a long tail of recombined regions with up to 50 or 60 SNPs per kilobase (Fig. 2D-E). As explained in the methods, the distributions of SNP densities is well described by a mixture of a Poisson distribution for the clonally inherited regions plus a negative binomial for the recombined regions (solid line fits in Fig. 2D-F). In this way we can estimate, for each pair of strains, the fraction of the genome that is clonally inherited, and the number of SNPs that fall in clonally inherited versus recombined regions. In addition, to estimate the lengths of the recombined fragments we focused on very close pairs for which recombination events are sparse enough so that overlapping events are very unlikely, and used a Hidden Markov model to estimate the distribution of lengths of recombined regions (see Materials and Methods). We find that recombined blocks are typically in the range of 10 — 70 kilobases long with a median of almost 19.5 kilobases (Fig. 2J).

From this analysis we see that, whenever the pairwise divergence is less than 0.001, the large majority of blocks is clonally inherited, which is indicated as the light-green segment in Fig. 2G. However, over a narrow range of divergence between 0.001 and 0.01 the fraction of clonally inherited DNA drops dramatically (yellow segment in Fig. 2G) and at a divergence of about 0.014 essentially the entire alignment has been overwritten by recombination and all clonally inherited DNA is lost (blue segment in Fig. 2G). Notably, 80% of all strain pairs lie in this fully recombined regime (Fig. 2I). Thus, for the large majority of pairs of strains, none of the DNA in their alignment derives from their clonal ancestor, making it impossible to estimate the distance to their clonal ancestor from comparing their DNA. Moreover, as shown in Fig. 2H, even for pairs that are so close that most of their genomes are clonally inherited, the large majority of the SNPs derives from the recombined regions.

For later comparison with the data on other species, we summarize our observations from the pairwise analysis by a few key statistics. First, half of the genome is recombined at a *critical divergence* of 0.0032. Second, at this critical divergence, the fraction of all SNPs that is in recombined regions is 0.95. Third, the fraction of mostly clonal pairs is 0.077, and finally, the fraction of fully recombined pairs is 0.78 (see Materials and Methods). All these statistics suggest that pairwise divergences between strains are almost entirely driven by recombination and do not reflect distances to their clonal ancestors. To understand how a consistent phylogenetic structure can still emerge when the full core genomes of all strains are compared, we need to go beyond studying pairs.

### Simulations of a simple evolutionary model including drift, mutation, and recombination

To confirm the validity of our methods and to compare observed results with results from data that derive from well understood evolutionary dynamics, we performed simulations of a relatively simple evolutionary model that includes drift, (neutral) mutation, and recombination. As described in the materials and methods, we simulated populations of constant size *N* with non-overlapping generations, a constant mutation rate *μ* per generation per base, and assuming all mutations are neutral. We also included recombination by, after each genome duplication, inserting genomic fragments from randomly chosen other individuals in the population at a rate *ρ* per position per generation. In order to make the simulation results comparable to our core genome alignment of *E. coli* strains we focused on genome alignments of a sample of *S* = 50 individuals from the population and set the mutation rate *μ* such that the fraction of polymorphic columns in the simulated alignment is similar to that observed in the real data, i.e. 10%. Apart from simulations without recombination, we performed simulations with a wide range of recombination to mutation rates from *ρ/μ* = 0.001 to *ρ/μ* = 10. In addition, we made sure to design these simulations such that a clonal tree of the sample of S = 50 genomes is drawn once according to the well-known Kingman coalescent [6], and then kept fixed in all simulations. In each simulation we explicitly tracked how many times each position in the genome was overwritten by recombination along each branch of the clonal phylogeny (see Materials and Methods).

To give an idea of the extent to which the clonal history is conserved during the evolution for recombination-to-mutation rates *ρ/μ* ranging from 0.001 to 10, Suppl. Fig. S3 shows pictures of the clonal tree with their branches colored according to the fraction of positions in the genome that were clonally inherited (i.e. not affected by recombination) along each branch. At a very low rate of recombination of *ρ/μ* = 0.001, the large majority of all positions in each branch are clonally inherited. When the ratio *ρ/μ* is raised to 0.01, evolution is still mostly clonal along most of the branches toward the leafs of the tree, but for the few long internal branches near the root that separate the major ‘clades’, most positions have already been affected by recombination. When the ratio *ρ/μ* is further raised to 0.1, almost all branches are dominated by recombination except for a few branches near the leafs. For *ρ/μ* = 0.3 or larger, almost every position in every branch has been overwritten by recombination.

To test to what extent the pair analysis introduced in the previous section correctly reflects the known ground truth for the simulations we applied the same pairwise analysis to alignments of the S = 50 sample genomes from each of the simulations. Supplementary Figure S4 shows the results, analogous to those of Fig. 2G-I, for the data from simulation with recombination to mutation ratios of *ρ/μ* = 0,0.001,0.01, 0.1, and 0.3. For the simulations without recombination, the pairwise analysis correctly infers that all of the genome is clonally inherited for all pairs. For the very low recombination rate *ρ/μ* = 0.001 the model also correctly infers that clonal evolution dominates, i.e. more than 50% of the genome is clonally inherited for all pairs, and more than 90% of the genome is clonally inherited for about 40% of all pairs. The pairwise analysis of the simulation data with recombination rate *ρ/μ* = 0.01 are also consistent with the ground truth of Suppl. Fig. S3, i.e. for half of the pairs there is a substantial fraction of clonally inherited genome, whereas for the other half of more distally related pairs recombination has already affected more than 90% of the genome. Similarly, for *ρ/μ* = 0.1 the pairwise analysis correctly infers that pairs which are not yet fully recombined have become rare, and for *ρ/μ* = 0.3 essentially all pairs have become fully recombined. The pairwise analysis of the simulated data thus paints a correct picture of the ground truth shown in Suppl. Fig. S3.

To get a more quantitative assessment of the accuracy of the pairwise analysis, Suppl. Fig. S5 directly compares the true fraction of clonally inherited genome for each pair with the estimated fraction from the pairwise analysis for the simulations with *ρ/μ* = 0.01 and *ρ/μ* = 0.1. We see that for all pairs the estimated fraction of the genome that was clonally inherited is close to the true fraction. In fact, it appears that the estimation method tends to slightly overestimate the fraction of clonally inherited genome for almost all pairs. Finally, as for the real data, we also used close pairs from the simulations with *ρ/μ* = 0.01 and *ρ/μ* = 0.1 to estimate the sizes of the recombined fragments. As shown in Suppl. Fig. S5, the sizes of the recombined regions estimated by the method are very close to the ground truth of 12kb recombination fragments.

In summary, application of the pairwise analysis to the simulation results strongly support that this pairwise analysis can accurately quantify the fraction of the genome that was affected by recombination for each pair, as well as estimate the lengths of the recombined segments.

### SNPs in the core genome alignment correspond to splits in the local phylogeny

Whereas there may not be a single phylogeny that captures the evolution of our genomes, each *single position* in the core genome alignment, i.e. each alignment column, will have evolved according to some phylogenetic tree. A key observation is that our set of strains is sufficiently closely related that there are almost no alignment for which more than one substitution occurred in its evolutionary history. In particular, of the almost 2.5 million columns in the core genome alignment, almost 90% show an identical nucleotide for all 92 strains, i.e only 10.85% are polymorphic. Moreover, almost all of these SNP columns are bi-allelic, i.e. for 93.6% of the SNPs only 2 nucleotides appear, 6.3% have 3 nucleotides, and in 0.2% all 4 nucleotides occur. These statistics strongly suggest that most positions have not undergone any substitutions, and that columns with multiple substitutions are rare. Notably, these statistics are still inflated due to the occurrence of an outgroup of 9 strains that is far removed from the other strains (the clade from B5 to B3 visible on the right in Fig. 1). We observe that almost 36% of all SNPs correspond to SNPs in which all 9 strains of this outgroup have one nucleotide, and all other 83 strains have another nucleotide. If we remove the outgroup from our alignment, the fraction of SNPs in the alignment drops from 10.85% to 6.7%, and the fraction of SNPs that are bi-allelic increases to 95.5%.

Whenever a bi-allelic SNP corresponds to a single substitution in the evolutionary history of the position, the SNP pattern provides an important piece of information about the phylogeny at that position in the alignment: whatever this phylogeny is, it must contain a split, i.e. a branch bi-partitioning the set of strains, such that all strains with one letter occur on one side of the split, and all strains with the other letter on the other side (Fig. 3).

**Figure 3.**
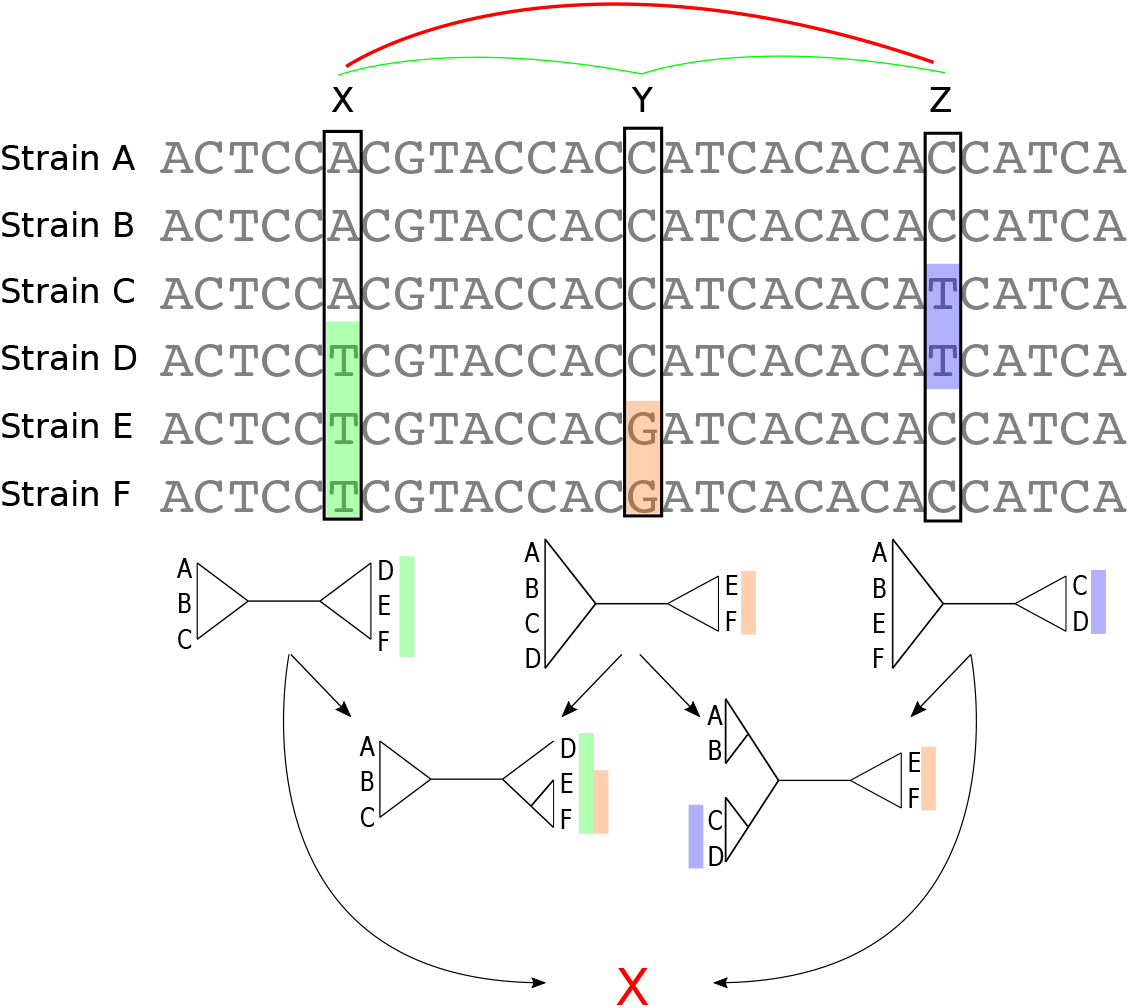
SNP columns correspond to phylogeny splits. A segment of a multiple alignment of 6 strains containing 3 bi-allelic SNPs, *X, Y*, and *Z*. Assuming that each SNP corresponds to a single substitution in the evolutionary history of the position, each SNP constrains the local phylogeny to contain a particular split, i.e. bi-partition of the strains, as illustrated by the 3 diagrams immediately below each SNP. In this example, the neighboring pairs of SNPs (*X, Y*) and (*Y, Z*) are both consistent with a common phylogeny and can be used to further resolve the phylogeny in the local segment of the alignment as shown in the second row with two diagrams. However, SNPs *X* and *Z* are mutually inconsistent with a common phylogeny (red cross at the bottom) indicating that somewhere between *X* and *Z* at least one recombination event must have occurred.

As illustrated in Fig. 3, pairs of SNPs can either be consistent with a common phylogeny, i.e. columns *X* and *Y* or columns *Y* and *Z*, or they can be inconsistent with a common phylogeny, i.e. columns *X* and *Z*. The pairwise comparison of SNP columns for consistency with a common phylogeny is known as the four-gamete test and is a very commonly used method in the literature on sexual species, where it has been used, for example, to give a lower bound on the number of recombination events in an alignment [17]. However, so far it has rarely been used for quantifying recombination in bacteria ([29] and [30] being the only exceptions we are aware of). In the rest of this paper we show how analysis of bi-allelic SNPs (which from now on we will just call SNPs) can be systematically used to quantify not only the overall amount of recombination in alignments, but also the relative rates with which different lineages have recombined.

However, since all these analyses assume that bi-allelic SNPs correspond to single substitutions, it is important to quantify how accurate this assumption is. In particular, some apparent SNPs might correspond to sequencing errors rather than true substitutions and, more importantly, some bi-allelic SNPs may correspond to multiple substitution events (often called homoplasies). In the Materials and Methods we show that sequencing errors must be so rare that they can be safely neglected. In addition, to estimate the fraction of bi-allelic SNPs that correspond to homoplasies we analyzed the frequencies of columns with 1, 2, 3 and 4 different nucleotides using a simple substitution model, separately analyzing positions that are under least selection (third positions of 4-fold degenerate codons) and positions under most selection (second positions in codons), and either including or excluding the outgroup (see Materials and Methods). These analyses indicate that only 2 — 6% of bi-allelic SNPs correspond to homoplasies. We confirmed the accuracy of this estimation procedure using simulation data, for which the true fraction of homoplasies is 2.56% and our simple method estimates 2.43%.

In addition, to put an upper bound on how much our results could be affected by a small fraction of homoplasies, we developed a method that removes, from the core genome alignment, a given fraction of alignment columns that exhibit most inconsistencies with alignment columns in their neighborhood (see Materials and Methods), and checked how much the various statistics that we calculate are altered when we remove either 5% or 10% of such potentially homoplasic positions. Finally, we also apply each of our analyses to data from simulations with known ground truth, to further validate the accuracy of our methods.

### SNP statistics are inconsistent with a single consensus phylogeny

One of the key uses of a phylogeny is in explaining the differences between the strains by an evolutionary dynamics that takes place along the branches of the phylogeny. However, for such an approach to apply, it is important that the large majority of evolutionary changes were indeed introduced along the branches of the phylogeny.

That there are significant differences between the SNP statistics as predicted by the core tree, and the actual SNP statistics observed in the data is already evident from the fact that the observed pairwise divergences between strains are systematically less than their divergences along the branches of the core phylogeny (Suppl Fig. S6, left panel). In addition, the observed number of SNPs corresponding to each branch of the core tree are up to 100-fold lower than number of SNPs that are predicted to occur on the branch (Suppl Fig. S6, right panel).

To test to what extent the observed evolutionary changes occurred along the branches of the core tree, we calculated the fraction of observed SNP splits that fall on the branches of the core phylogeny. Since each branch in the core tree corresponds to a split, we calculated what fraction of SNPs correspond to a branch in the core tree, and what fraction are inconsistent with the core tree. Overall, 58% of the SNPs that are shared by at least 2 strains correspond to a branch of the core tree, whereas 42% clash with it (SNPs that occur in only a single strain are consistent with any phylogeny). However, this relatively high fraction results almost entirely from SNPs on the single branch connecting the outgroup to the other strains, which is responsible for almost 36% of all SNPs. When the outgroup is removed, only 27.4% of all SNPs are consistent with the core tree. Since the core tree was constructed using a maximum likelihood approach that assumes the entire alignment follows one common tree, we investigated to what extent the number of tree supporting SNPs can be improved by specifically constructing a tree to maximize the number of supporting SNPs (see Materials and Methods). However, such trees only marginally improve the number of supporting SNPs by 0.1%.

Since the overall fraction of SNPs that correspond to branches of the core phylogeny is dominated by a few very common SNP patterns, it is more informative to assess to what extent each individual branch of the core tree is consistent with the observed SNPs. We thus calculated, for each branch in the core tree, the number of supporting SNPs S that match the split, and the number of clashing SNPs *C* that are inconsistent with the split, to calculate the fraction *f* = *S*/(*S + C*) of SNPs supporting the branch. Figure 4 shows that, for two-thirds of the branches, there are more clashing than supporting SNPs. Moreover, for half of the branches in the core tree, the fraction of supporting SNPs is less than 5%, i.e. there are at least 20-fold more clashing than supporting SNP columns. To check to what extent homoplasies may affect these statistics, we calculated the same statistics on core genome alignments from which either 5% or 10% of potentially homoplasic sites were removed, and observed that the distribution of SNP support is almost unchanged (Suppl. Fig. S7).

**Figure 4.**
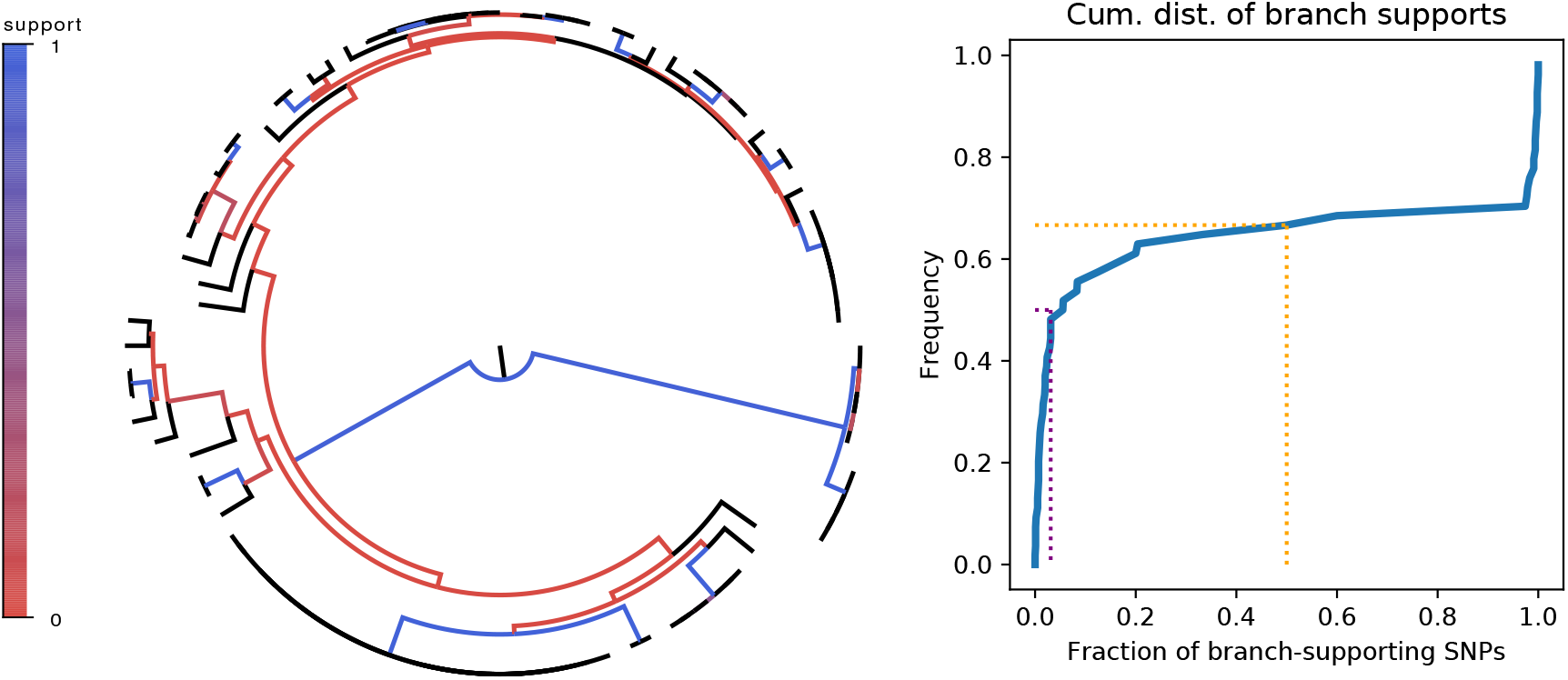
SNP support of the branches of the core genome tree. **Left Panel**: Fraction of supporting versus clashing SNPs for each branch of the core tree. **Right panel**: Cumulative distribution of the fraction of supporting SNPs across all branches. The purple and orange dotted lines show the median and the frequency of branches with 50% or less support, respectively.

To further elucidate the meaning of these SNP support distributions, we calculated the same statistics for the simulations with different recombination rates, including on alignments where 5% or 10% of potentially homoplasic columns were removed (Suppl. Fig. S8). In general the removal of 5% or even 10% of potentially homoplasic sites has only a minor effect on the distribution of SNP support except for the simulations without recombination. When there is no recombination all inconsistencies are due to homoplasies, and we indeed see that all branches are fully supported after 5% of potential homoplasic sites have been removed. For a very small recombination rate of *ρ/μ* = 0.001 most branches in the core tree still have strong SNP support but already at a recombination rate of *ρ/μ* = 0.01 more than 80% of the branches are supported by less than half of the informative SNPs, and half of the branches have less than 20% support. It is instructive to compare this result with the fractions of the genome that are clonally inherited along each branch, i.e. the bottom left panel in Suppl. Fig. S3, and the pair statistics (middle row of Suppl. Fig. S4) for this simulation. We see that, although there is some recombination in most branches, most of the genome is clonally inherited for all but the long inner branches. However, even though most of the genome is clonally inherited along most branches, for most pairs the large majority of the SNPs derive from recombination (middle panel in Suppl. Fig. S4). This means that, at *ρ/μ* = 0.01, we’re in a parameter regime where most of the genome is still clonally inherited, and the clonal tree can thus also be reconstructed from the data, but the large majority of the differences between strains already derive from recombination events. This shows that, even if recombination is rare enough that the clonal tree can be successfully reconstructed from the genome sequences, it might already be incorrect to assume most genomic changes were introduced along the branches of this clonal phylogeny.

For recombination rates of *ρ/μ* = 0.1 or larger recombination dominates completely in that there are essentially no branches with significant SNP support. The distribution of support for *E. coli* does not look like any of the distributions from the simulations, but could be described as a hybrid of one third branches having strong clonal support and two thirds branches that are dominated by recombination. Besides the branch to the outgroup, the supported branches all correspond to branches toward groups of highly similar strains near the bottom of the tree. We thus wondered if it would be possible to construct well supported subtrees for clades of closely-related strains near the bottom of the tree. We devised a method that builds subtrees bottom-up by iteratively fusing clades so as to minimize the number of clashing SNPs at each step (see Materials and Methods and Suppl. Fig S9, left panel). As shown in Suppl. Fig. S9, while the fraction of clashing SNPs is initially low, it rises quickly as soon as the average divergence within the reconstructed subtrees exceeds 10^-4^, which is more than 100-fold below the typical pairwise distance between *E. coli* strains. Thus, while some groups of very closely-related strains that have a recent common ancestor can be unambiguously identified, only a minute fraction of the overall sequence divergence falls within these groups, and the bulk of the sequence variation between the strains is not consistent with a single phylogeny.

To assess whether any of these statistics could be skewed due to a few strains with aberrant behavior, we also investigated whether SNPs are more consistent with a dominant phylogeny when we do not consider all strains, but only subsets of the strains. We focused on the smallest subsets of strains that have meaningfully different phylogenetic tree topologies. For a quartet of strains (*I, J, M, N*), there are 3 possibly binary trees, i.e with (*I, J*) and (*M, N*) nearest neighbors, with (*I, M*) and (*J, N*), or with (*I, N*) and (*J, M*) (See Suppl. Fig. S10). We selected quartets of roughly equidistant strains and checked, for each quartet, whether the SNPs clearly supported one of the tree possible topologies. However, we find that alternative topologies are always supported by a substantial fraction of the SNPs, and that for most quartets the most supported topology is supported by less than half of the SNPs (Suppl. Fig S10).

In summary, consistent with the picture that emerged from our analysis of pairs of strains, most of the differences between the *E. coli* strains did not occur along the branches of a single phylogeny. This suggests that, rather than describing the relationships between the strains by a single phylogeny, we should think of multiple different phylogenies occurring along the genome alignment.

### Phylogeny changes every few dozens of base pairs along the core alignment

So far we have analyzed SNP consistency without regard to their relative positions. We now analyze to what extent mutually consistent SNPs are clustered along the alignment. In particular, we calculate the lengths of segments along the alignment that are consistent with a single phylogeny.

We first assessed the length-scale over which phylogenies are correlated by calculating a standard linkage disequilibrium (LD) measure as a function of distance along the alignment (Fig. 5A and Materials and Methods). LD drops quickly over the first 100 base pairs and becomes approximately constant at distances beyond 200 — 300 base pairs, indicating that segments of correlated phylogenies are much shorter than the typical length of a gene. Very short linkage profiles were recently also observed in thermophilic *Cyanobacteria* isolated in Yellowstone National Park [28]. Instead of using correlation between SNPs at different distances, one can also calculate the probability for a pair of SNPs to be consistent with a common phylogeny, as a function of their genome distance [30]. As shown in Suppl. Fig. S11, like LD, pairwise compatibility of SNPs also drops quickly over the first 100 base pairs. Note, however, that even at large distances the pairwise compatibility of SNPs is close to 90%. The reason for this is that most SNPs are shared by only a small subset of strains, and as long as two SNPs are shared by non-overlapping subsets of strains, they will be compatible with a tree. In order to more efficiently detect recombination using SNP compatibility, we need to check for the mutual consistency of *all* SNPs within a given segment of the alignment.

**Figure 5.**
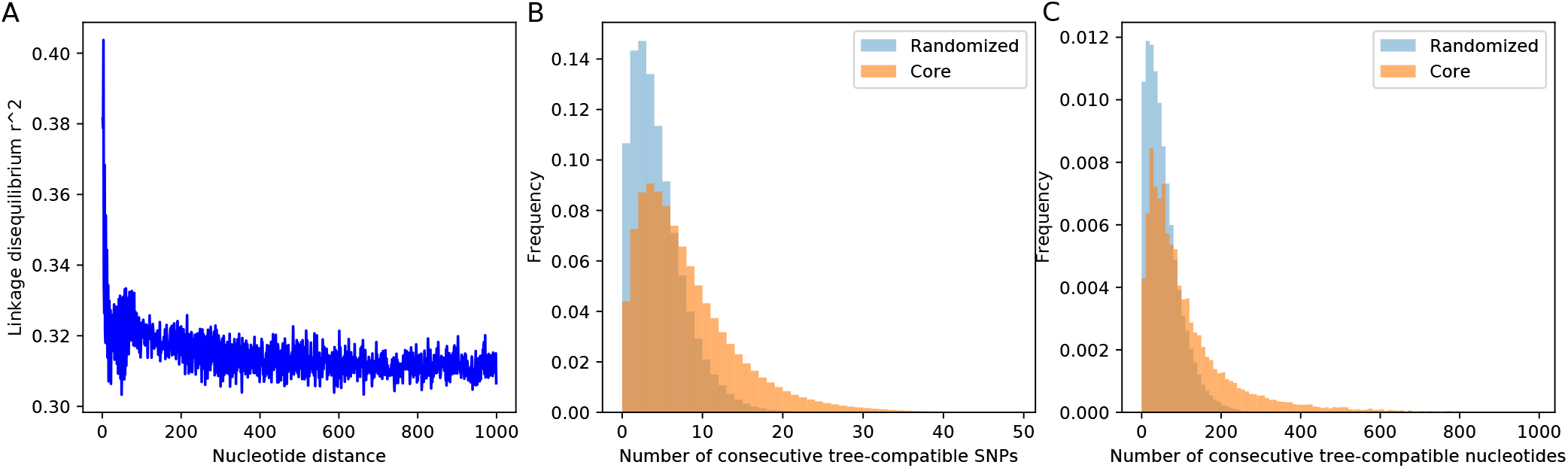
SNP compatibility along the core genome alignment. **A**: Linkage disequilibrium (squared correlation, see Materials and Methods) as a function of the separation of a pair of columns in the core genome alignment. **B**: Probability distribution of the number of consecutive SNP columns that are consistent with a common phylogeny for the core genome alignment (orange) and for an alignment in which the position of all columns has been randomized (blue). **C**: Probability distribution of the number of consecutive alignment columns consistent with a common phylogeny for both the real (orange) and randomized alignment (blue).

Starting from each SNP s, we determined the number of consecutive SNPs *n* that are all mutually consistent with a common phylogeny. As shown in Fig. 5B, the distribution of the lengths of tree-compatible stretches has a mode at *n* = 4, and stretches are very rarely longer than 20 consecutive SNPs. In terms of number of base pairs along the genome, tree-compatible segments are typically just a few tens of base pairs long, and very rarely more than 300 base pairs (Fig. 5C). Thus, stretches of tree-compatible segments are very short. For comparison, we also calculated the distribution of tree-compatible segment lengths in an alignment where the positions of all columns have been completely randomized and observe that these are still a bit shorter (blue distributions in Fig. 5). Thus, while there is some evidence that neighboring SNPs are more likely to be compatible than random pairs of SNPs, this compatibility is lost very quickly, typically within a handful of SNPs.

To elucidate how the recombination-to-mutation ratio determines the lengths of tree-compatible stretches we calculated the same distribution for the data from the simulations with different recombination rates. In addition, to assess the affect of homoplasies on the distribution of the lengths of tree-compatible stretches, we also calculated these distributions for alignments from which 5% or 10% of potentially homoplasic sites were removed (Suppl. Fig. S12). In addition, Suppl. table S1 shows the average length of tree-compatible segments for each of these datasets. We see that removal of homoplasies only has a significant effect on the distribution of tree-compatible stretches for *ρ/μ* ≤ 0.01. Since the fraction of SNPs that are homoplasies is 2.5%, this shows that homoplasies only significantly affect the distribution when *ρ/μ* is less than the rate of homoplasies. For the *E. coli* data, removal of homoplasies has only a minor effect, i.e. the average number of consecutive tree-compatible SNPs increases from 7.6 to 8.8 when 5% potentially homoplasic sites are removed. Although there is of course no reason to assume that the simple evolutionary dynamics of any of our simulations realistically describes the genome evolution of the *E. coli* strains, we note that the distribution of tree-compatible stretches are between what is observed for simulations with *ρ/μ* = 0.3 and *ρ/μ* =1. As is clear from Suppl. Figs S3, S4, and S8, at *ρ/μ* ≥ 0.3 the evolution along every branch of the phylogeny is almost completely dominated by recombination.

In summary, our analysis shows that, as one moves along the core genome alignment, the phylogeny changes typically every 5 — 10 SNP columns, i.e. every 50 — 100 nucleotides. We next use this to put a lower bound on the ratio of the number of phylogeny changes to mutations in the alignment.

### A lower bound on the ratio of phylogeny changes to substitution events

Every time inconsistent SNP columns are encountered as one moves along the core genome alignment, the local phylogeny must change. For example, somewhere between columns *X* and *Z* in Fig. 3 the phylogeny must change. This in turn implies that the start (or end) of at least one recombination event must occur between columns *X* and *Z*. By going along the core genome, and determining the minimum number of times the phylogeny must change, one can thus derive a lower bound on the total number of recombination events and, in the study of sexually reproducing species, this has been a standard method to put a lower bound on the number of recombination events within a genome alignment [17] (see Materials and Methods). Using this we find that the phylogeny must change at least *C* = 43’575 times along the core phylogeny. However, this neglects that some of the inconsistencies may result from homoplasies. To correct for this we remove 5% of potential homoplasic positions by removing 5% of the SNP columns that are most inconsistent with neighboring columns (Materials and Methods) and find that the phylogeny must still change *C* = 34’030 times along this 5% homoplasy-corrected alignment. Because homoplasies are relatively rare, the number of bi-allelic SNPs in the alignment is a good estimate for the total number of mutations in the alignment and the ratio *C/M* thus provides a lower bound for the ratio between the total number of phylogeny changes and substitutions that occur along genome alignment. For the 5% homoplasy-corrected alignment we obtain a ratio *C/M* = 0. 129

Apart from the full alignment, we can calculate the lower bound on the ratio of phylogeny changes to substitution events *C/M* for any subset of strains. Figure 6 shows the ratio *C/M* for random subsets of our 92 strains as a function of the number of strains in the subset, using again the 5% homoplasy-corrected alignment. For comparison, Suppl. Fig. S13 also shows the same results for the full alignment, which would give to ratios *C/M* that are about 20% higher.

**Figure 6:**
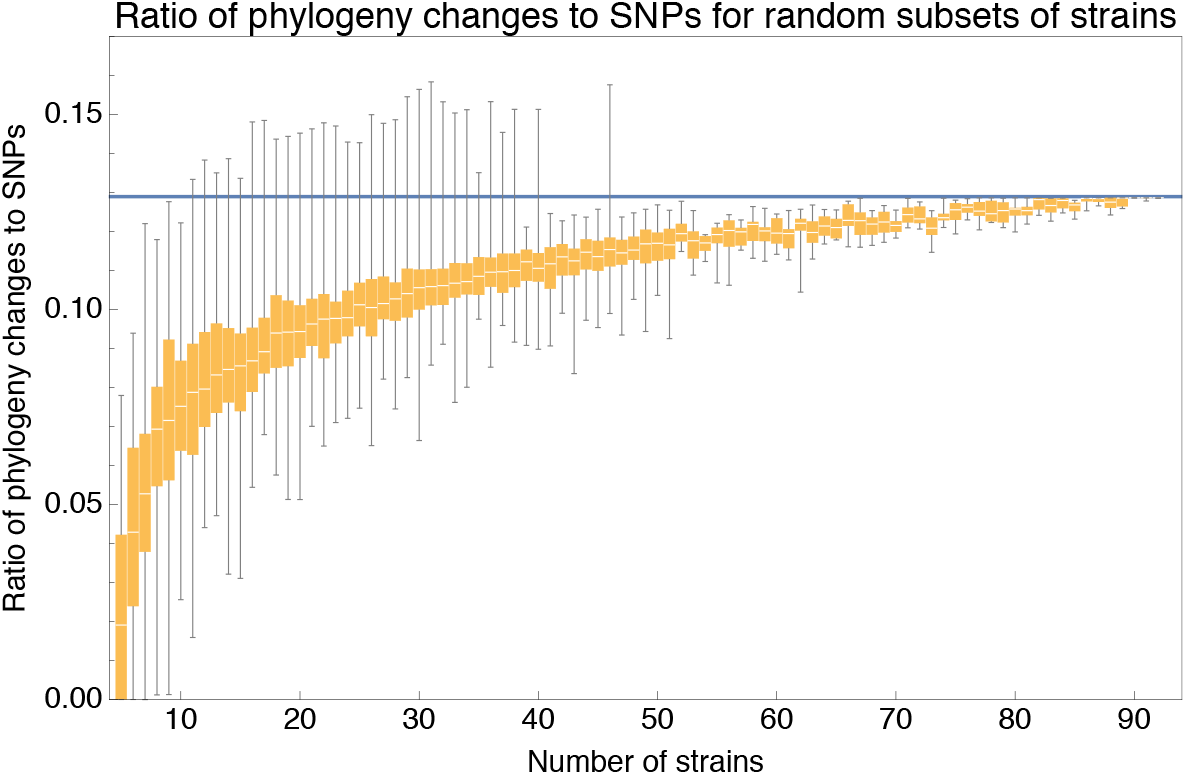
Ratio *C/M* of the minimal number of phylogeny changes C to mutations M for random subsets of strains using the alignment for which 5% of potentially homoplasic positions have been removed. For strain numbers ranging from *n* = 4 to *n* = 92, we collected random subsets of *n* strains and calculated the ratios *C/M* of phylogeny changes to SNPs in the alignment. The figure shows box-whisker plots that indicate, for each strain number *n*, the 5th percentile, first quartile, median, third quartile, and 95th percentile of the distribution of *C/M* across subsets. The blue line shows *C/M* = 0.129.

We see that, for small subsets of strains, the ratio *C/M* shows substantial fluctuations. For example, for subsets of *n* =10 strains, the ratio *C/M* ranges from 0.026 to 0.112, with a median of 0.075. However, as the number of strains in the subset increases, the ratio converges to the value *C/M* = 0.129 and for large subsets of strains there is little variation in the ratio *C/M*.

### *C/M* within phylogroups

Apart from random subsets of strains, we also calculated *C, M* and *C/M* for subsets of strains from the same phylogroup (Suppl. Table S2, and see Suppl. Fig. S1 for the phylogroup annotation). The ratios *C/M* that are observed for these phylogroups increase with the overall divergence within the phylogroup. For phylogroups that are more than 1% diverged (B1, B2, and D), the ratio *C/M* is close to that of the full alignment, there are at least thousands of phylogeny changes along their sub-alignments. The two phylogroups with lower divergence, i.e. A with divergence 0. 0024 and the outgroup with divergence 0. 003 have lower ratios *C/M*. The ratio is particularly low for the outgroup O for which only 2 phylogeny changes are detected. While this may suggest that recombination rates are particularly low within lineages of the outgroup, it should be noted that such low values of *C/M*, while rare, were also observed for some random subsets of strains of the same size (Fig. 6).

### *C/M* for the simulation data

Note that, because each recombination event introduces at most two phylogeny changes (one at the start of the recombined segment and one at its end), *C/*2 is a lower bound on the number of recombination events that occurred in the evolutionary history of an alignment. However, this lower bound may significantly underestimate the true number of recombination events. To obtain more insight into the relationship between this lower bound and recombination rates we compared the ratios *C/M* of each simulated dataset (using the 5% homoplasy-corrected alignment) with the ratio *ρ/μ* of recombination and mutation rate used in the simulation (Suppl. Fig. S14). The results show that when recombination rates are very low, i.e *ρ/μ* ≤ 0.01 the ratio *C/M* is almost exactly equal to 2*ρ/μ*. In this regime recombination events are so sparse on the alignment that many SNP columns occur between every two consecutive phylogeny changes, and this causes almost every phylogeny change to introduce an inconsistency. Since each recombination event introduces two breaks, *C/M* equals twice the number of recombinations per mutation *ρ/μ*. However, as *ρ/μ* increases, fewer SNP columns occur between consecutive phylogeny changes, and more and more of the phylogeny changes go undetected because they do not introduce inconsistencies between the SNPs. Consequently, the ratio *C/M* becomes systematically lower than *ρ/μ* and the difference can become very large. For example, at *ρ/μ* = 1, the observed ratio *C/M* is almost tenfold lower than *ρ/μ*. Note also that since the lower bound *C/M* cannot exceed 1 (i.e. a phylogeny change at every SNP), the ratio *ρ/μ* can exceed *C/M* by arbitrarily large factors at high *ρ/μ*.

### Estimating the number of times each genomic position has been overwritten by recombination

We can also provide a rough estimate for the typical number of times *T* that each position in the genome has been overwritten by recombination in its history. If *L* is the total genome length, *L_r_* the average length of recombination segments, and *C*/2 a lower bound on the number of recombination events in the alignment, then a lower bound on the average number of times positions in the genome have been overwritten by recombination is *T* = *L_r_C*/(2*L*). For the *E. coli* data with C = 34’030 phylogeny changes for the 5% homoplasy-corrected alignment of length *L* = 2’756’541, and the value of *L_r_* ≈ 20’000 base pairs that we estimated previously (Fig. 2J), we obtain *T* ≈ 125. That is, on average each position in the genome has been overwritten at least 125 times by recombination.

In the simulations we specifically tracked the number of times each position in the alignment was overwritten by recombination along each branch of the clonal phylogeny, and we thus know precisely how many times each position of the alignment was overwritten by recombination in its evolutionary history. Supplementary Figure S15 shows the distribution of the number of times each position in the alignment was overwritten for the simulations with *ρ/μ* ranging from 0.001 to 10. We see that, in line with all previous analyses, positions that are clonally inherited along the whole tree only occur for the lowest recombination rate of *ρ/μ* = 0.001, at *ρ/μ* = 0.01 each position has been overwritten 5 — 25 times, and at *ρ/μ* = 0.1 each position has already been overwritten 120 — 200 times. For each *ρ/μ* we calculated the average number of times *T_true_* that positions were overwritten by recombination and compared this with the estimated number of times using the simple estimate *T_est_* = *L_r_C*/(2*L*). Supplementary Figure S16 shows that, while the estimate is in fact quite accurate at very low mutation rate *ρ/μ* = 0.001, as the recombination rate increases the estimate severely underestimates the true number of times positions in the alignment were overwritten by recombination, e.g. at *ρ/μ* = 1 the true number *T_true_* = 1561 is more than twenty times as large as the estimated number *T_est_* = 67. Thus, for the *E. coli* data it is certainly not implausible that each position in the alignment was in fact overwritten by recombination more than 1000 times.

The observed ratio *C/M* for the *E. coli* alignments is very similar to the value of *C/M* observed in the simulations with *ρ/μ* = 1 (Suppl. Fig. S14). Although it is tempting to conclude from this that the ratio of recombination to mutation rate must be close to 1 for the *E. coli* strains, such a conclusion would be unwarranted. The evolutionary dynamics of the simulations makes several strong simplifying assumptions, i.e. that the clonal phylogeny is drawn from the Kingman coalescent process, and that the population is completely mixed so that all strains are equally likely to recombine. Both these assumptions may not apply to the evolution of *E. coli* in the wild. Indeed, we will see below that there is strong evidence that relative recombination rates vary highly across lineages so that instead of a single recombination rate, there is a wide distribution of recombination rates between different lineages. Therefore, it could be misleading to describe *E. coli*’s evolution by a single recombination rate *ρ* and we instead focus on providing a lower bound *C/M* on the number of phylogeny changes per SNP column, which is a meaningful quantity independent of the precise evolutionary dynamics that caused the substitutions and phylogeny changes, and can be calculated independently for any subset of strains.

### Other bacterial special show similar patterns of recombination-dominated genome evolution

To investigate to what extent the observations we made for *E. coli* generalize to other species of bacteria, we selected 5 additional species from different bacterial groups for which sufficiently many complete genome sequences of strains were available, and used Realphy to obtain a core genome alignment of the strains for each species (see Table S3 for a list of the species, the number of strains, and other core genome statistics for each species). We then performed most of the analyses that we presented above for *E. coli* on each of these core alignments. Figure 7 presents a summary of the results that we observe across the species.

**Figure 7.**
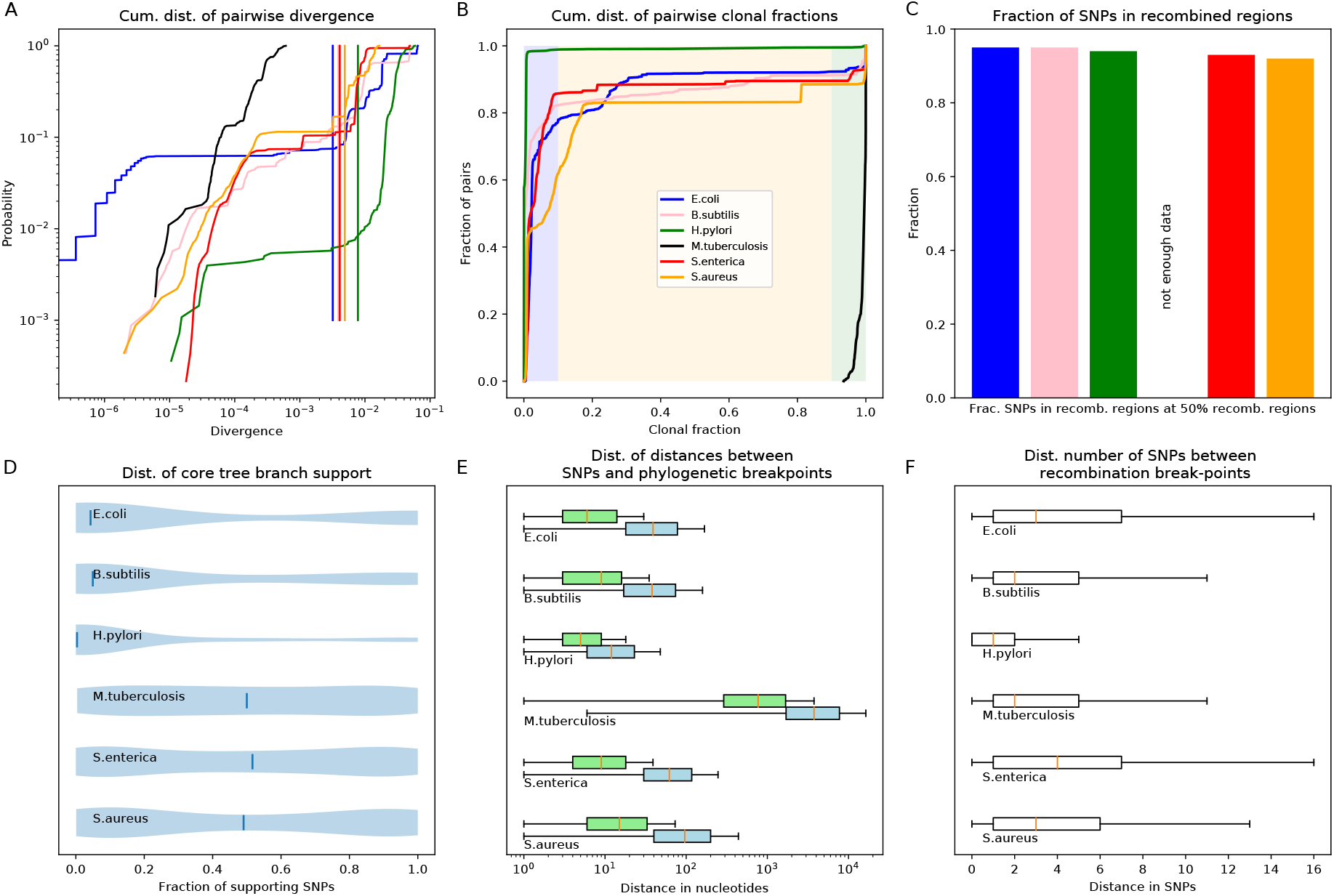
Quantification of the importance of recombination across species. **A**: The cumulative distribution of pairwise divergences is shown as a different colored line for each species (see legend in panel B). Both axes are shown on logarithmic scales. The vertical lines in corresponding colors show the critical divergences at which half of the genome is recombined for each species. **B**: Cumulative distributions of clonal fractions across the pairs of strains for each species, with the blue, yellow, and green shaded regions indicating the fully recombined, partially recombined, and mostly clonal regimes, respectively, i.e. analogous to Fig. 2I **C**: For each species, the height of the bar shows the fraction of SNPs that fall in recombined regions for pairs of strains for which half of the genome is recombined, i.e. see Fig. 2H. **D**: The violin plots show, for each species, the distribution of branch support, i.e. the relative ratio of SNPs supporting versus clashing with each branch split, analogous to the right panel of Fig. 4. The blue lines correspond to the medians of the distributions. **E**: Box-whisker plots showing the 5, 25, 50, 75, and 95 percentiles of the distributions of nucleotide distances between consecutive SNPs (green) and phylogeny breakpoints (blue, i.e. analogous to Fig. 5C), for each species. The axis is shown on a logarithmic scale. **F**: Box-whisker plots of the distributions of the number of consecutive SNPs in tree-compatible segments, i.e. analogous to Fig. 5B.

Figure 7A shows the cumulative distributions of pairwise divergences between strains for all species. We see that, while among our *E. coli* strains that were sampled from a common habitat there is a small percentage of very close pairs with divergence below 10^-6^, for the strains of the other species the closest pairs are at divergence 10^-5^. With the exception of *M. tuberculosis*, where the median pair divergence is around 10^-4^, the median pairwise divergence in all other species is around 10^-2^ or larger. The vertical lines in Fig. 7A indicate the critical divergences, for each species, where half of the alignment is recombined. With the exception of *M. tuberculosis*, where all pairs are mostly clonal, the critical divergences lie in a fairly narrow range of 0.003 — 0.01. Figure 7B shows the cumulative distributions, across pairs of strains, of the fraction of the alignment that is clonally inherited, i.e. as for Fig. 2I for *E. coli*. Note that, for all species except *M. tuberculosis*, the large majority of the pairs is fully recombined. For *H. pylori* the fraction of pairs that still contain clonally inherited DNA is almost zero, whereas for *S. aureus* the fraction of pairs with a substantial fraction of clonally inherited DNA is largest, i.e about 15% with more than 20% clonally inherited DNA. Thus, we see that for almost all species the situation is similar to what we observed in *E. coli*: for most pairs the distance to their common ancestor cannot be estimated from their alignment, because the entire alignment has been overwritten by recombination events. Note also that, for all species, there is only a relatively small fraction of pairs that lie in the partially recombined regime (yellow segment in Fig. 7B).

For *E. coli* we found that, even for close pairs for which a substantial fraction of the genome was clonally inherited, most of the SNPs between them still derive from recombination (Fig. 2H). We find that, with the exception of *M. tuberculosis*, this applies to the other species as well. Figure 7C shows, for each species, the fraction of all SNPs that derive from recombination, for pairs of strains that are at the critical divergence where half of the alignment is recombined. Even though this critical divergence occurs for pairs that are relatively close compared to the typical distance between pairs, for all species more than 90% of the SNPs derive from recombination. That is, we also see that for all 5 species the divergence between close strains is dominated by SNPs that are introduced through recombination.

Another way to quantify to what extent the observed genomic variation can be explained by evolution along the branches of the core genome phylogeny, is to calculate what fraction SNPs supports versus reject each branch of the core tree. Figure 7D summarizes the distributions of support of the branches of the core tree as violin plots, i.e. as shown for *E. coli* as a cumulative distribution in Fig. 4. In *E. coli* most branches have many more SNPs that reject the split than support it, and even stronger rejection of the branches of the core tree are observed for *B. subtilis* and *H. pylori.* For the other three species, including *M. tuberculosis*, an almost uniform distribution of branch support is shown, i.e. for these species there are roughly as many branches that are strongly supported by the SNPs, strongly rejected by the SNPs, or supported and rejected by roughly equally many SNPs.

Figure 7E summarizes, for each species, the distribution of distances between SNPs along the core alignment as box-whisker plots (green) as well as the distribution of distances between phylogeny breakpoints (blue), i.e. as shown in Fig. 5C for *E. coli*. The figure shows that, with the exception of *M. tuberculosis*, the inter-SNP distances range from a few to a few dozen base pairs, with a median inter-SNP distance of 4 *(H. pylori)* to 15 (S. *aureus)* base pairs. For these 5 species, the median distances between phylogeny breakpoints range from around 10 (H. *pylori)* to about 100 base pairs for *S. aureus.* Note that, for all species, the tail of the distributions stretches to very short distances between breakpoints, whereas distances between breakpoints of more than 200 bps are very rare for all these 5 species. Thus, for these species the segments that are consistent with a single phylogeny are always much shorter than the typical length of a gene. In contrast, for *M. tuberculosis* both the distances between SNPs and the distances between breakpoints are almost two orders of magnitude larger, indicating that these strains are much more closely related than the strains of the other species.

Figure 7F shows box-whisker plots for the distributions of the number of consecutive SNPs between breakpoints, as was shown for *E. coli* in Fig. 5B. We see that for all species, including *M. tuberculosis*, there are typically less than a handful of SNPs in a row before a phylogeny breakpoint occurs, and very rarely more than a dozen SNPs. The smallest number of SNPs per breakpoint is observed for *H. pylori*, i.e. typically less than 2 SNPs per breakpoint, but the range of SNPs per breakpoint is very similar across all species. Finally, Suppl. Table S4 shows the fraction of alignment columns that are SNPs, the lower bound *C* on the number of phylogeny changes, and the lower bound *C/M* on the ratio of phylogeny changes to SNP columns, for each of the 6 species. We see that with the exception of H. pylori, which has a high ratio *C/M* ≈ 0.3, and M. tuberculosis which has significantly lower ratio *C/M* ≈ 0.08, the other species have ratios *C/M* similar to that observed for *E. coli*.

In summary, with the exception of *M. tuberculosis* for which all strains are quite close and most DNA has been clonally inherited for most pairs, all other species show the same pattern as *E. coli* with genome evolution being dominated by recombination. For most pairs no DNA in their alignment was clonally inherited, even for close pairs most of the SNPs derive from recombination events, and the phylogeny changes thousands of times along the core genome alignment, typically within a handful of SNPs.

### Phylogenetic structures reflect the distribution of relative recombination rates across lineages

All our results so far show that the core tree does not capture the evolutionary relationships between the strains. In fact, rather than a single phylogeny representing the evolutionary relationships between the strains, the SNP data show that the phylogeny changes thousands of times along the core genome alignment, suggesting a large number of different phylogenies occurs across the alignment. It may thus seem all the more puzzling that, when trees are constructed from sufficiently many genomic loci, the core tree reliably emerges (Fig. 1, bottom).

As we mentioned in the introduction, some researchers interpret this convergence to the core tree to mean that the core tree must correspond to the *clonal* phylogeny of the strains. The interpretation is that the SNP patterns in the data are a combination ‘clonal SNPs’ that fall on the clonal phylogeny, plus a substantial number of ‘recombined SNPs’ that were affected by recombination, which act so as to introduce noise on the clonal phylogenetic signal. In this interpretation, trees build from individual loci can differ from the core tree because the ‘recombination noise’ can locally drown out the true clonal signal, but once sufficiently many loci are considered, the recombination noise ‘averages out’ and the true clonal structure emerges. However, this interpretation makes two assumptions that do not necessarily hold. First, assuming that the effects of recombination will ‘average out’ when sufficiently many loci are considered only holds if recombination is effectively random and does not have clear population structure itself. Second, for the clonal phylogeny to emerge one has to assume that there are sufficiently many genomic loci left that have not been affected by recombination, whereas our analyses above have shown that there is no clonally inherited DNA left for most pairs, and that each locus in the genome has been overwritten many times by recombination.

Indeed, if we remove all SNP columns from the core genome alignment that correspond to branches of the core tree, and then reconstruct a phylogeny from this edited alignment, the resulting tree is still highly similar to the core tree (Suppl. Fig. S17). That is, we obtain a very similar tree from this edited alignment, even though virtually *all* SNPs in this alignment are inconsistent with the tree. This confirms that the structure in the core tree does not derive from the subset of SNPs that fall on the core tree, but rather suggests that it must reflect overall statistical properties of the SNP patterns that result from structure in the recombination patterns.

We propose that, to understand the phylogenetic structure evident in the data, we should not think of the SNP patterns deriving from a single ‘true’ phylogeny plus noise generated by recombination, but rather accept that many different phylogenies occur along the alignment, and that the core tree reflects statistical properties of the *distribution* of phylogenies. To explain this, we will use data from a species for which there is no question that recombination dominates and no ‘clonal’ structure can exist.

### Phylogenies reconstructed from human genome sequences also exhibit robust phylogenetic structure

We randomly selected 40 genomes from the 1000 Genomes project, used PhyML to build a phylogenetic tree from chromosomes 1 — 12 of these genomes, and then investigated to what extent branches in this tree also occur in trees build from random subsets of the genomic loci. As shown in Suppl. Fig. S18, a robust phylogeny also emerges for the human data. In particular, individuals with the same geographic ancestry consistently form clades in the trees build from large numbers of loci.

However, it is of course clear that this phylogeny cannot correspond to the ‘clonal phylogeny’ of the human sequences, because there is no such thing as a ‘clonal phylogeny’ for the human sequences. Perhaps the closest analog of a ‘clonal tree’ for the human data would be the phylogeny of the strictly maternal lineage. However, since at each generation roughly half of the autosomal chromosomes derives from the mother, and half from the father, there are few if any loci in the genome that follow this strictly maternal lineage, i.e. almost every locus was paternally inherited in at least *some* generations. Instead, it is clear that there are many different phylogenies along the genome, each with different ancestries.

The reason that a robust phylogeny emerges is that the different phylogenies that occur along the genome are not completely random. That is, human populations are not completely mixed but people are more likely to mate with others from the same geographic area. Because of this, recombination tends to occur among people of the same geographic area, and because of this population structure the phylogenies that occur along the alignment of a set of human genomes are not sampled uniformly from all possible phylogenies, but some topologies are more likely to occur than others. In particular, individuals from the same geographic area will have recent ancestors in a larger fraction of the phylogenies along the genome than individuals from different geographic areas. Indeed, a simple principal component analysis of SNP statistics in genomes of European ancestry recapitulates geographic structure in remarkable detail [31].

Of course, there are good reasons that human population geneticists don’t typically represent the ancestral population structure of human sequences by a single phylogeny. We know that many different phylogenies occur along the genome and it would be misleading to pretend that this distribution of phylogenies can be summarized by a single tree. However, if one insists on representing the population structure by a single tree and asks PhyML to build a single phylogeny from many loci of the human sequences, then one indeed does get convergence to a common phylogeny for the human data as well. The reason one gets this convergence is because asking for a single tree is analogous to asking for the ‘average’ of a distribution. The average of almost any distribution becomes very robust given enough samples, even if this average does not actually represent any typical sample. For example, instead of the total divergence between a pair of individuals representing the distance to their clonal ancestor, the total divergence represents the *average* distance to the many different common ancestors of the pair across the many different phylogenies along the genome, and this average is a function of the rates at which their lineages have recombined. That is, the distances in this tree reflect the relative rates of recombination among the different lineages.

We propose that the situation for the bacterial data is very similar to the situation for the human data. Our analysis above has shown that many different phylogenies occur along the alignment and that the core tree must reflect the statistics of this distribution of phylogenies. We propose that, in the same way as for the human data, the core tree that results from applying PhyML to the core genome of a set of bacterial strains reflects the relative rates with which different lineages have recombined. To support this, we return to the *E. coli* data and use the SNP statistics to quantify the relative rates with which different lineages have recombined.

### Recombination rates across lineages follow approximately scale-free distributions

The analyses above have shown that the core alignment of the *E. coli* strains consists of tens of thousands of short segments with different phylogenies. Thus, one approximate way of thinking about the core genome alignment is that the phylogeny at every genomic locus is drawn from some distribution of possible phylogenies. In a freely recombining population, every strain would be equally likely to recombine with any other, leading to a uniform distribution over all possible phylogenies. Consequently, for any pair of strains, the average distance to their common ancestors across the phylogenies would become roughly the same, so that all pairs would become approximately equidistant. Phylogenies built from large numbers of genomic loci would than take a star-shape, which is clearly at odds with our observations. This suggests that the phylogenies along the genome are drawn from a highly non-uniform distribution in which some lineages are much more likely to have recombined recently than others.

The distribution of observed SNP types in fact contains extensive information about the relative frequencies with which different lineages have recombined at different times in the past. For example, imagine a SNP where two strains share a nucleotide which differs from the nucleotide that all other strains possess. We will denote such SNPs as 2-SNPs or pair-SNPs. If, at some genomic locus *g*, we find a 2-SNP shared by strains si and *s*_2_, then it follows that, whatever the phylogeny is at locus *g*, the strains *s*_1_ and *s*_2_ must be nearest neighbors in the tree, and the SNP corresponds to a mutation that occurred on the branch connecting the ancestor of *s*_1_ and *s*_2_ to all other strains. Formally, the number of 2-SNPs (*s*_1_, *s*_2_) is proportional to the fraction of all genomic loci where *s*_1_ and *s*_2_ are nearest neighbors in the phylogeny, times the average length of the branch from the ancestor of *s*_1_ and *s*_2_ to the next common ancestor with other strains.

Thus, to quantify the extent to which the lineages of a strain s have recently recombined with the lineages of the other strains, we can extract all 2-SNPs in which *s* shares a letter with one other strain *s*’ and compare their frequencies. As an example, Fig. 8A graphically shows the frequencies of all pair-SNPs (*A*1, *s*) in which A1 shares a SNP with one other strain *s*. Note that, if there was a dominant clonal phylogeny, then A1 should have 2-SNPs with its nearest neighbor in this dominant phylogeny much more often than with any other strain. If, on the other hand, all lineages were freely recombining at the same rate, then we would expect roughly equal frequencies of all possible 2-SNPs (*A*1, *s*). However, we observe neither of these patterns. We see that A1 shares 2-SNPs with 17 of the 92 strains in our collection at a wide range of frequencies. Instead of a clear ‘clonal partner’ with most 2-SNPS, A1 shares 2-SNPs almost equally frequently with 3 strains, i.e. A2, A11, and D8 each share about 200 2-SNPs each with A1. In contrast, for 11 of the 17 strains the number of occurrences is 10 or less, and for 4 strains a 2-SNP with A1 is observed only once.

**Figure 8.**
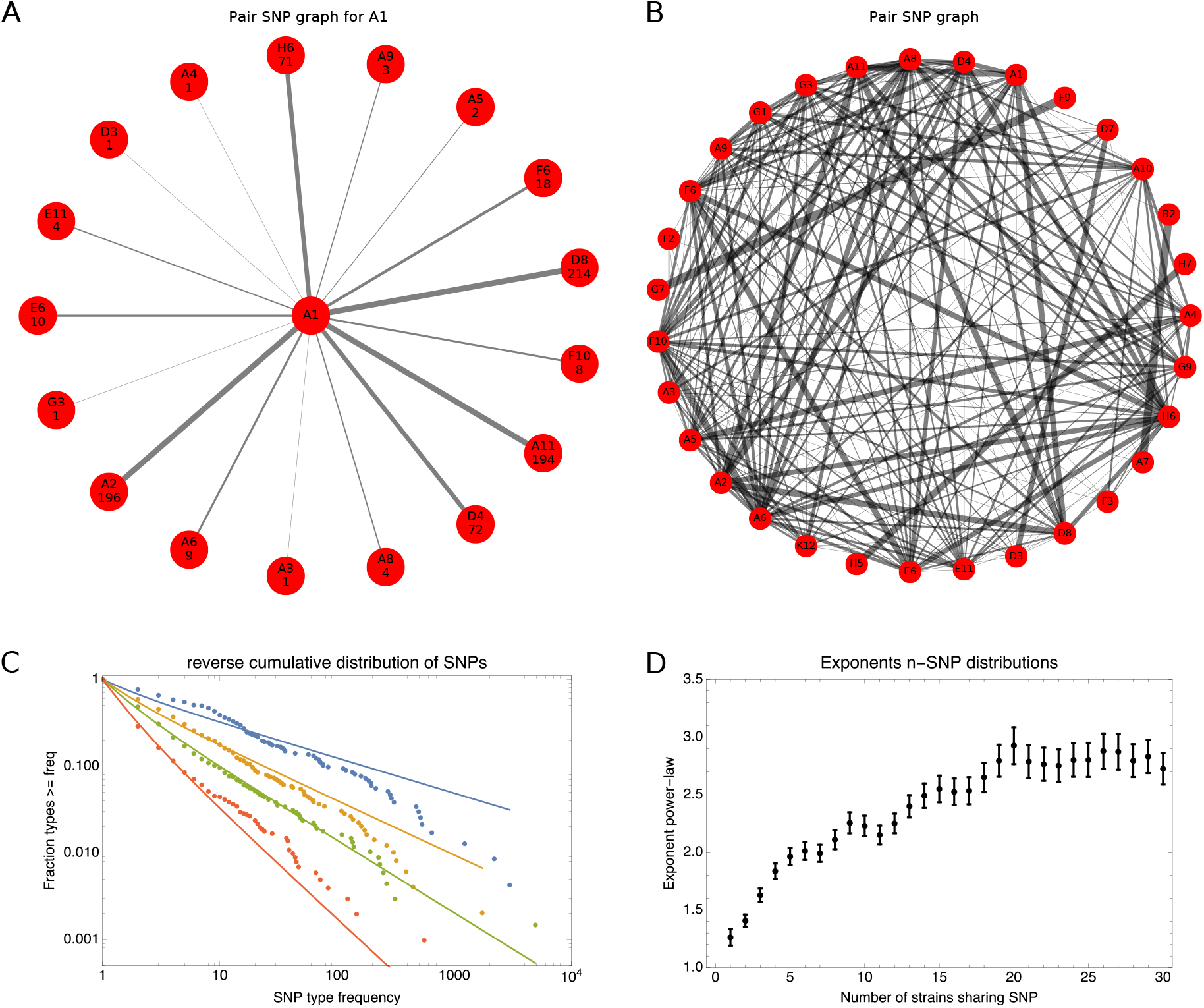
SNP-type frequencies follow approximately power-law distributions. **A**: Frequencies of 2-SNPs of the type (*A*1, *s*) in which a SNP is shared between strain A1 and one other strain *s*. Each edge corresponds to a 2-SNP (*A*1, *s*) and the thickness of the edge is proportional to the logarithm of the number of occurrences of the 2-SNP. The frequency of each edge is also indicated at the corresponding outer node. **B**: A graph showing all 2-SNPS (*s, s*’) that were observed in the core genome alignment. Each node corresponds to a strain and each edge to a 2-SNP, with the thickness of the edge proportional to the logarithm of the number of occurrences of the SNP. **C**: Reverse cumulative distributions of the frequencies of all observed 2-SNPs (blue dots), 3-SNPs (orange dots), 4-SNPs (green dots), and 12-SNPs (red dots). The solid lines in corresponding colors show power-law fits. Both axes are shown on a logarithmic scale. **D**: Exponents of the power-law fits to the *n*-SNP frequency distributions, as a function of the number of strains sharing a SNP *n*. Error bars correspond to 95% posterior probability intervals.

The relative frequencies of different pair-SNPs (*s*_1_, *s*_2_) reflect the relative rates with which the lineages of different pairs of strains have recently recombined. Figure 8B shows a graph representation of all observed pair-SNPs, with the thickness of the edges proportional to the logarithm of the frequency of occurrence of each 2-SNP type. We see that each strain is connected through 2-SNPs to a substantial number of other strains, indicating a high diversity of recent recombination events across the strains. At the same time, the large variability in the thickness of the edges indicates that some pairs occur much more frequently than others. Figure 8C shows the reverse cumulative distribution of the frequencies of all observed 2-SNPs (blue dots), i.e. the distribution of the thickness of the edges in Fig. 8B. Note that, if the strains were to recombine freely, each 2-SNP would be equally likely to occur, and the distribution of 2-SNP frequencies would be peaked around a typical number of occurrences per type. Instead, we see that 2-SNP frequencies *f* vary over more than 3 orders of magnitude, i.e. from an occurrence of just *f* = 1 for many 2-SNPs to *f* = 2965 occurrences for the most common 2-SNP. Moreover, the reverse cumulative distribution of 2-SNP frequencies follows an approximate straight-line in a log-log plot. In other words, the distribution of frequencies *P*(*f*) roughly follows a power-law distribution, i.e *P*(*f*) ∝ *f*^-α^. Fitting the 2-SNP data to a power-law (see Materials and Methods) we find that the exponent equals approximately *α* ≈ 1.41 (blue line in Fig. 8C).

Note that, while the distribution of 2-SNP frequencies is clearly not a perfect power-law, the distribution is long tailed and is much better fit by a straight line in a log-log plot than by a straight line in either a linear or semi-log plot. Importantly, this means that there is no typical frequency of 2-SNPs, and that one cannot naturally divide 2-SNPs into common and rare types. Instead, the distribution of 2-SNP frequencies is approximately scale-free.

Beyond SNPs shared by pairs of strains, we can of course also look at SNPs shared by triplets, quartets, and so on. Besides the distribution of 2-SNP frequencies, Fig. 8C also shows the reverse cumulative distributions of 3-SNPs (orange dots), 4-SNPs (green dots), and 12-SNPs (red dots). We see that all these distributions follow approximately straight lines in a log-log plot and can thus be approximated with power-law fits (solid lines). The *n*-SNP distributions drop more steeply as *n* increases. Figure 8D shows the exponent *α* of the *n*-SNP distribution as a function of *n*, showing that the exponents range from *α* ≈ 1.25 for singlets, i.e. *n* = 1, to *α* ≈ 2.8 for *n* ≥ 20.

We find that essentially all *n*-SNP distributions are approximately scale-free, i.e. can be fitted with power-laws. Thus, while some subgroups of *n* strains share a common ancestor much more often than others subgroups of *n* strains, their frequencies fall along a scale-free continuum, so that there is no natural way of dividing the clades of *n* strains into ‘common’ and ‘rare’ clades. Note also that each *n*-SNP corresponds to a mutation that occurred on the branch leading to the ancestor of a group of *n* strains. Therefore, *n*-SNPs for larger *n* typically correspond to mutational events that occurred further back in time. The fact that *n*-SNP distributions become more steep as *n* increases means that the average number of occurrences per *n*-SNP decreases as *n* increases. Thus, the diversity of *n*-SNPs tends to be larger further back in time (see also Suppl. Text S1.1).

One might wonder to what extent ‘clonal’ SNPs contribute to the observed distributions of *n*-SNPs, e.g. whether the most frequent *n*-SNPs in the tails of the distributions correspond to clonal SNPs. To investigate the effect of potentially clonal SNPs on the *n*-SNP distributions we removed all *n*-SNPS that fall on the branches of the core tree. As shown in Suppl. Fig. S19, the *n*-SNP distributions look virtually identical after all *n*-SNPs corresponding to branches in the core tree have been removed. In addition, the *n*-SNP distributions for the 5% homoplasy-corrected alignments also look virtually identical (Suppl. Fig. S19), showing that homoplasies do not significantly affect the *n*-SNP distributions either.

One might also wonder to what extent these *n*-SNP statistics are different from what would be observed for populations evolving completely clonally, or for completely mixed populations that evolve under mutation and recombination. To investigate this, Supplementary Text S1.1 extensively compares the *n*-SNP statistics observed for *E. coli* with those observed for simulations with different amounts of recombination. This comparison shows that none of the simulation data shows *n*-SNP statistics that mimic what is observed for *E. coli*. At low recombination rate the *n*-SNPs are dominated by clonal SNPs and have low diversity, whereas at higher recombination rates different *n*-SNPs all have similar frequencies, and no long-tailed distributions are observed. These observations further support that *E. coli’s n*-SNP statistics reflect population structure.

Although it is tempting to interpret the *n*-SNP distributions as reflecting an almost ‘scale free’ population structure, they depend in a complex manner on both the distributions of topologies and branch lengths of the phylogenies across the alignment. As we do not yet have a general theory for the evolutionary process that generates the distribution of phylogenies along the alignment, it is at this point difficult to disentangle the contributions from variations in topologies and branch lengths to the *n*-SNP distributions. Consequently, it is difficult to give a precise interpretation of either the approximate power-law form or the meaning of the exponents.

### Phylogenetic entropy profiles of individual strains

Another way to characterize the structure of the *n*-SNP distributions is to quantify, for each strain *s*, how diverse the clades are that *s* occurs in at different *n*. To illustrate the idea we refer back to Fig. 8A, which shows all the 2-SNPs in which strain A1 occurs. Note that these 2-SNPs are all mutually inconsistent, i.e. in any one phylogenetic tree the strain A1 can only occur in a pair with *one* other strain. Therefore, the relative frequencies of the different 2-SNPs in which A1 occurs reflect the relative frequencies of phylogenies in which A1 is paired with different strains, and the diversity of these pairs can be quantified by the *entropy* of this distribution. That is, if *n_s_,A*_1_ is the number of 2-SNPs of type (*s, A*1), then the fraction of pairs for which A1 occurs with *s* is 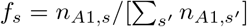, and the entropy of the distribution of 2-SNPs for A1 is 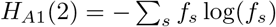.

Note that the same calculation can be done for any strain *s* and any *n*. For example, if the strain *s* occurs in 10 different quartets of strains *q*, then all quartets *q* are mutually inconsistent, and the diversity of quartets in which *s* occurs can be calculated as the entropy *H*_4_(*s*) = – ∑_*q*_ *f_q_* log(*f_q_*) of the relative frequencies with which the different quartets *q* occur. In this way, for each strain *s* we can calculate an entropy profile *H_s_*(*n*) that contains the entropies of the *n*-SNP distributions in which strain *s* occurs, as a function of *n* (see Materials and Methods). Figure 9 shows the entropy profiles of all strains (right panel), as well for a selection of six example strains (left panel), as calculated on the 5% homoplasy-corrected alignments.

**Figure 9.**
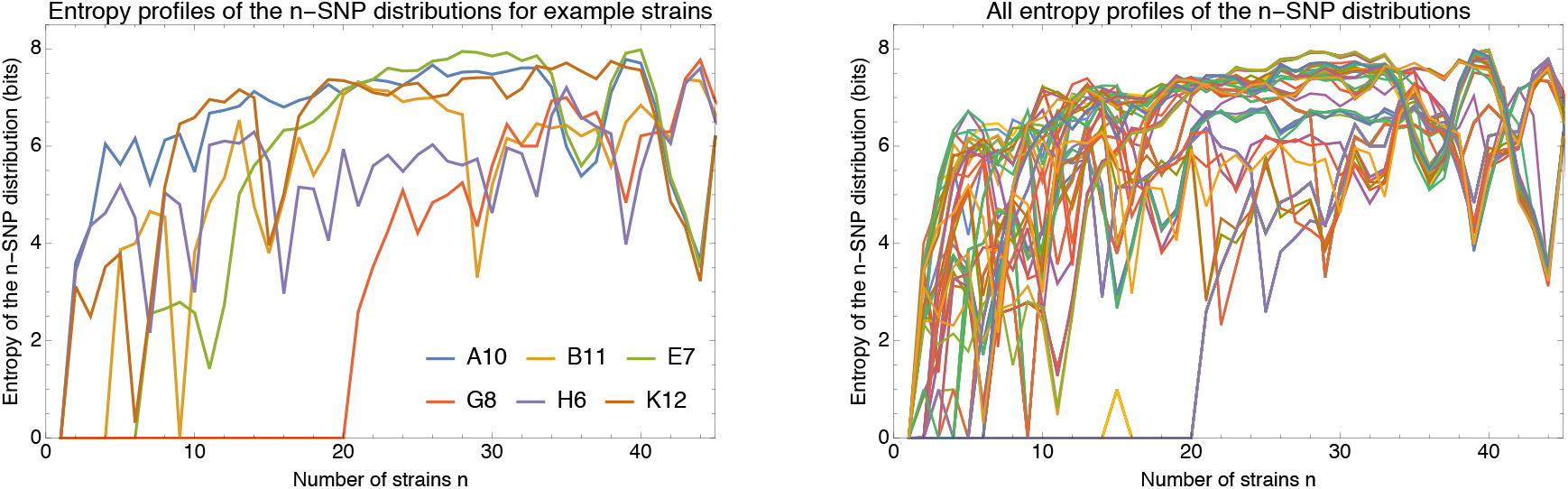
Phylogenetic entropy profiles of the *E. coli* strains. **Left panel**: Entropy profiles *H_s_*(*n*) (in bits) for six example strains, indicated in the legend. **Right panel**: Entropy profiles *Hs*(*n*) for all *E. coli* strains

Note first that, if evolution were strictly clonal, then all entropies *H_n_*(*s*) would be zero but we see entropies going up to 6 — 8 bits for all strains. However, probably the most striking feature of the entropy profiles is their great variability across the strains. If all lineages were recombining with each other at equal rates, we would expect the diversity of *n*-SNPs to increase in the same way for each strain, but the data shows almost as many distinct entropy profiles as there are strains.

For most strains the entropy increases quickly with *n*, e.g. for strain A10 the entropy rises to *H*_4_(*A*10) ≈ 6 bits, which is equivalent to strain A10 occurring in approximately 64 different quartets with equal frequency. In contrast, for strain E7 the entropy stays zero until *n* > 6, and for strain G8 even until *n* > 20. As can be seen in the core tree (Fig. 1), E7 and G8 are parts of groups of *n* =6 and *n* = 20 very closely-related strains, respectively. Both these groups had a clonal ancestor so recently that essentially no recombination events occurred since. Consequently, the strains in these groups occur in at most 1 type of *n*-SNP for *n* ≤ 6 and *n* ≤ 20, respectively. Thus, for groups of *m* strains with a very recent common ancestor, the entropies *H_n_*(*s*) are essentially zero when *n* ≤ *m*.

Although the entropies generally go up with *n*, they do not increase monotonically. For example, for strain B11, the entropy increases to around *H_n_*(*B*11) ≈ 4 for *n* = 5 to *n* =8, but then drops back to almost zero at *n* = 9. Notably, this strain B11 is a member of the outgroup of 9 strains that are highly diverged from all the other strains (Fig. 1) and the fact that *H*_9_(*B*11) = 0 reflects the fact that 9-SNPs shared by all strains of the outgroup are vastly more numerous than any other 9-SNP in which B11 occurs.

Another feature that is evident is that the entropy profiles of some strains appear to nearly merge at large *n*. For example, the entropy profiles of A10 and E7 become very similar from *n* = 34 onward, and from about *n* = 40 on-wards the profile of K12 becomes very similar to these two as well. However, not all entropy profiles merge, and different groups of profiles remain up until *n* = 46. As can be seen in Suppl. Fig. S20, we generally observe that strains from the same phylogroup tend to ‘merge’ their entropy profiles at large *n*. Importantly, however, although the entropy profiles become highly similar at large *n*, they do not become identical. In contrast, for strains with a recent clonal ancestor, the entropy profiles are perfectly identical. For example, the entropy profiles of the close pairs B6 and C1 from phylogroup *F*, and A7 and B2 from phylogroup B2, completely overlap so that only one of the two colors of each pair is visible in the plot (Suppl. Fig. S20). We developed a simple statistical test, based on the Fisher-exact test (Materials and Methods) to quantify the extent to which the *n*-SNP statistics of a pair of strains are significantly distinct. As shown in Suppl. Fig. S21, we find that only very closely related pairs of strains that had a recent common ancestor, corresponding to about 5% of all pairs, have statistically indistinguishable *n*-SNP statistics. Thus, while strains with a recent clonal ancestor have identical entropy profiles, the entropy profiles of strains from a phylogroup have similar but not identical *n*-SNP distributions at large *n*. Moreover, strains from different phylogroups can have very different entropy profiles even at large *n*. To provide additional insights into what the *n*-SNP statistics show about recombination patterns, Suppl. Text S1.2 provides an in-depth analysis of the *n*-SNP statistics for the relatively small phylogroup B2.

Thus, the general picture that emerges from the entropy profiles is that, while some of the structure at small *n* is due to clonal relationships, at large *n* the entropy profiles reflect the statistics of recombination of different lineages. While strains from the same phylogroup tend to show highly similar *n*-SNP statistics at high *n*, suggesting that their lineages have similar recombination statistics sufficiently far into the past, strains from different lineages have clearly different recombination statistics even far into the past.

Finally, the entropy profiles observed for the *E. coli* strains differ markedly from what is observed for the data from simulations (Suppl. Fig. S22). For the simulations without recombination the entropies are almost all zero and at the very low recombination rate of *ρ/μ* = 0.001, where clonal SNPs dominate along all branches of the clonal tree, all entropies are small. In contrast, in the regime where recombination dominates along all branches of the clonal tree, i.e. for *ρ/μ* ≥ 0.3, the entropy profiles of all strains are highly similar. This confirms that when lineages recombine at equal rates, highly similar entropy profiles result. Finally, in the intermediate regime of *ρ/μ* = 0.01 and *ρ/μ* = 0.1 where branches near the leafs are mostly clonal and internal branches are dominated by recombination, we see some moderate variation in the entropy profiles at small *n* due to clonal structure, whereas at high *n* the entropy profiles again merge.

### *n*-SNP statistics and entropy profiles in other species

We next investigated whether the *n*-SNPs of the other species also exhibit approximately power-law distributions, as observed in *E. coli*. Supplementary Figure S23 shows the reverse cumulative distributions of 2-SNPs, 3-SNPs, 4-SNPs, and 12-SNPs across all 6 species together with power-law fits. Although the curves often deviate substantially from simple straight lines, they all exhibit long tails and range over several orders of magnitudes, i.e. up to 5 orders of magnitude for 2-SNPs in *S. enterica.* Note that *M. tuberculosis* is again an exception. The total number of different *n*-SNP types is so small for this species that the *n*-SNP distributions are not well defined. Figure 10 (left panel) shows the fitted exponents of the power-law distributions of *n*-SNPs as a function of *n* for all species. With the exception of *M. tuberculosis*, for which the exponents are small for all *n*, we see that the exponents generally increase with *n* indicating that the phylogenetic diversity generally increases as one moves further back in time, i.e. to larger *n*. Consistent with other observations, *H. pylori* shows the highest exponents, i.e. the highest diversity, and *M. tuberculosis* the lowest. While the exponents become roughly constant for *n* > 20 for *E. coli, H. pylori* and *S. aureus, B. subtilis* and *S. enterica*, exhibit more complex patterns with sudden drops in exponent at particular values of *n*, suggesting more complex population structures for these species.

**Figure 10.**
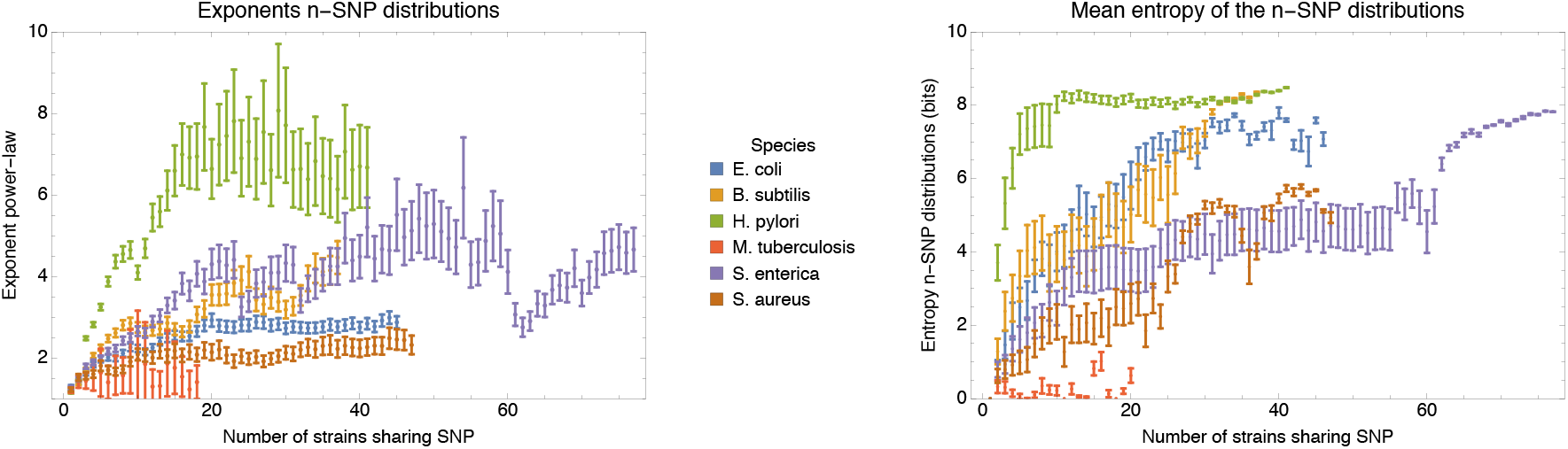
**Left panel**: Exponents of the power-law fits to the *n*-SNP frequency distributions, as a function of the number of strains sharing a SNP *n* for each of the species (different colors). Error bars correspond to 95% posterior probability intervals. **Right panel**: Mean entropy of the entropy profiles *H_n_*(*s*), averaged over all strains *s*, as a function of the number *n* of strains sharing the SNP, for each of the species (different colors). The error bars correspond to two standard-errors of the mean.

Supplementary Figure S24 shows the entropy profiles *H_n_*(*s*) for all strains s in each of the species. As we observed in *E. coli*, different strains have highly variable entropy profiles in other species as well, showing that also in these other species almost every strain has a unique ‘fingerprint’ of frequencies with which its lineage shares ancestors with those of other lineages. Although the entropy rises quickly to values in the range 4 — 8 for most strains, we also see strains for which the entropy only rises after *n* exceeds some fairly large value of *n*, e.g. at *n* = 10 for some strains in *H. pylori*, and at *n* = 24 and *n* = 62 for some *S. enterica* strains, suggesting that these strains are part of groups of very closely-related strains. Note also that these events appear to correspond to the sudden drops in the exponents of the *n*-SNP distributions of those strains (Fig. 10, left panel), reiterating that these *n*-SNP statistics encode information about the population structure of each species. To summarize the entropy profiles of each species, the right panel of Fig. 10 shows the mean and standard-error of the entropy profiles, averaged over all strains, as a function of *n*. As for most other statistics, *M. tuberculosis* is an outlier whose strains generally only show low phylogenetic entropy. For all other species, the average entropy clearly increases as *n* increase, indicating again that the phylogenetic diversity increases further back in the past. For 4 of the 6 species, the mean entropy at large *n* falls in a narrow range between 7 and 8 bits, suggesting that the effective number of different ancestries far back in the past is relatively similar for these species.

Finally, for comparison we decided to also calculate these patterns on the human data. In particular, extracted SNP data for chromosomes 1 — 12 for 2504 humans from the 1000 Genomes project [32]. Supplementary Fig. S25 shows examples of the *n*-SNP distributions for human together with the fitted exponents for *n* ranging from 1 to 30, as well as the entropy profiles for the 40 randomly chosen individuals used previously in Fig. S18. Apart from the 2-SNPs, which appear to show a biphasic pattern, all other *n*-SNP distributions in human are all well fit by power-law distributions. Moreover, for *n* ≥ 5 the exponents are almost constant around a value of 2.8. Interestingly, the entropy profiles of all 40 individuals quickly rise to around 10 bits. However, from that point onward the entropy profiles of individuals with African ancestry deviate from those of the other individuals, showing consistently higher entropy, confirming the well-known fact that African populations have higher genetic diversity. In contrast to what we observe for the bacterial genomes, for the human data the entropy profiles of individuals with the same geographic ancestries all look very similar.

## Discussion

In this work we have introduced new methods to analyze prokaryotic genome evolution from multiple alignments of the core genomes of strains from a species. In particular, showing that almost all bi-allelic SNPs in the core genome alignment correspond to single mutations in the history of that position in the alignment, we showed several new ways in which these SNPs can be used to quantify phylogenetic structures and the role of recombination in genome evolution within prokaryotic species.

Our analysis shows that, for all but one of the species studied here, evolution of the core genome is almost entirely driven by recombination. For example, recombination is so common that for the large majority of pairs of strains, none of the DNA in their pairwise alignment derives from their clonal ancestor. Although pairs of strains exist that are so closely related that most of their DNA was clonally inherited, even for such pairs the large majority of mutations that separate them derive from recombination events. When considering the core genome alignment of a collection of roughly 100 strains, we found that each position has been overwritten many times by recombination, i.e. hundreds if not more than a thousand times for *E. coli*. Given this, there is no reason to assume that a phylogeny reconstructed using maximum likelihood from the core genome alignment corresponds to the clonal phylogeny of the strains. Moreover, methods that aim to reconstruct the clonal phylogeny from a subset of positions that have not been affected by recombination are also inherently problematic because our analysis shows that, for most species, such positions simply do not exist.

Although we cannot completely exclude that sufficient information about the ancestral phylogeny is still encoded in some way into the core alignment, it is clear that currently no method exists that is capable of extracting this information, and we suspect that it is in fact impossible, i.e. that recombination has destroyed the necessary information. However, even if it were possible to reconstruct the ancestral phylogeny, it is not clear how useful this clonal phylogeny would be for understanding core genome evolution. As already mentioned, even for very close pairs most of their divergence results from recombination events. Moreover, our analysis of SNP compatibility along the core alignment shows that the phylogeny changes every few dozen base pairs (and every handful of SNPs), so that the core genome alignment fragments into many thousands of short segments with different phylogenies. Thus, modeling sequence evolution in the core genome as occurring along the branches of a fixed phylogenetic tree is clearly inappropriate.

One might infer from these statistics that bacterial species are quasi-sexual and recombining freely, but as has been noted previously [33], this assumption is also inconsistent with the data. For example, strains do not appear roughly equidistant and phylogenies build from large numbers of genomic loci clearly converge to a well-defined phylogeny. We argued that, instead of the clonal phylogeny, this core genome phylogeny represents the statistics of the *distribution* of phylogenies that occur along the core genome alignment, and that this distribution in turn reflects population structure, i.e. the relative rates with which different lineages have recombined. To support this, we showed how bi-allelic SNPs can also be used to quantify the population structure. In particular, the frequencies of different *n*-SNP types reflect the relative frequencies with which different subsets of *n* strains share a common ancestor across the phylogenies. We find that the frequencies of *n*-SNP types vary over 3 — 5 orders of magnitude and follow roughly power-law distributions. Notably, since the *n*-SNP distributions follow smooth long-tailed distributions that do not appear to have a characteristic scale, it is not possible to naturally subdivide subsets into highly and rarely recombining types. Rather, there seems to be a large continuum of relative rates, with population structure on all scales. Given that recombination rates vary over orders of magnitude across different lineages, the idea of an effective single recombination rate for a species might be misleading, and it seems problematic to fit the data to models that assume a constant rate of recombination within a species [34].

Essentially all population genetics and coalescent models start from assuming one or more populations of individuals that, for the purpose of the model, are exchangeable. However, our analysis of the entropy profiles and *n*-SNP statistics showed that, apart from some groups of extremely closely-related strains that share a common ancestor before any recombination events occurred, almost every strain has its own distinct *n*-SNP statistics. This suggests that almost every strain has a unique pattern of relative recombination rates between its lineages and the lineages of the other strains, so that models that start from populations of exchangeable individuals may be inappropriate by definition.

Given that models that assume either a single consensus tree, a fixed rate of recombination across strains, or exchangeable individuals, are all clearly at odds with the data on prokaryotic genome evolution, this raises the question of what would be an appropriate mathematical ‘null model’ that can capture the statistics that we observed here. In such a model, almost every lineage must have distinct rates of recombination with other lineages, these rates must vary over multiple orders of magnitude, and the model should reproduce the roughly power-law distributions of *n*-SNP frequencies, ideally with exponents that can be tuned by parameters in the model. It is currently unclear how to construct such a model.

Apart from the question of how to mathematically model the observed patterns, a second key question is what determines these highly variable relative rates of recombination across lineages. For example, it is currently not clear whether the differences in rates are shaped mainly by natural selection, e.g. that due to epistatic interactions only recombinant segments from other strains with similar ‘ecotypes’ are not removed by purifying selection, as suggested by some previous works [24, 35], or that recombination rates may be set mostly by parameters such as the frequency with which lineages co-occur at the same geographical location. It is also conceivable that phages are a major source of transfer of DNA between strains, so that recombination rates may reflect the rates at which different lineages are infected by the same types of phages. It is also noteworthy that homologous recombination requires sufficient homology between the endpoints of the DNA fragment and the homologous segment in the host genome. Thus, recombination rates will intrinsically decrease with the nucleotide divergence between strains and previous studies have estimated that the rate of successful recombination decreases exponentially with nucleotide divergence [36, 37]. In this regard it is interesting that the critical divergence at which half of the genome is recombined varies over a relatively small range, i.e. from 0.003 — 0.01 (Fig. 7A). It is thus conceivable that a species is essentially defined by the collection of strains that are sufficiently close to allow efficient recombination [38]. However, the statistics reported here seem to suggest a much larger range of relative recombination rates than such a simple DNA-homology based model would predict.

While we here studied the frequency distribution of *n*-SNP types as well as the entropies *H_n_*(*s*) of the *n*-SNP distributions for each strain, it appears to us that this is just the tip of the iceberg of possible ways in which *n*-SNPs can be used to study the evolution of a set of strains from their core genome alignment. Our analyses indicate that prokaryotic genome evolution is driven by recombination that occurs at a very wide distribution of different rates between different lineages, and there is now a strong need for identifying what sets these rates, and the development of new mathematical tools and models that can accurately describe this kind of genome evolution.

## Materials and Methods

### Data

The *E. coli* sequences analyzed here can be accessed on NCBI Bioproject via the accession number PRJNA432505 [7,19]. These genomes were sequenced with Illumina HiSeq 2000 technology. Samples were multiplexed, 24 per lane, producing 100bp paired end reads which resulted in an average coverage depth of between 125x and 487x (with 4 strains over-represented at more than 1000x). In Table S5 strains names and details for the reference strains used for Figure S1 can be found.

Genome sequences for all other species were downloaded from ftp.ncbi.nlm.nih.gov/genomes/refseq/bacteria/. All strain names and download dates are listed in Table S6.

### Core genome alignment and core tree

To build a core genome alignment for the SC1 strains we used the Realphy tool [20] with default parameters and Bowtie 2 [39] for the alignments. The Illumina sequencing used to sequence the *E. coli* strains can still make sequencing errors at a certain rate and obtaining accurate base calls genome-wide relies on having sufficient read coverage at each position to confidently call the consensus nucleotide. We note that, in order to avoid sequencing errors, Realphy is quite conservative in its construction of the core genome alignment. In particular, in order for a position to be included each strain has to be represented, the coverage has to be at least 10 in each strain, and at least 95% of the reads from each strain have to agree on the nucleotide. A rough bound on the rate of sequencing errors can be obtained by noting that some of our strains are extremely closely-related, i.e. while all 92 strains are unique in their full genome, there are only 82 unique core genomes. This means that there are 10 genomes that are identical in the core to another genome. Given that the length of the core alignment is *L* = 2’756’541, this means there were 10 × *L* nucleotides sequenced without a sequencing error and a simple Bayesian calculation shows that this implies that, with 99% posterior probability, the rate of sequencing errors is less than *μ_s_* = 1.1 × 10^-7^.

Realphy used PhyML [21] with parameters -m GTR -b 0 to infer trees from the whole and parts of the core alignment. The tree visualizations were made using the Figtree software [40].

### Analysis of core alignment blocks

For each 3 Kb block of the core alignment we used PhyML using the option -c 1 to infer a phylogeny while restricting the number of relative substitution rate categories to one. Furthermore, to calculate the log-likelihood of a given 3Kb block under the tree topologies of other blocks, we reran PhyML using the -o ‘lr’ option, which only optimizes the branch lengths as well as the substitution rate parameters but doesn’t alter the topology of the phylogeny.

### Pairwise analysis and mixture modeling

For each pair of strains we slide a 1Kb window over the core genome alignment of the pair, shifting by 100 bp at a time, and build a histogram of the number of SNPs per kilobase by counting the number of SNPs in each window. That is, we obtain the distribution *P_n_* of the fraction of 1 Kb windows that have *n* SNPs. We then assumed that the 1 kilobase blocks can be separated into a fraction *f_a_* of ‘ancestral blocks’, i.e. regions that were inherited from the clonal ancestor of the pair, and a fraction (1 — *f_a_*) that have been recombined since the pair diverged from a common ancestor. Although in previous work a simple *ad hoc* scheme was used in which it was assumed that blocks with less than a particular number of SNPs are ancestral and blocks with more SNPs are recombined [16], we found that this approach is not satisfactory and results significantly depend on the cut-off chosen.

We thus decided to employ a more principled mixture model approach. For the ancestral regions, the number of SNPs per kilobase should follow a simple Poisson distribution *P_n_* = *μ^n^e^-μ^*/*n*!, with *μ* the expected number of mutations per block. For the recombined regions, we note that these regions themselves will consist of mosaics of sub-regions that have been recombined. Consequently, the recombined regions will consist of a mixture of Poisson distributions with different rates. It is well-known that mixtures of Poisson distributions with rates that are (close to) Gamma-distributed follow a negative binomial distribution and we found empirically that negative binomial distributions give excellent fits to the observed SNP distributions in our data. For the recombined regions we thus assume a negative binomial distribution of the form

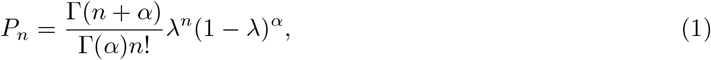

where 0 ≤ λ ≤ 1 and *α* ≥ 1 are parameters of the distribution. We thus fit the observed distribution of SNPs per block *P_n_* using the following mixture:

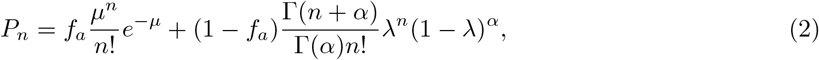

where *f_a_* is the fraction of the genome that is ancestral. Fits were obtained using maximum likelihood. While expectation maximization was used to fit the parameters *f_a_, μ*, and *λ*, a grid search was employed to find the optimal dispersion parameter *α*.

Note that, in terms of the fitted parameters, the total number of mutations in ancestral blocks is *μf_a_*, and the number of mutations in recombined blocks is (1 – *f_α_*)*αλ*/(1 – *λ*).

To estimate the lengths of recombination events, we first extracted pairs that are sufficiently close (divergence less than 0.002) such that multiple overlapping recombination events are unlikely. We then used a two-state HMM with the same two components, i.e. a Poisson and a negative binomial component corresponding to ancestral and recombined segments, and having fixed transition rates from the ancestral to the recombined state and vice versa, to parse the pairwise alignment into ancestral and recombined segments. Note that the parameters of the HMM are fitted separately for each pair of strains. For each pair, we took as recombined segments those contiguous stretches that were assigned to the recombined state by the HMM.

We define mostly clonal pairs as pairs with more than 90% of the alignment classified as ancestral, fully recombined pairs as pairs with less than 10% of the alignment classified as ancestral, and all other pairs as transition pairs. In order to estimate the critical divergence at which half of the genome is recombined we fit a linear model to the observed relationship between divergence and clonal fraction in all transition pairs, and define the critical divergence as the divergence at which the linear fit has a clonal fraction of 50%. To calculate the fraction of mutations that derive from recombined segments at the critical divergence we compute the fraction of mutations in recombined segments for all transition pairs (using the results from the mixture model) and fit a linear model to the observed dependence between the ancestral fraction an the fraction of mutations in recombined segments. We then define the fraction of mutations in recombined regions at the critical divergence as the fraction of mutations in the linear fit when the ancestral fraction is 50%.

### Simulated data sets

In order to simulate genome evolution under a simple evolutionary model that includes drift, mutation, and recombination, we developed new software (manuscript in preparation) based on general purpose GPU programming (GPGPU), which is available from https://github.com/thomsak/GPUprokEvolSim. Our software explicitly simulates the evolution of a population of *N* DNA sequences of length *L_g_* using a Wright-Fisher model with non-overlapping generations. In order to be able to not only evolve the sequences, but also track recombination events occurring along the clonal history of a sample of *S* genomes from the population of *N*, we proceed as follows.

The evolution of the *n* genomes is simulated for 8*N* generations. First, a ‘guide’ clonal history for the sample of *S* sequences is determined using the Kingman coalescent [6] for *S* individuals within the population of *N*. In particular, for the subset of *S* genomes, their ancestry is determined for 8*N* generations into the past. Notably, because the *S* sequences will have a common ancestor much more recently than 8*N* generations in the past, for many of these generations there will only be a single ancestor.

We then simulate 8*N* generations of the evolution of *n* genomes of length *L_g_* forward in time by iterating:

1. For each individual corresponding to an ancestor from the clonal phylogeny of the sample of *S* genomes, its parent in the previous generation is chosen according to the guide clonal phylogeny. For each other individual, a random parent is chosen from the previous population of *N* individuals.
2. For each individual of the new generation, the genome of the ancestor is copied, scanned from left to right, and at each position a recombination event is initiated with probability *ρ*. If a recombination is chosen to occur at position *i*, one of the *N* — 1 other members of the population is chosen at random, and the section of its genome from position *i* +1 to *i* + *L_r_* is copied into the current genome. After this, each position in the genome is mutated with probability *μ*. The target nucleotide is chosen randomly using a transition-transversion ratio of 3-to-1.

Apart from tracking the *N* genome sequences, we also track, for each branch of the clonal phylogeny of the *S* individuals, how many times each position was overwritten by recombination during the evolution along the branch. This allows us to calculate what fraction of positions are inherited clonally along each branch, how many times each position in the final alignment of *S* genomes was overwritten by recombination during its clonal history, and what fraction of positions was clonally inherited for each pair of strains.

#### Parameter settings

To keep the simulations computationally feasible, we simulated a population of *N* = 3600 individuals. To make the simulation directly comparable with the data from *E. coli* we focused on samples of *S* =50 individuals, used a genome size of *L_g_* = 2’560’000 bp, and used a size of *L_r_* = 12’000 bp for the recombined fragments. Note that this length of recombined fragments is toward the lower range of the sizes of recombined fragments estimated from close pairs of *E. coli* strains. For a Kingman coalescent, the expected fraction of columns that is not polymorphic in an alignment of *S* individuals is given by

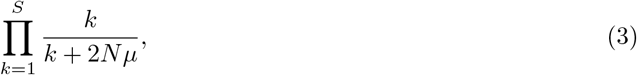

and we set *μ* such that this fraction is 0.9, i.e. similar to what is observed for the alignment of *E. coli* strains. In particular we set *μ* = 3.28 × 10^-6^ so that *Nμ* = 0.012.

Finally, apart from simulations without recombination, i.e. *ρ* = 0, we performed simulations with 6 different recombination rates, corresponding to ratios of recombination to mutation of *ρ/μ* = 0.001, *ρ/μ* = 0.01, *ρ/μ* = 0.1, *ρ/μ* = 0.3, *ρ/μ* = 1, and *ρ/μ* = 10, i.e. covering 4 orders of magnitude. Note that the clonal phylogeny for the subset of *S* strains was created once, and then used in all simulations with different recombination rates to facilitate direct comparisons.

### Estimating the fraction of SNPs that correspond to single substitution events

Our analyses assume that bi-allelic SNPs correspond to single substitution events in the evolutionary history of the genomic position. This assumption may break down due to apparent SNPs caused by sequencing errors, as well as due to homoplasies, i.e. bi-allelic columns for which more than one substitution event occurred. We here estimate the frequency of both types of events.

#### SNPs due to sequencing errors are negligible

Above we estimated, from the fact that we observe 10 strains that are identical in their core genome to another strain, that the rate of sequencing errors is at most *μ_s_* = 1.1 × 10^-7^. Using this, we expect a sequencing error to occur in at most *L* (1 – (1 – *μ_s_*)^92^) = 28 of the columns of our length *L* = 2’756’541 core genome alignment. Note that, because the fraction of columns with more than 1 sequencing error is negligible, all these sequencing errors will produce a mutant nucleotide in only 1 strain, because the expected columns with two sequencing errors is negligible. Thus, if a sequencing error occurs in a column that is otherwise non-polymorphic, it will create a bi-allelic SNP in which only 1 strain carries a differing base and such SNPs do not affect any of the phylogenetic analyses. Thus, sequencing errors affecting the phylogenetic analysis only occur when the sequencing error occurs in a SNP column, and creates one of the two nucleotides that already existed. Since only about 10% of columns are polymorphic, the expected number of SNP columns that are informative for phylogeny and affected by sequencing errors is at most 28/10 = 2.8. Given that there are 247’822 phylogeny informative SNPs in total, the expected fraction of these that are affected by sequencing errors is roughly 10^-5^.

#### Estimating the rate of homoplasy

The relatively low frequency of SNPs and the fact that almost all SNPs are bi-allelic strongly suggests that almost all bi-allelic SNPs correspond to single substitution events in the evolutionary history of the position. Here we use a simple substitution model to estimate the fraction of bi-allelic SNPs that correspond to multiple substitution events, i.e. homoplasies. To do this we will analyze the observed frequencies of columns with 1, 2, 3, and 4 different nucleotides under a simple substitution model. Note that, in this simple model we assume that all substitutions are neutral, so that there is essentially no difference between mutations and substitutions. In the real data some mutations are deleterious and some of these are removed by purifying selection, leading to lower rates of substitutions at some positions than at others. In particular, we observed that SNPs at synonymous sites are almost ten-fold more common than at second positions in codons. To assess the effects of selection, we will consider the frequencies of columns with different numbers of nucleotides both for the subset of positions that should be under relatively little selection, i.e. third positions in fourfold degenerate codons, and positions that should be under relatively strong selection, i.e. second positions in codons. Since a large fraction of all SNPs corresponds to SNPs in which the outgroup of 9 strains has a nucleotide that differs from the nucleotide of all other strains, we will also do this separately for all strains, and all strains minus the 9 strains of the outgroup

For a given position in the alignment, let *μ* denote the product of the mutation rate times the total length of the branches in the phylogeny at that position. The variable *μ* thus corresponds to the expected number of mutations in the evolutionary history of this position. The probability that *n* mutations took place at this position is given by a Poisson distribution:

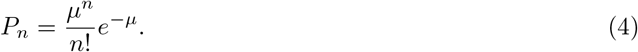

We will assume a simple substitution model in which transitions are r times as likely to occur as transversions. Let *d* denote the number of different nucleotides in the column and let *T*(*d*’|*d*) be the matrix of probabilities, that under a single mutation, the number of different nucleotides transitions from *d* to *d*’. We have *T*(2| 1) = 1, *T*(2|2) = (2 + *r*^2^)/(2 + *r*)^2^, *T*(3|2) = (2 + 4*r*)/(2 + *r*)^2^, *T*(3|3) = 2/3, *T*(4|3) = 1/3, *T*(4|4) = 1, and all other transition probabilities are zero. Starting from a single nucleotide in the column, the probability *P*(*d|n*) to end up with *d* different nucleotides after *n* mutations is given by the *n*-th power of the transition matrix *T*, i.e. *P*(*d|n*) = *T*^*n*^(*d*|1). From this we can work out the probability *P*(*d|μ*) to end up with *d* different nucleotides as a function of the expected number of mutations *μ* as

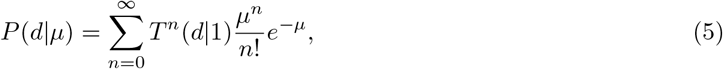

which can be easily numerically evaluated to sufficient precision.

Assume we observe *c_d_* columns with *d* different nucleotides, with *d* running from 1 to 4. The log-likelihood of this count data given *μ* is

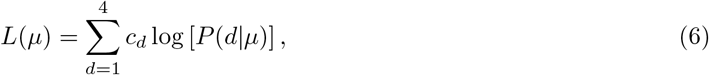

and by maximizing this likelihood with respect to *μ* (numerically), we obtain an estimate *μ*\*. Finally, given *μ*\*, the fraction *f_h_* of homoplasies, i.e. bi-allelic SNPs that correspond to multiple substitutions, is given by

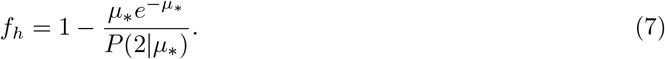

Table 1 shows the estimated expected number of mutations per column *μ*_*_ and the estimated fraction of homoplasies *f_h_* for the 5 different subsets of columns, using a transition-to-transversion ratio of *r* = 3. We see that the fraction *f_h_* is at most 6.3% and less than 1% for second positions in codons.

**Table 1.** Estimated expected number of mutations per position *μ*_*_ and estimated fraction of homoplasies for 5 different subsets of core alignment columns: all columns, all synonymous positions (third positions in fourfold degenerate codons), second positions in codons, synonymous positions excluding the outgroup, second positions in codons excluding the outgroup.

| Column set | $\mu_*$ | $f_h$ |
| --- | --- | --- |
| All columns | 0.118 | 0.026 |
| Synonymous positions | 0.287 | 0.063 |
| Second positions in codons | 0.0258 | 0.006 |
| Synom. pos. without outgroup | 0.149 | 0.033 |
| Sec. pos. without outgroup | 0.0172 | 0.004 |

In addition, Supplementary Fig. S26 shows a comparison of the observed and predicted frequencies of columns with 1, 2, 3, and 4 letters. Since effects of selection are likely least for the synonymous positions, we expect the simple model to fit the data best and we indeed observe that, for the synonymous positions, the simple model can reasonably accurately fit the observed frequencies, and even for the set of all alignment columns the fits are quite accurate (Suppl. Fig. S26). In contrast, for the second positions in codons, we can see the effects of selection in that, from the larger fractions of columns without SNPs, the model infers a lower *μ*\*, and this leads to an underestimation of columns with 4 nucleotides. Thus, the true fraction *f_h_* is more likely close to the values inferred from the synonymous positions. Note that *f_h_* = 0.063 when including the outgroup and *f_h_* = 0.033 when the outgroup is excluded. The difference between these two estimates derives from the very high fraction of SNP columns in which the 9 strains of the outgroup have another nucleotide than all other strains. For this subset of SNPs the fraction of columns that have more than one mutation is much higher than for any other SNP column. Thus, for all other SNP columns, the estimate that a fraction of 3.3% correspond to homoplasies is likely the most accurate.

As an additional test, we also applied this simple method to estimate the rate of homoplasy for the simulation data. For the simulation data without recombination we explicitly kept track of homoplasies and determined that a fraction *f_h_* = 0.0256 of bi-allelic positions correspond to homoplasies. Applying our estimation method to the alignment of the *S* = 50 sample genomes from this simulation, we estimated *f_h_* = 0.0243, which is within 2% of the true value, further supporting our method for estimating the homoplasy rate.

In summary, our estimates strongly suggest that the rate of homoplasies among bi-allelic SNPs in our core genome alignment of the *E. coli* strains lies somewhere in the range of 2 — 6%.

### Removing potentially homoplasic positions from the core genome alignment

Even though the analysis of the previous section has shown that homoplasies are only a very small fraction of all bi-allelic SNPs, we decided to investigate to what extent this small fraction of homoplasies may affect the various statistics that we calculate. Ideally, if we knew which alignment columns correspond to homoplasies, we could simply remove all homoplasic columns and recalculate all statistics of interest on the reduced alignment from which these homoplasic sites were removed. However, since we only know the approximate fraction *f_h_* of homoplasies and do not know which columns are homoplasies, a conservative approach is to remove those columns that are most phylogenetically inconsistent with other columns in their neighborhood.

In particular, for each bi-allelic SNP s in the alignment, we check its phylogenetic consistency with the nearest 200 SNP columns to the left and nearest 200 SNP columns to the right, i.e. its consistency with others SNPs within a roughly 4 kilobase region. We then assign each SNP column an inconsistency score *I_s_* corresponding to the fraction of the 400 neighboring columns that are phylogenetically inconsistent with it. We then sort all SNP columns by their inconsistency *I_s_* and remove a fraction f of SNP columns with the highest inconsistency. After this, we can recalculate all statistics of interest on the alignment from which these potentially homoplasic columns have been removed. In particular, we generated reduced alignments from which a fraction *f* = 0.05 and a fraction of *f* = 0.1 of most inconsistent columns were removed.

### Constructing a tree that maximizes the number of compatible SNPs

We classify all SNPs in the core genome alignment into *SNP types* as follows. For each bi-allelic SNP, we map all letters with the majority nucleotide to a 0 and the minority nucleotide to a 1 and sort the bits according to the alphabetic order of the strain names. For SNPs where one allele occurs in exactly half of the strains the minority allele is not defined and the ambiguity is resolved by setting the first bit of the string to 0. In this way, each SNP is mapped to a binary sequence of length 92. This binary sequence defines the SNP type. Note that a SNP type corresponds to a particular bi-partition of the strains.

We next counted the number of occurrences *n_t_* of each SNP type *t* and sorted the SNP types from most to least common. We then used the following greedy algorithm to a collect a subset *T* of mutually compatible SNP types that accounts for as many SNPs as possible. We seed *T* with the most common SNP type, i.e. the SNP type occurring at the top of the list. We then go down the list of SNP types, iteratively adding SNP types *t* to the set *T* that are compatible with all previous types in the set *T*.

### Bottom up tree building

In this procedure we build phylogenies of sub-clades in a bottom-up manner, starting from the full set of 92 strains and iteratively fusing pairs, minimizing the number of incompatible SNPs at each step.

For any subset of strains *S*, we define the number of supporting SNPs *n_S_* as the number of SNPs that fall on the branch between the subset *S* and the other strains, i.e. the number of SNPs in which all strains in *S* have one letter, and all other strains another letter. Similarly, we define the number of clashing SNPs *c_S_* as the number of SNPs that are incompatible with the strains in *S* forming a subclade in the tree.

The iterative merging procedure is initiated with each of the 92 strains forming a subclade *S*. At each step of the iteration we calculate, for each pair of existing sub-clades *S*_1_, *S*_2_, the number of clashing SNPs *c_S_* and supporting SNPs *n_S_* for the set of strains *S = S*_1_ ∪ *S*_2_ consisting of the union of the strains in *S*_1_ and *S*_2_. We then merge the pair (*S*_1_, *S*_2_) that minimizes the clashes *c_S_* and, when their are ties, maximizes the number of supporting SNPs *n_S_*. At each step of the calculation we keep track of the total number of SNPs on the branches of the subtrees build so far, as well as the total number of SNPs that are inconsistent with the subtrees build so far. In addition, we calculate the average pairwise divergences of the strains within the sub-clades. Supplementary Fig. S9 shows the ratio of clashing to supporting SNPs as a function of the divergence within the sub-clades.

### Quartet analysis

Quartets were assembled in the following way. We construct a grid of target distances *d* starting at 0.00001 and having 50 points with 0.0005 sized distance. For every target distance *d* we scan the alignment for four strains which have all pairwise distances within 1.25 fold of distance *d*. Every target distance *d* for which no quartet can be found fulfilling these criteria is ignored.

For each quartet we extract all SNP columns where two strains have a specific nucleotide and the other two strains have another nucleotide. Every such SNP column unambiguously supports one out of three possible tree topologies for this quartet. For each quartet we determine which topology has the largest number of supporting SNPs, and what the fraction of SNPs is that support this topology.

### Linkage Disequilibrium measure

A standard measure of linkage disequilibrium of SNPs at a given distance is given by the average squared-correlation of the genotypes at these positions [41]. For a pair of loci with bi-allelic SNPs there are 4 possible genotypes which we indicate as binary patterns 00, 01, 10, and 11. If the frequencies of these genotypes are *f*_00_, *f*_o1_, *f*_10_, and *f*_11_, then the squared correlation is calculated as

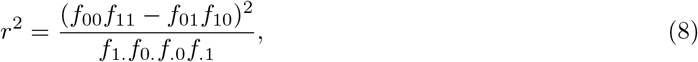

where the variables with dots correspond to marginal probabilities, e.g. *f*_1_. = *f*_10_ + *f*_11_, *f*._1_ = *f*_01_ + *f*_11_, and so on.

### Distribution of tree-compatible stretches

To calculate the distribution of the number of consecutive tree-compatible SNPs we start from each SNP *s* in the core genome alignment and count the number *n_s_* of SNP columns immediately following *s*, until a SNP column occurs that is incompatible with at least one of the *n_s_* SNP columns. Similarly, to obtain the distribution of the number of consecutive tree-compatible nucleotides we start from each position *p* in the core genome alignment and count the number *n_p_* of consecutive nucleotides until a SNP column occurs that is incompatible with at least one of the SNP columns among the *n_p_* nucleotides.

### Minimum number of phylogeny changes *C*

We iterate over all SNP columns along the core genome alignment and add the current SNP to a list if it is pairwise compatible with all SNPs currently in the list. If it is incompatible with at least one SNP in this list we empty the list, re-initialize the list with the current SNP, and increase the phylogeny counter *C* by one.

### Phylogenies of human genome sequences

We selected 40 individuals at random from the 1000 Genome project [32] and build a ‘core tree’ by applying PhyML to the sequences of chromosomes 1 through 12. To investigate the robustness of this core tree, we cut the alignment in 1000 bp blocks and did 100 random resampling of half of the blocks, building a phylogeny for each resampling using PhyML. We then determined the fraction of times each split in the core tree occurred in the trees of the 100 resamplings.

### Power-law fits of *n*-SNP distributions

We extract each *n*-SNP from the core genome alignment and count the frequency, i.e. the number of occurrences, *f_t_* of each *n*-SNP type *t* as well as the total number *T* of *n*-SNP types that occur at least once. We assume the *n*-SNP type occurrences are drawn from a power-law of the form

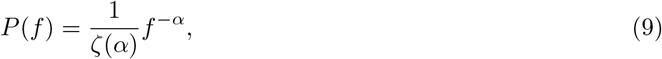

where *ζ*(*α*) is the Riemann zeta function defined by

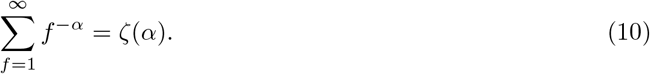

The log-likelihood of the frequencies *f_t_* as a function of *α* is given by

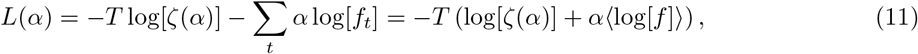

where 〈log[*f*]〉 is the average of the logarithm of the SNP-type frequencies. Using a uniform prior on *α*, the posterior distribution of *α* is simply proportional to the likelihood function. The optimal exponent *α*\* is the solution of

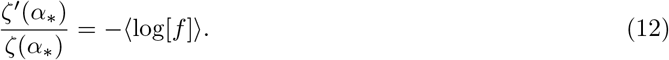

To calculate error-bars on the fitted exponentials we approximate the posterior by a Gaussian by expanding the log-likelihood to second order around the optimal exponent *α*\*. We then find for the standard-deviation of the posterior distribution:

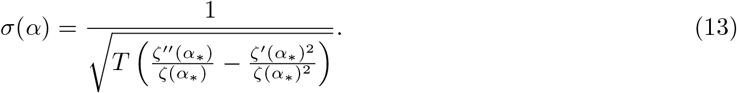

### Entropy profiles of *n*-SNP distributions

For a given strain *X* we first extract all SNP types *t* for which *X* is one of the strains that shares the minority nucleotide. We then further stratify these SNP types by the number of strains *n* sharing the minority nucleotide. For each *n* we thus obtain a set *S*(*X, n*) of *n*-SNPs in which strain *X* is one of the strains sharing the SNP. We denote the number of occurrences of a SNP of type *t* by *f_t_* and the total number of *n*-SNPs within set *S*(*X, n*) as |*S*(*X, n*)|, i.e.

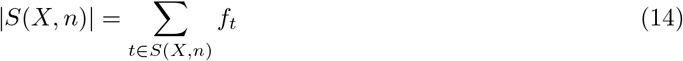

The entropy *H*(*X, n*) of the *n*-SNP distribution of strain *X* is then defined as

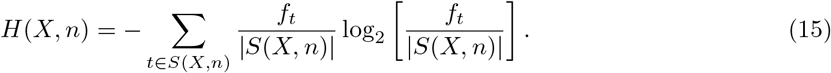

### Comparing *n*-SNP statistics of pairs of strains

The considerable variability of the entropy profiles of *E. coli*’s strains suggests that the lineages of different strains must have recombined at different rates with other lineages. Indeed, we observe much less variation in the entropy profiles of the data from simulations in which each strain recombines at an equal rate with each other strain, than we observe for the *E. coli* data (Suppl. Fig. S22). However, since we currently lack a concrete evolutionary model that can reproduce all the statistics that we observe in the data, it is difficult to quantify how different the recombination rates of different lineages have to be in order to reproduce the observed variation in entropy profiles.

Nonetheless, it is straight-forward to define a simple statistical measure of the difference in the *n*-SNP statistics of a given pair of strains (*x,y*). Each SNP in the core genome alignment can be categorized by the subset of strains *S* that share the minority allele. Given a pair of strains (*x,y*), this subset *S* can take on 4 possible types: *S*_0_ = (*Z*), *S*_2_ = (*xyZ*), *S_x_* = (*xZ*), and *S_y_* = (*yZ*), where *Z* is a subset of strains that does not include *x* or *y*. That is, either neither *x* or *y* carry the minority allele, they both do, or only one of them does. We are interested in comparing the relative frequencies with which *x* and *y* co-occur with different subsets *Z* in SNPs across the alignment. Note that to this end, we can ignore SNPs of the type *S*_0_ and *S*_2_ because *x* and *y* occur together in these SNPs, so that the frequency with which different subsets *Z* occur in types *S*_0_ and *S*_2_ is per definition the same for both *x* and *y*. Thus, the relevant SNPs are of the type *S_x_* and *S_y_*.

Let *n_xZ_* be the number of SNPs of the type *S_x_* = (*xZ*), *n_yZ_* the number of SNPs of type *S_y_* = (*yZ*), and the totals *N_x_ ∑_Z_ n_xZ_*, and *N_y_* = ∑_*Z*_ *n_yZ_*. We can then define two probability distributions over all possible subsets *Z*, i.e. 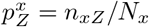 and 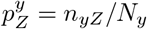. These two probability distributions give the relative frequencies with which *x* and *y* are observed to occur in SNPs with all other sets of strains *Z* and there are standard methods to quantify to what extent these two probability distributions are statistically significantly different. A standard orthodox statistical test is the Fisher exact test, which uses the test statistic

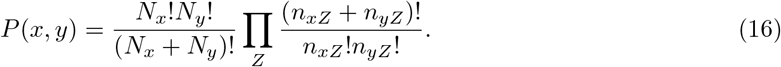

Note that, if the counts are large enough so that we can use the Stirling approximation log(*n*!) ≈ *n* log(*n*)—*n*, this can be rewritten as

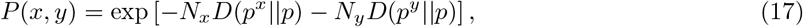

where the distribution *p* is the marginal distribution over the sets *Z*

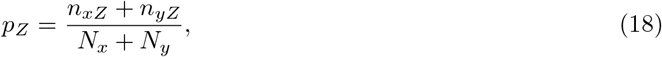

and *D*(*p||q*) is the Kullback-Leibler divergence (or relative entropy) of the distribution *p* with respect to distribution *q*. Note that the probability *P*(*x,y*) corresponds directly to the *p*-value of the Fisher exact test. The cumulative distribution of *p*-values across all pairs of strains is shown in Suppl. Fig. S21, left panel, showing that the *n*-SNP profiles are different for about 95% of all pairs. In the right panel we show a scatter of the distance — log[*P* (*x,y*)] of the *n*-SNP frequency profiles of each pair of strains (*x,y*) as a function of their nucleotide divergence *d*(*x,y*). This shows that only very close pairs that differ by less than 10 SNPs in their core genomes have statistically identical *n*-SNP frequency profiles. In addition, there is a very high correlation between the nucleotide divergence *d*(*x,y*) and the distance — log[*P*(*x,y*)] of the *n*-SNP frequency profiles.

Finally, one caveat of the Fisher exact test is that it presumes that all the observed *n*-SNPs of the types (*Zx*) and (*Zy*) are statistically independent, and in the absence of a concrete stochastic model for the evolutionary dynamics, we do not know whether this assumption holds. However, the *p*-values for most pairs are so low that, even if we assume the true numbers of independent events are 500-fold less, the fraction of significantly different pairs would still be larger than 90%.

## Acknowledgments

The authors thank Olin Silander and Diana Blank for their laboratory work on the sequencing of the SC strains. During the development of this work, the authors have benefited from discussions with many researchers including Olin Silander, Frederic Bertels, Sergei Maslov, Purushottam Dixit, Edo Kussell, Boris Shraiman, Eugene Koonin, Daniel Fisher, Paul Rainey, Oskar Hallatschek, Mikhail Tikhonov, Richard Neher, Bruce Levin, Otto Cordero, and Daniel Weissman. This work was supported by the Swiss National Science Foundation grant No. 31003A_135397. In addition, this work was done in part while the authors were visiting the Simons Institute for the Theory of Computing and was thus supported in part by NSF Grant No. PHY17-48958, NIH Grant No. R25GM067110, and the Gordon and Betty Moore Foundation Grant No. 2919.01. Calculations were performed at sciCORE (http://scicore.unibas.ch/) scientific computing core facility of the University of Basel.

## S1 Supplementary Figures, Tables, and Texts

**Figure S1.**
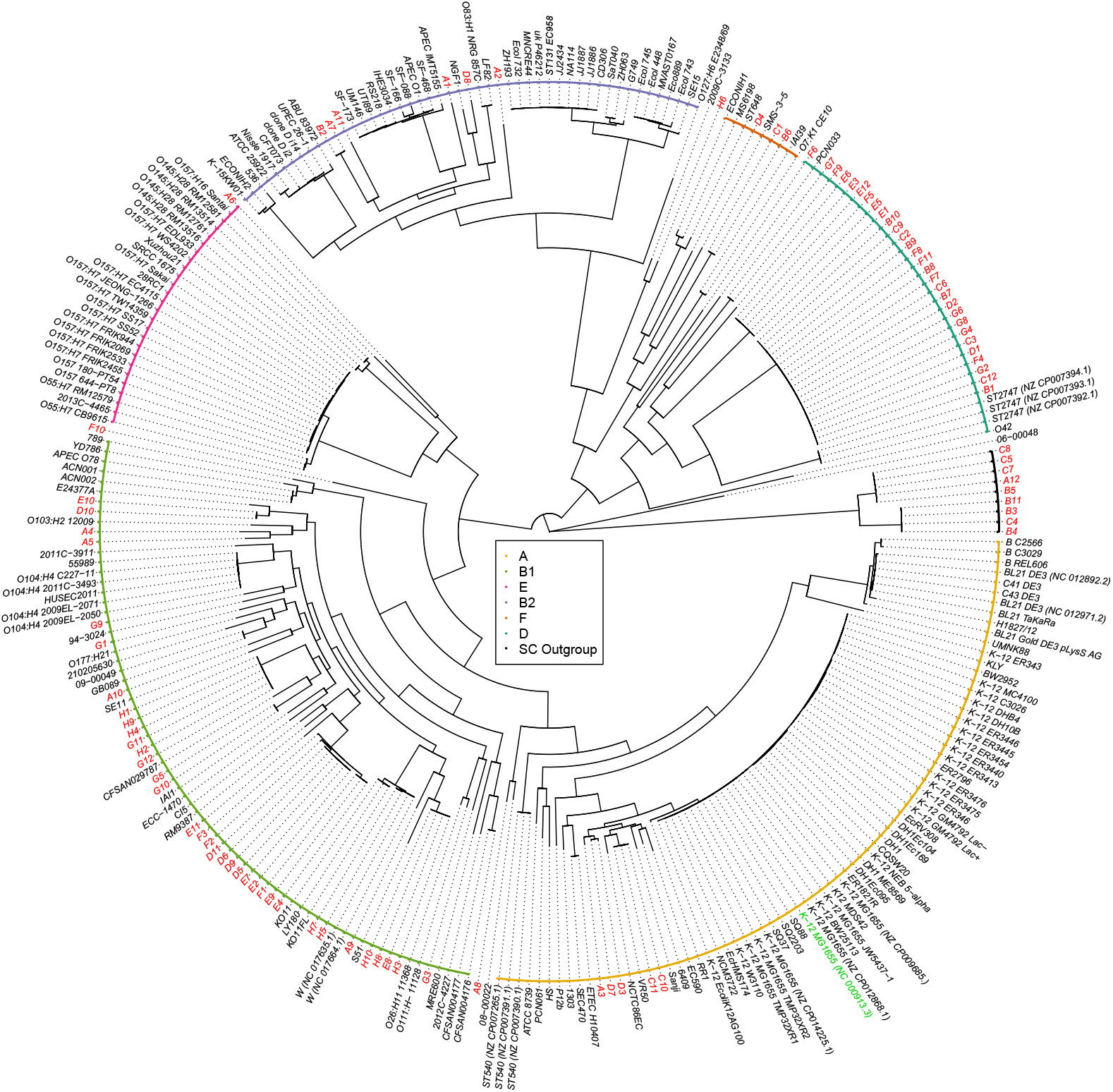
Maximum likelihood tree reconstructed from the core genome alignments of the SC1 strains (red font names), the K-12 lab strain (green font name), and 189 *E. coli* reference strains (black font names). Known phylogroups are indicated as different colored leaf nodes. The SC1 strains are distributed across essentially all known phylogroups, include strains that cannot be easily assigned to a phylogroup, and a distal ‘outgroup’ of 9 strains (black leafs) that are at 8% nucleotide divergence from other strains (branch shortened).

**Figure S2.**
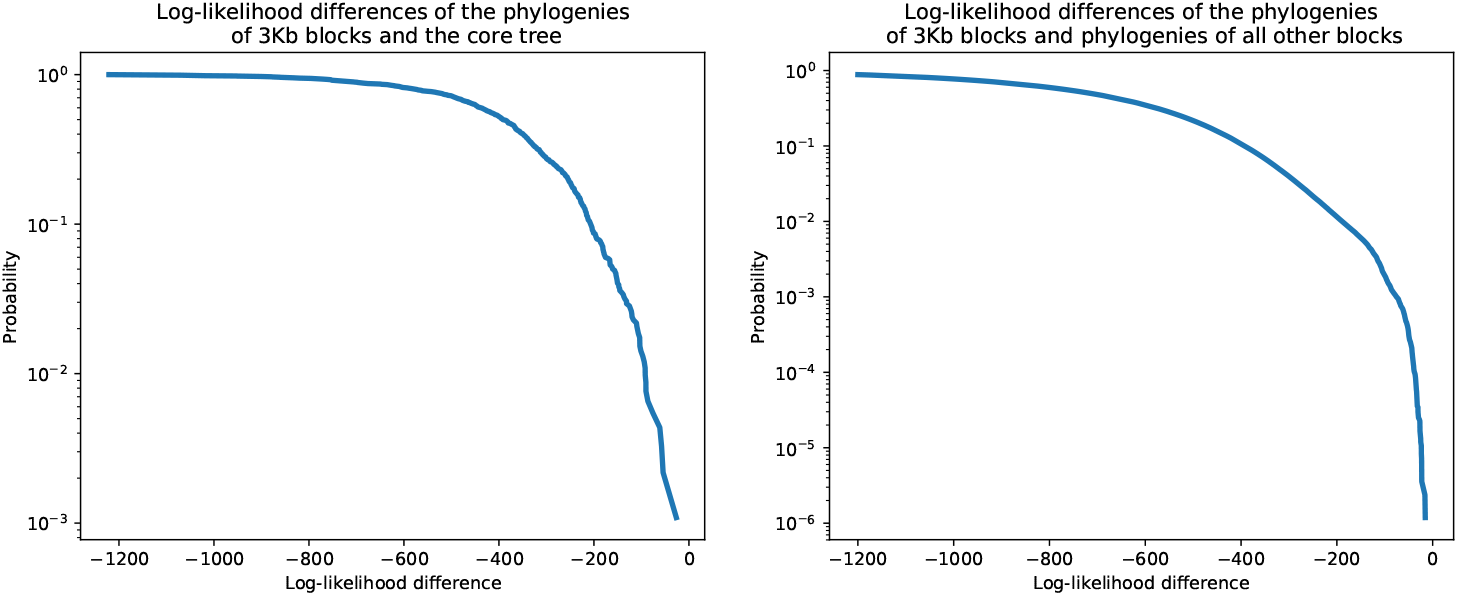
All 3kb alignment blocks reject the core tree topology as well as the topologies of the phylogenies reconstructed from all other blocks. **Left panel**: For each 3Kb block in the core alignment we used PhyML to reconstruct a phylogeny and then calculated the difference in the log-likelihood of the alignment block under the topology of the core tree and the log-likelihood of the reconstructed phylogeny. The figure shows the reverse cumulative distribution of these log-likelihood differences, with the vertical axis shown on a logarithmic scale. There are virtually no blocks for which the log-likelihood of the core tree topology is close to the log-likelihood under the block’s own phylogeny. **Right panel**: As in the left panel, but now we calculated, for each 3kb block, the log-likelihood differences for the topologies of the phylogenies reconstructed from all other blocks. Each block attains a substantially higher log-likelihood when using its own topology than using the topology of any other alignment block.

**Figure S3.**
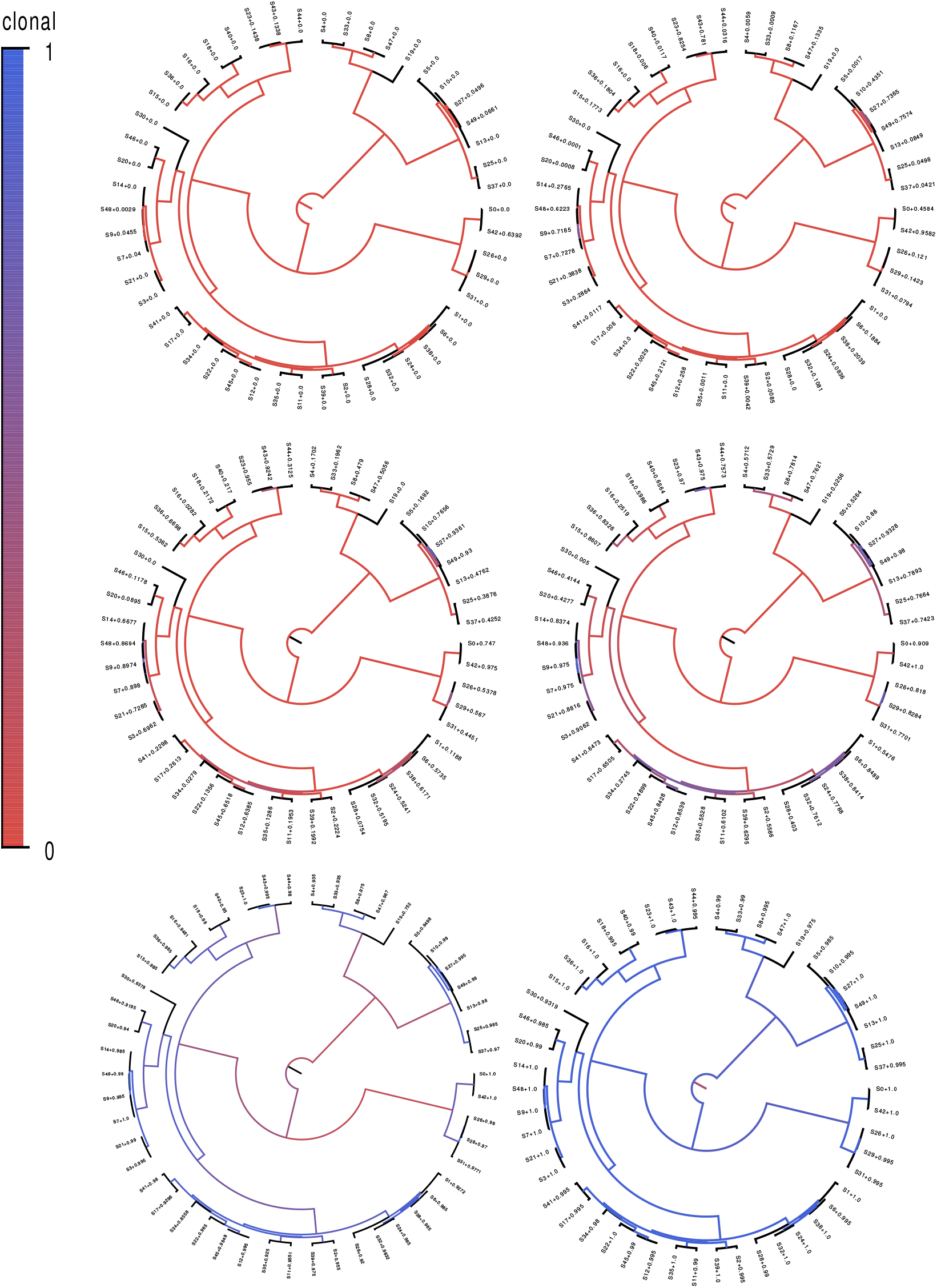
Fractions of positions that are clonally inherited (not affected by recombination) along each of the branches of the clonal phylogeny for the simulated datasets. Each panel shows the clonal phylogeny with the color of the branches indicating what fraction of positions were clonally inherited from all (blue) to none (red) along that branch. Panels correspond, from top-left to bottom-right, to recombination to mutation rate of *ρ/μ* equalling 10, 1, 0.3, 0.1, 0.01, and 0.001. The fractions at the leafs indicate the fraction of the genome that was clonally inherited along each terminal branch.

**Figure S4.**
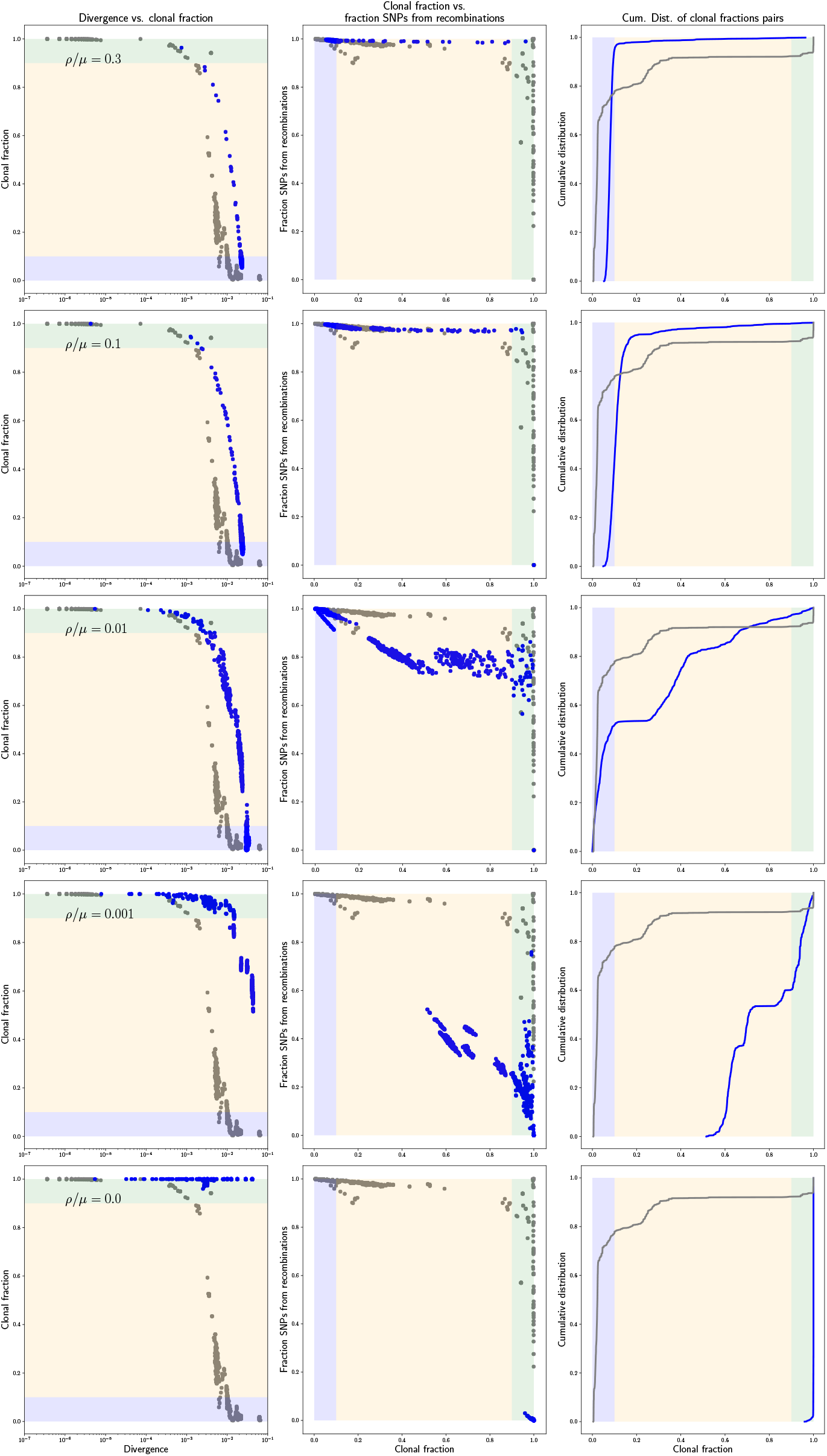
Statistics of the pairwise analysis on the simulated data. Each row of panels corresponds to simulations performed with a different recombination rate (indicated in each row) and shows, in blue, the same results as in panels G, H, and I of Fig. 2, i.e. the fraction of the genome that was clonally inherited as a function of divergence for each pair, the fraction of all mutations that derive from recombination as a function of the clonally inherited fraction, and the cumulative distribution of clonally inherited fractions. For reference, the corresponding results for the *E. coli* strains are shown in grey.

**Figure S5.**
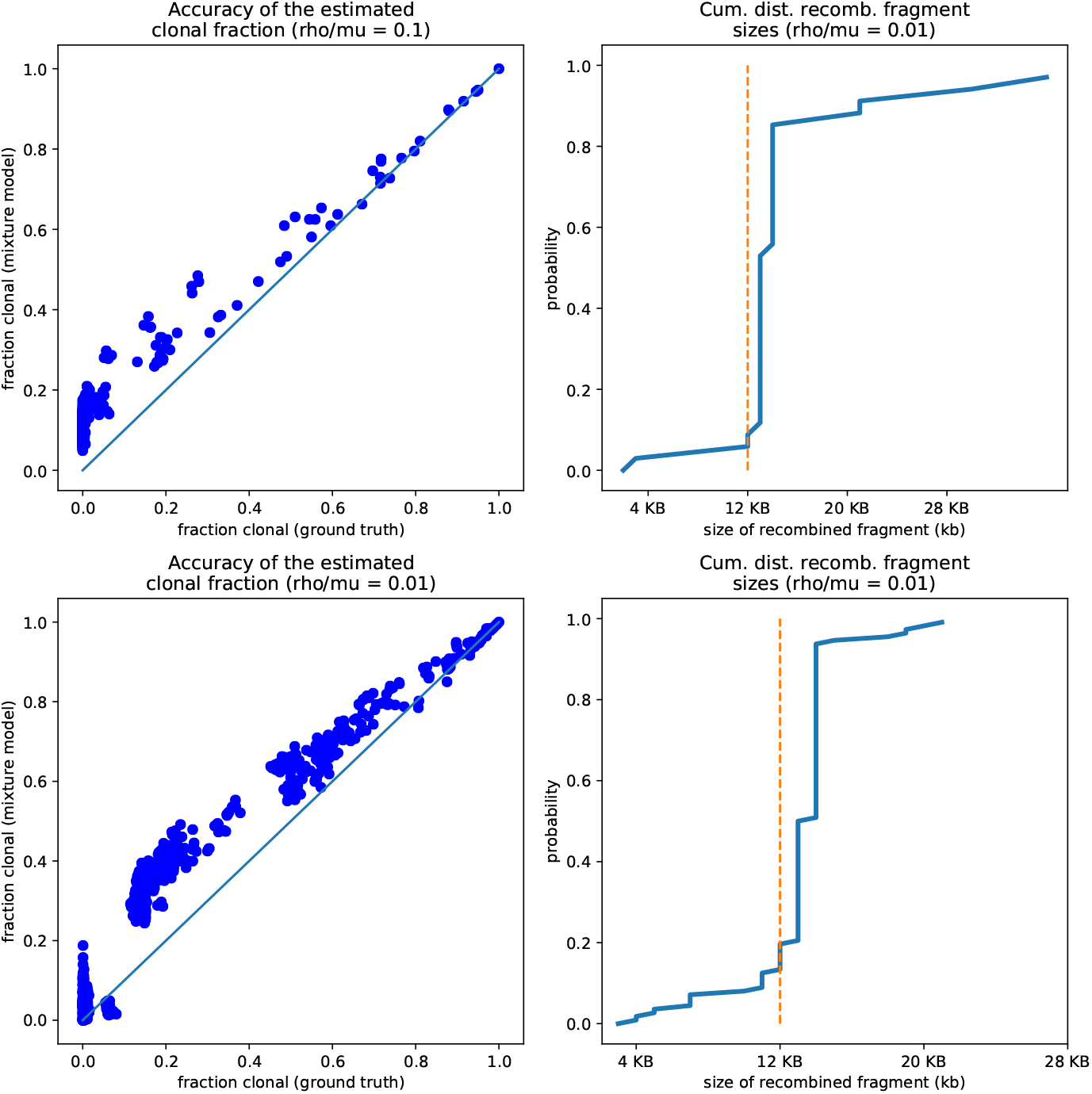
The pairwise analysis accurately estimates both the clonally inherited fractions and the sizes of the recombined segments. **Left panels:** Comparison of the true fraction of the genome that was clonally inherited (horizontal axis) and the estimated fraction from the pairwise analysis (vertical axis) for the simulation with *ρ/μ* = 0.1 (top) and 0.01 (bottom). Each dot corresponds to one pair and the diagonal line shows the line *y* = x. **Right panels:** Cumulative distribution of the lengths of the recombined segments, as estimated by the pairwise analysis of the simulations with *ρ/μ* = 0.1 (top) and 0.01 (bottom). The vertical dotted lines show the ground truth, i.e. all recombined segments were 12kb long in the simulations.

**Figure S6.**
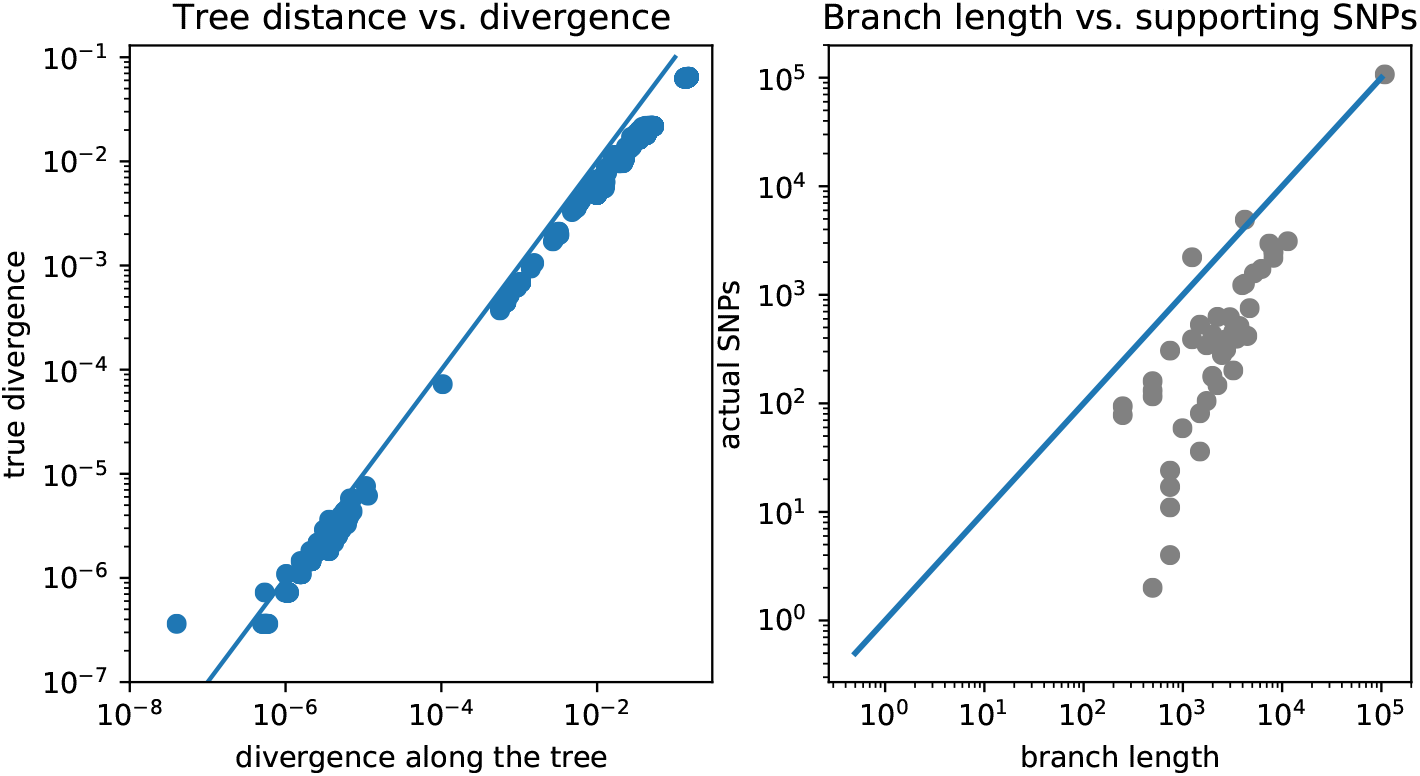
Comparison of pairwise distances and number of SNPs on each branch as predicted by the core tree, with pairwise distances and SNP numbers observed in the data. **Left panel**: Scatter plot of the pairwise divergences along the core tree (horizontal axis) with the observed pairwise divergences. **Right panel**: For each branch of the tree, the total number of SNPs predicted to fall on the branch, i.e. the probability of a substitution to occur along the branch times the length of the alignment, is shown (horizontal axis) against the actual number of SNPs that fall on the branch that are observed in the data. All axes are shown on logarithmic scales.

**Figure S7.**
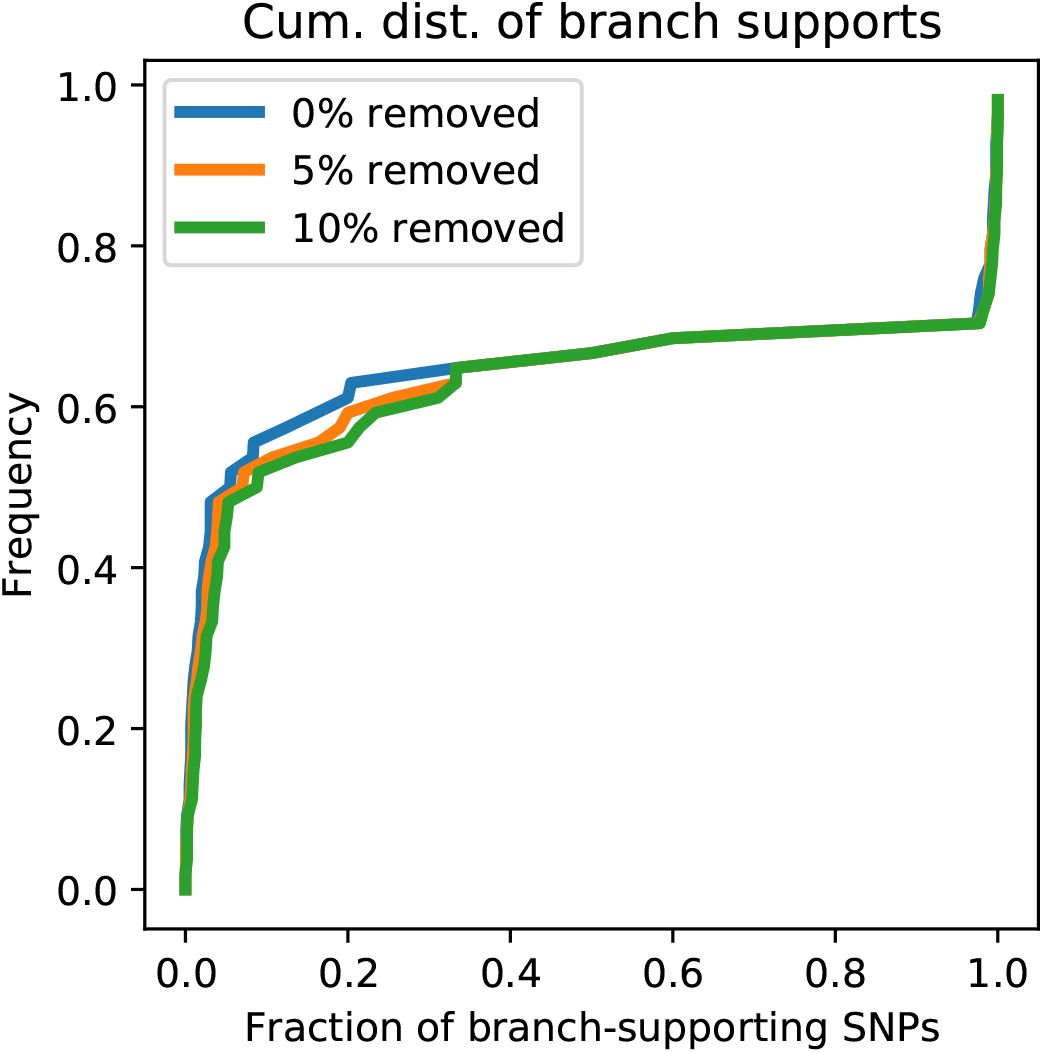
Cumulative distribution of the fraction of supporting SNPs across all branches of the core tree for the original alignment of *E. coli* strains (blue) as well as for alignment from which 5% (orange) and 10% of potentially homoplasic columns were removed. The distribution of SNP support is highly insensitive to small fraction of homoplasies among the SNPs.

**Figure S8.**
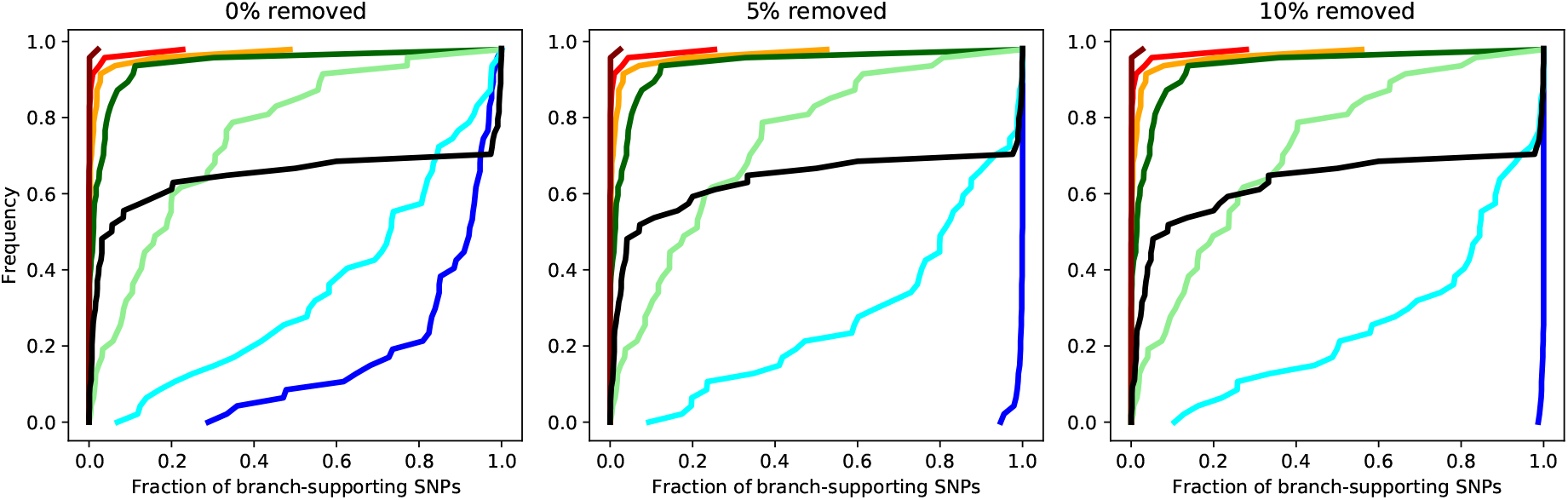
Cumulative distributions of the fraction of supporting SNPs across all branches of the core tree for the alignments resulting from the simulations with *ρ/μ* = 0 (dark blue), *ρ/μ* = 0.001 (light blue), *ρ/μ* = 0.01 (light green), *ρ/μ* = 0.1 (dark green), *ρ/μ* = 0.3 (orange), *ρ/μ* = 1 (red), and *ρ/μ* = 10 (dark red). The panels corresponds to the distribution for the original alignment (left), and alignments from which 5% (middle) and 10% of potentially homoplasic columns were removed. For reference, the distribution of support for the *E. coli* data is shown in black. Note that except for the simulations without any recombination where all inconsistencies are caused by homoplasies, the distributions are generally insensitive to the removal of homoplasic positions.

**Figure S9.**
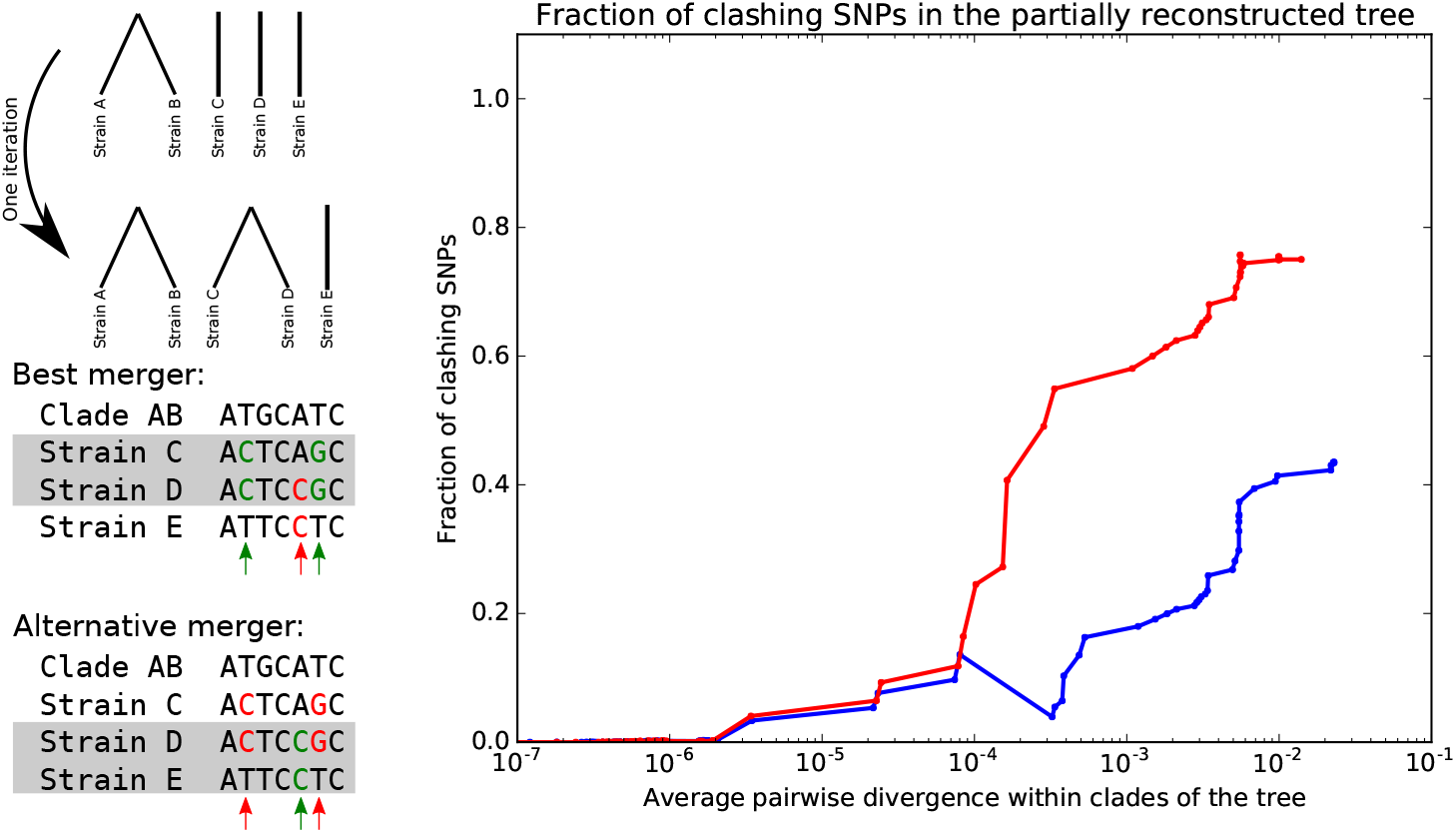
Bottom-up tree building, minimizing SNP clashes. **Left panel**: Illustration of the iterative bottom-up tree reconstruction. At each step the pair of clades is fused that minimizes the number of SNPs that clash with the fusion (red arrows). In case of multiple pairs that have the same number of clashing SNPs, the pair with the largest number of supporting SNPs (green arrows) is chosen. **Right panel**: Fraction of SNPs that support vs. clash with the partially reconstructed tree as a function of the average pairwise divergence of strains that occur within the clades of the partially reconstructed tree. The blue curve corresponds to the full set of strains and the red curve to all strains except for the outgroup. The horizontal axis is shown on a logarithmic scale.

**Figure S10.**
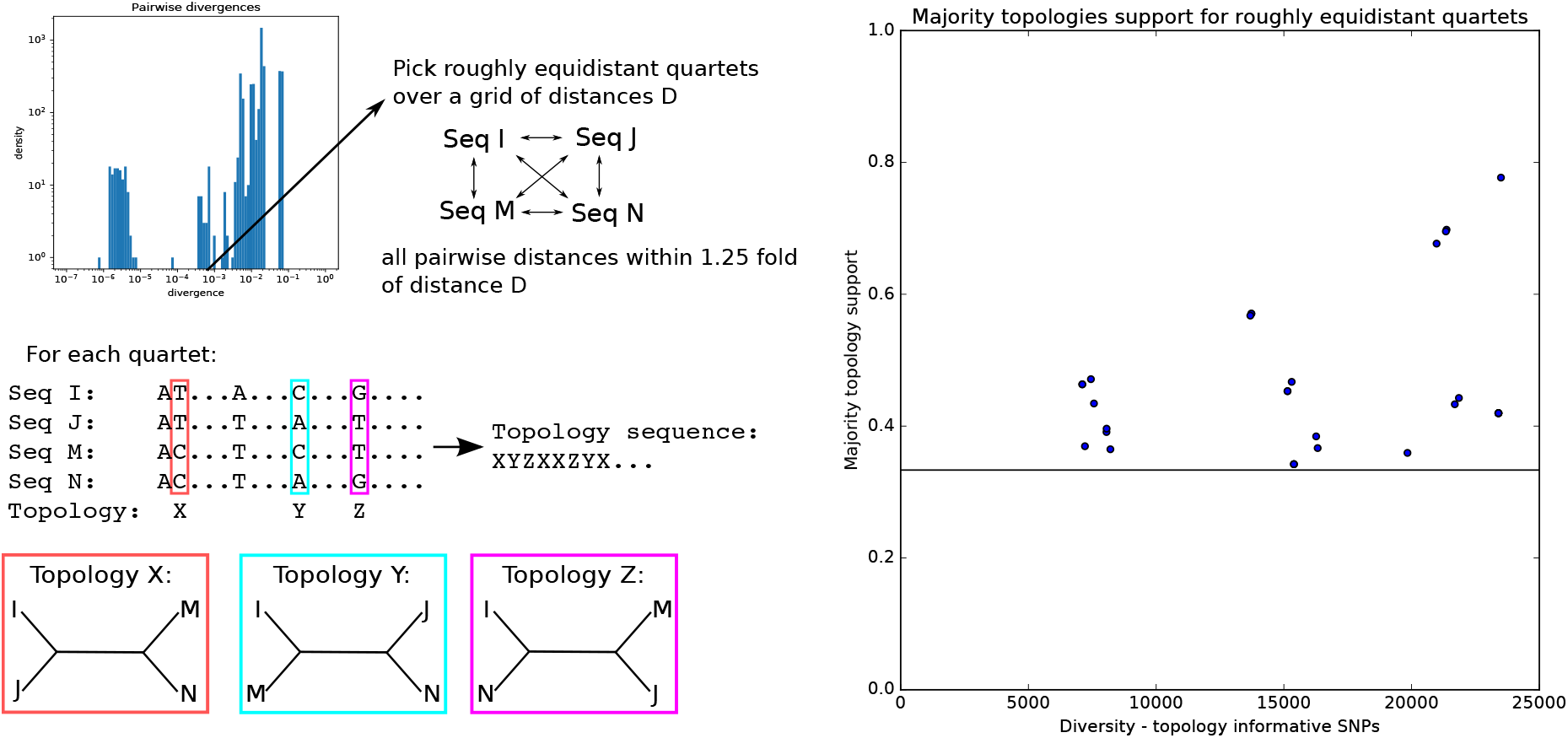
Quartets of roughly equidistant strains have no consensus phylogeny. **Left**: Using the distribution of pairwise distances (top panel) we select, for each pairwise distance *D*, quartets of strains whose pairwise distances are all within a factor 1.25 of D. SNPs for which two strains have one letter and two strains another are informative for the topology and each support one of the three possible topologies. **Right**: For each quartet we determined the topology that is supported by most SNPs and then calculated the fraction of topology-informative SNPs that supported the most common topology. The plot shows the fraction of SNPs supporting the most common topology (vertical axis) as a function of the total number of informative SNPs (horizontal axis). The horizontal line marks the minimal possible fraction, which is attained when all 3 topologies are supported by 1/3 of the SNPs. Note that, for the majority of quartets, the most common topology is supported by less than half of the informative SNPs.

**Figure S11.**
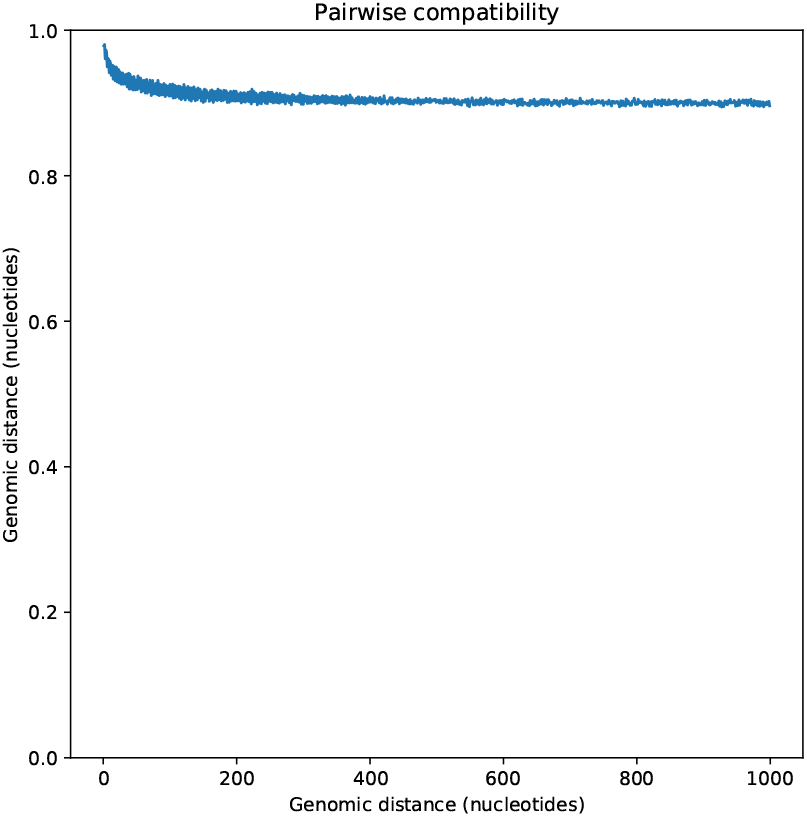
Pairwise SNP compatibility as a function of genomic distance. The plot show the fraction of SNP pairs that are compatible with a common phylogeny as a function of the genomic distance between the pair (in nucleotides).

**Figure S12.**
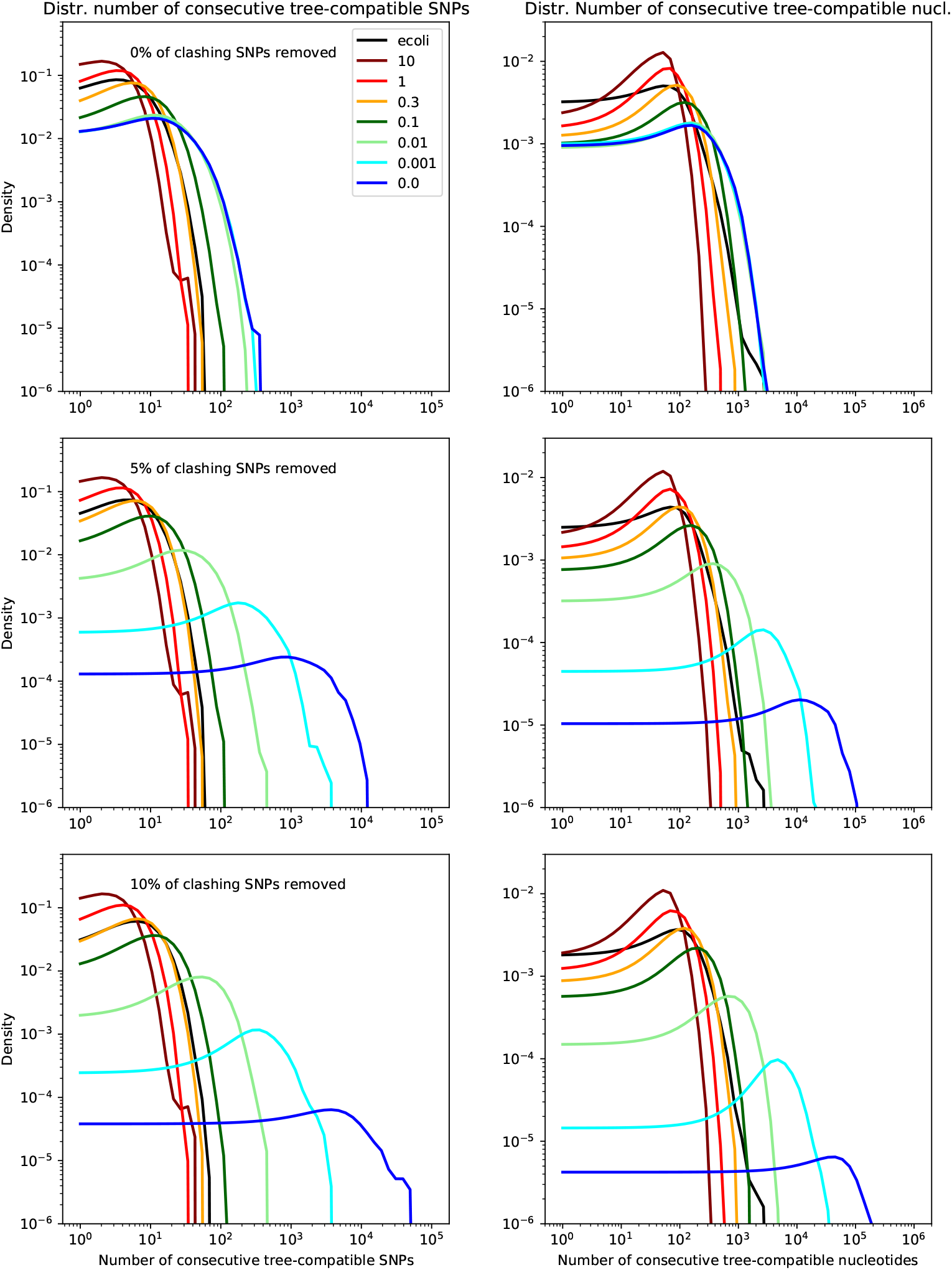
Probability distributions of the length of tree-compatible segments along the alignments of the *E. coli* genomes (black line) and the alignments of the sequences from the simulations with *ρ/μ* = 0 (dark blue), *ρ/μ* = 0.001 (light blue), *ρ/μ* = 0.01 (light green), *ρ/μ* = 0.1 (dark green), *ρ/μ* = 0.3 (orange), *ρ/μ* = 1 (red), and *ρ/μ* = 10 (dark red). The left panels show the distributions of the number of consecutive SNP columns and the right panels the distributions of the number of consecutive nucleotides. The top row corresponds to the full alignments, and the middle and bottoms row to the alignments from which 5% and 10% of potentially homoplasic positions have been removed, respectively.

**Figure S13.**
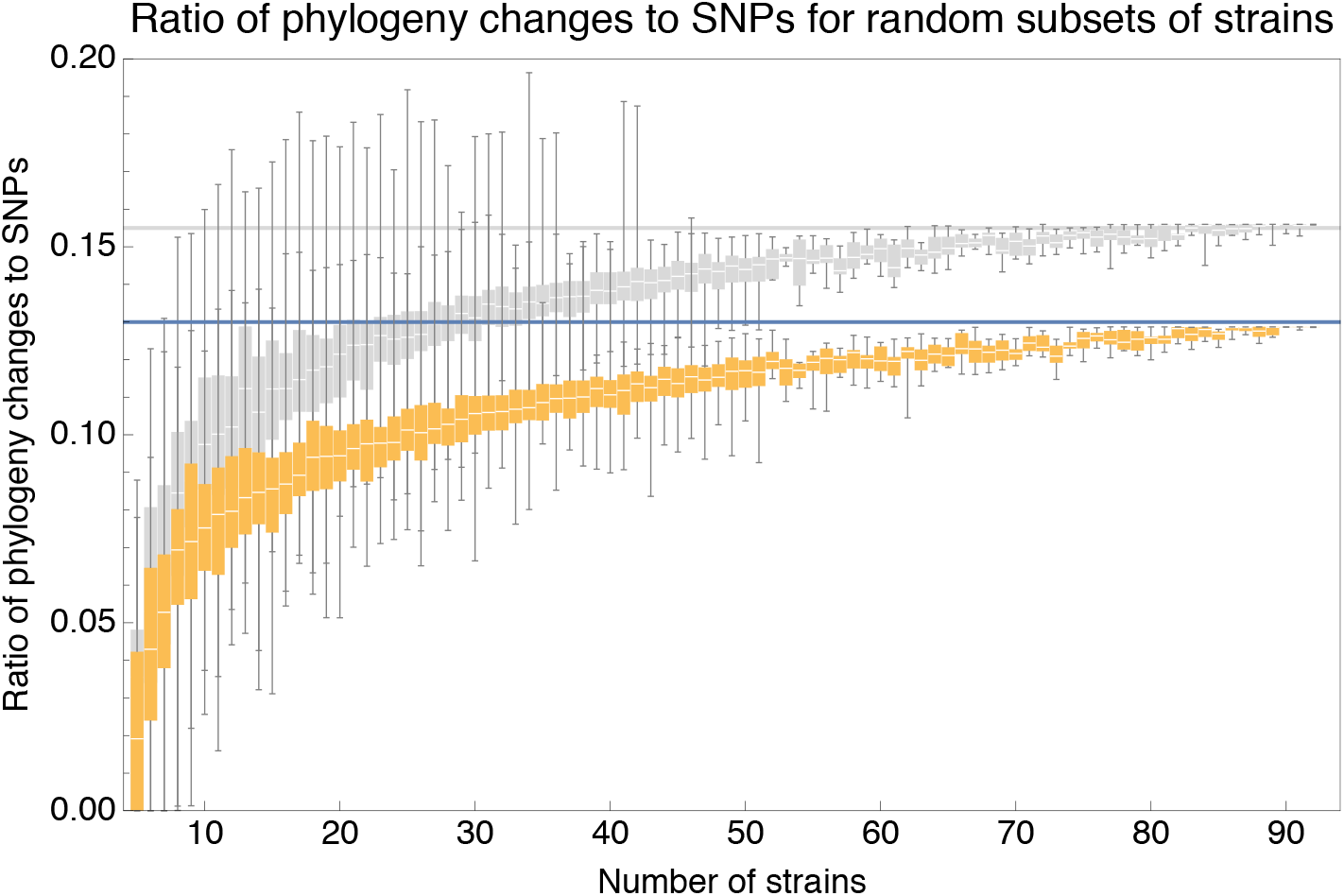
Ratio *C/M* of the minimal number of phylogeny changes *C* to mutations *M* for random subsets of strains using the alignment for which 5% of potentially homoplasic positions have been removed (orange) as well as for the full alignment (grey). For strain numbers ranging from *n* = 4 to *n* = 92, we collected random subsets of *n* strains and calculated the ratios *C/M* of phylogeny changes to SNPs in the alignment. The figure shows box-whisker plots that indicate, for each strain number *n*, the 5th percentile, first quartile, median, third quartile, and 95th percentile of the distribution of *C/M* across subsets. The blue line shows *C/M* = 0.13 and the grey line *C/M* = 0.155.

**Figure S14.**
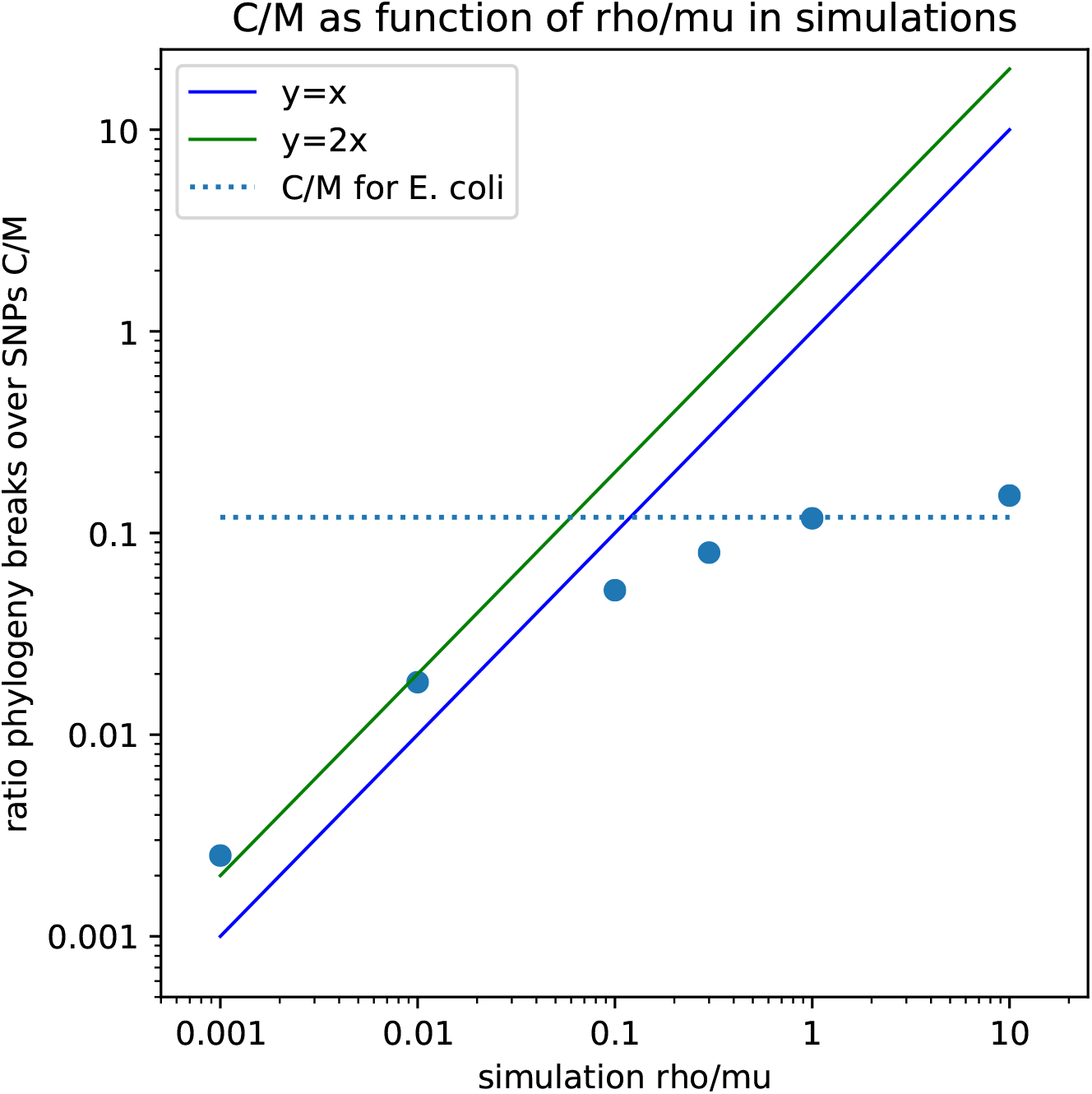
Observed ratio *C/M* of the minimal number of phylogeny changes *C* and SNPs *M* in the alignment (vertical axis) with the ratio of recombination and mutation rate *ρ/μ* used in the simulation (horizontal axis), as calculated on the simulation data using the alignments from which 5% of potentially homoplasic sites have been removed. Each blue dot corresponds to a simulation with a given value of *ρ/μ.* Both axes are shown on logarithmic scale. For guidance, the lines *y* = *x* (blue) and *y* = *2x* (green) are shown. The horizontal dashed line shows the ratio *C/M* = 0.13 that is observed for the 5% homoplasy corrected alignment of the *E. coli* strains.

**Figure S15.**
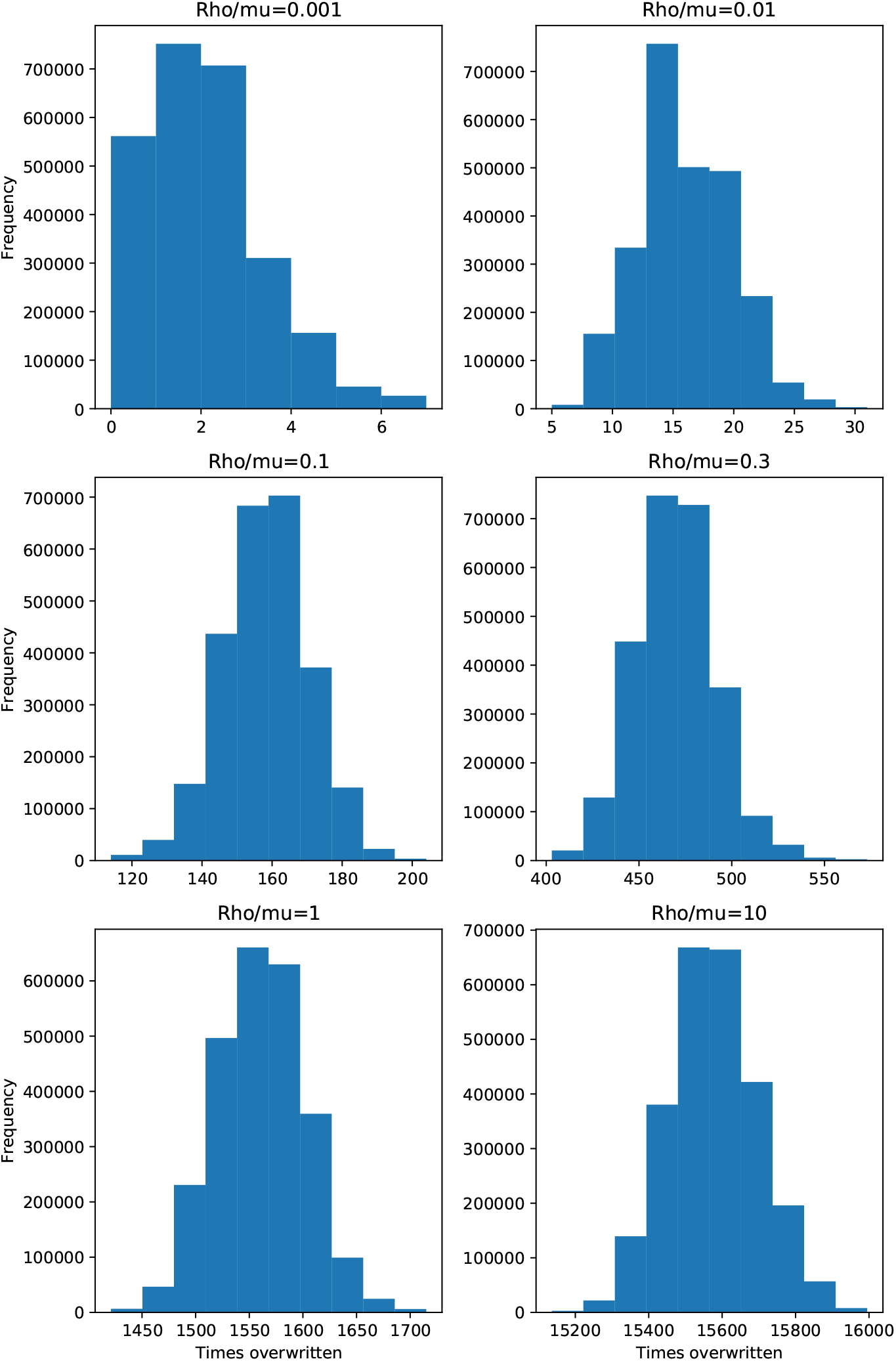
Histograms of the number of times each position in the genome was overwritten by recombination for the simulations with recombination-to-mutation ratios ranging from *ρ/μ* = 0.001 (top left), to *ρ/μ* = 10 (bottom right).

**Figure S16.**
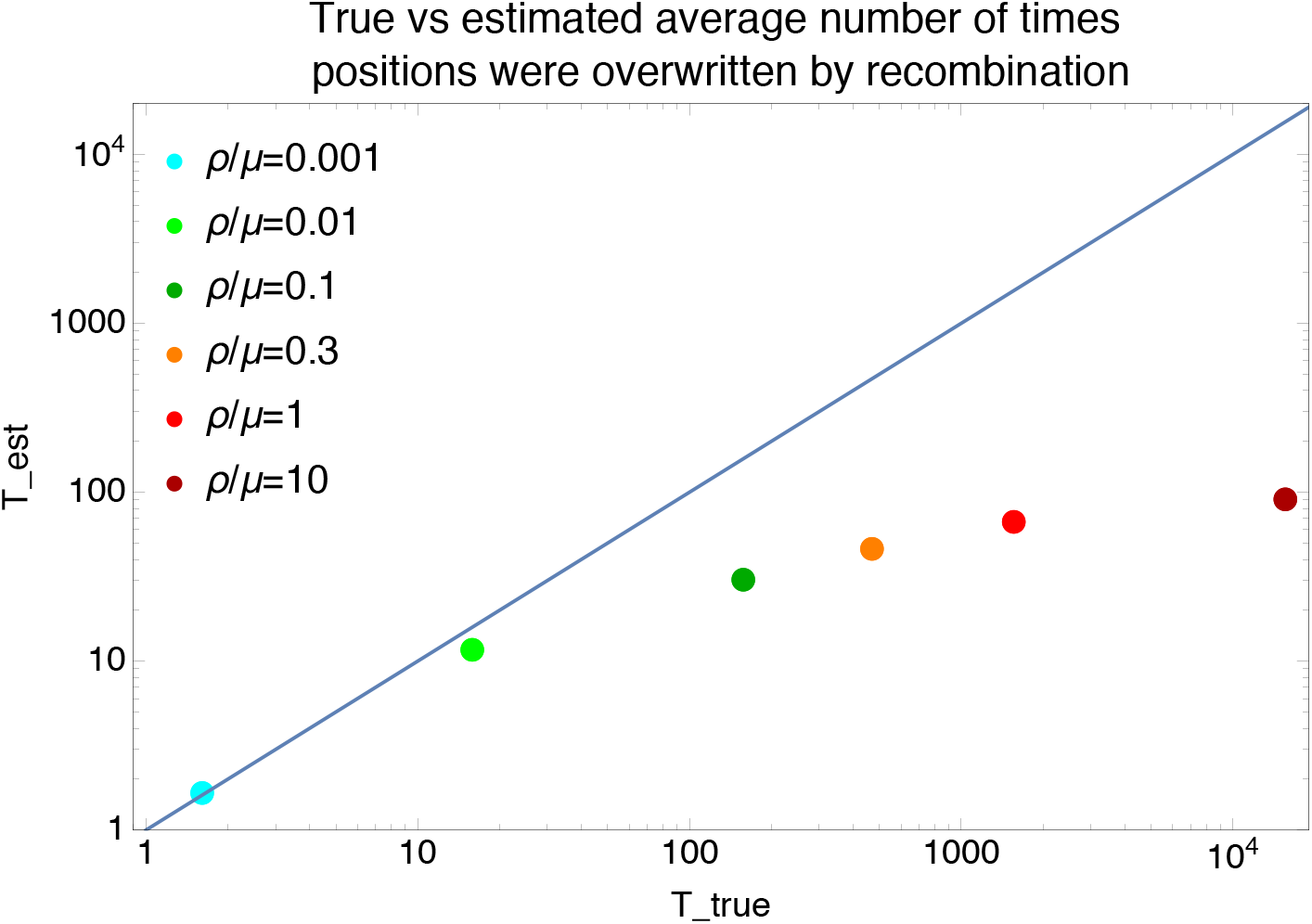
Comparison of the true average number of times *T_true_* that positions were overwritten by recombination (horizontal axis) versus the estimated number of times *T_est_* = *L_r_C*/(2*L*), as estimated from the lower bound on the number of phylogeny changes (with *L* the length of the alignment and *L_r_* the average length of the recombination segments), for the simulation data. Each dot corresponds to simulation data with different recombination to mutation rates *ρ/μ* (see legend). The diagonal line shows the line *T_est_ T_true_*.

**Figure S17.**
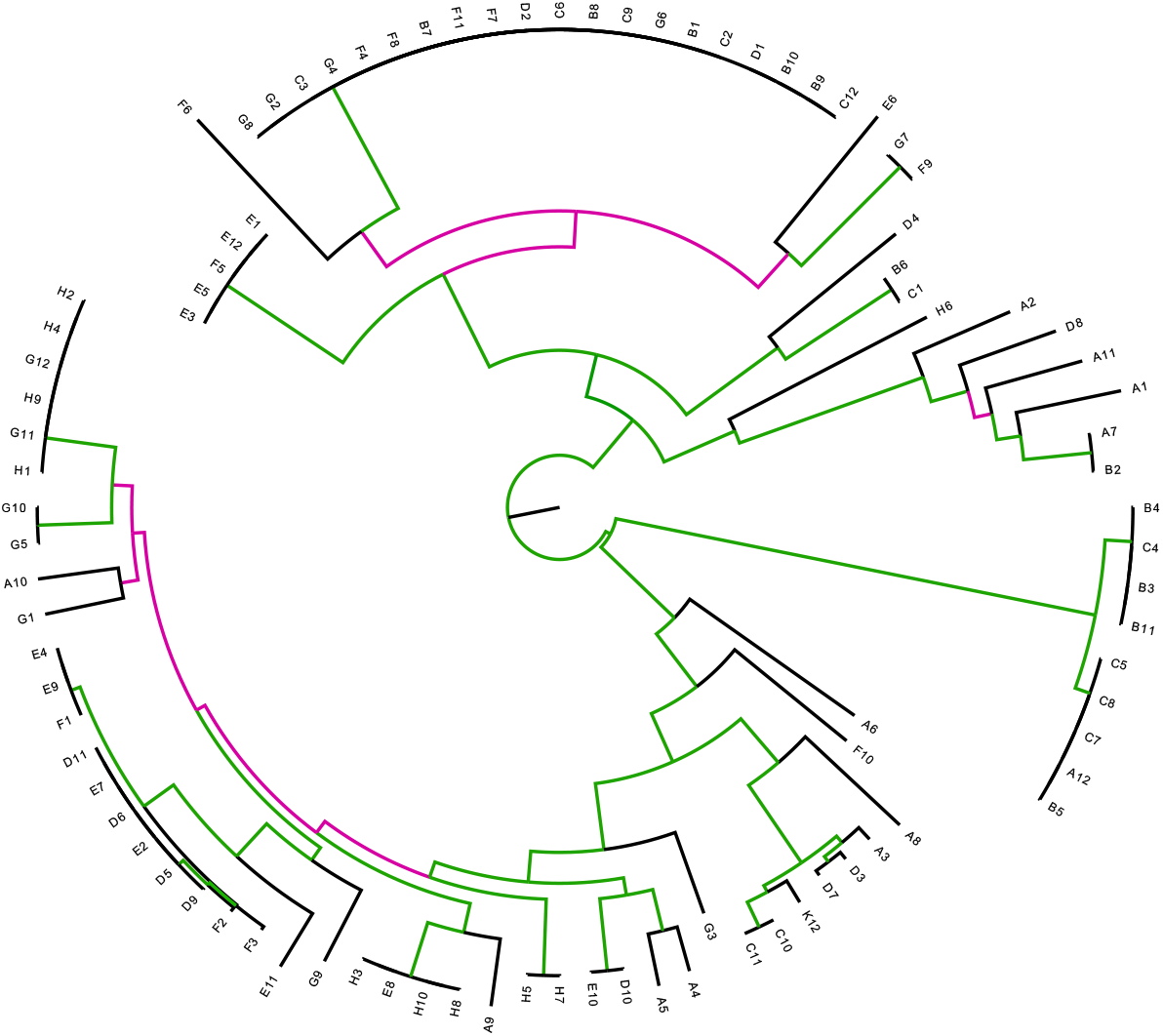
Differences between the core tree *T* and the tree *T*’ reconstructed from the alignment from which all SNPs that fall on branches of the core tree have been removed. Each branch of the core tree *T* is colored green when the branch also occurs in *T*’ and pink if it does not. The Robinson-Foulds distance between two trees is defined as the number of branches (i.e. bi-partitions) that occur in only one of the two trees. For *T* and *T*’ the Robinson-Foulds distance is 62 out of a maximal 178, i.e. a fraction 0.35 does not match. Note also that for tree *T*’ only 3% of the SNPs fall on its branches.

**Figure S18.**
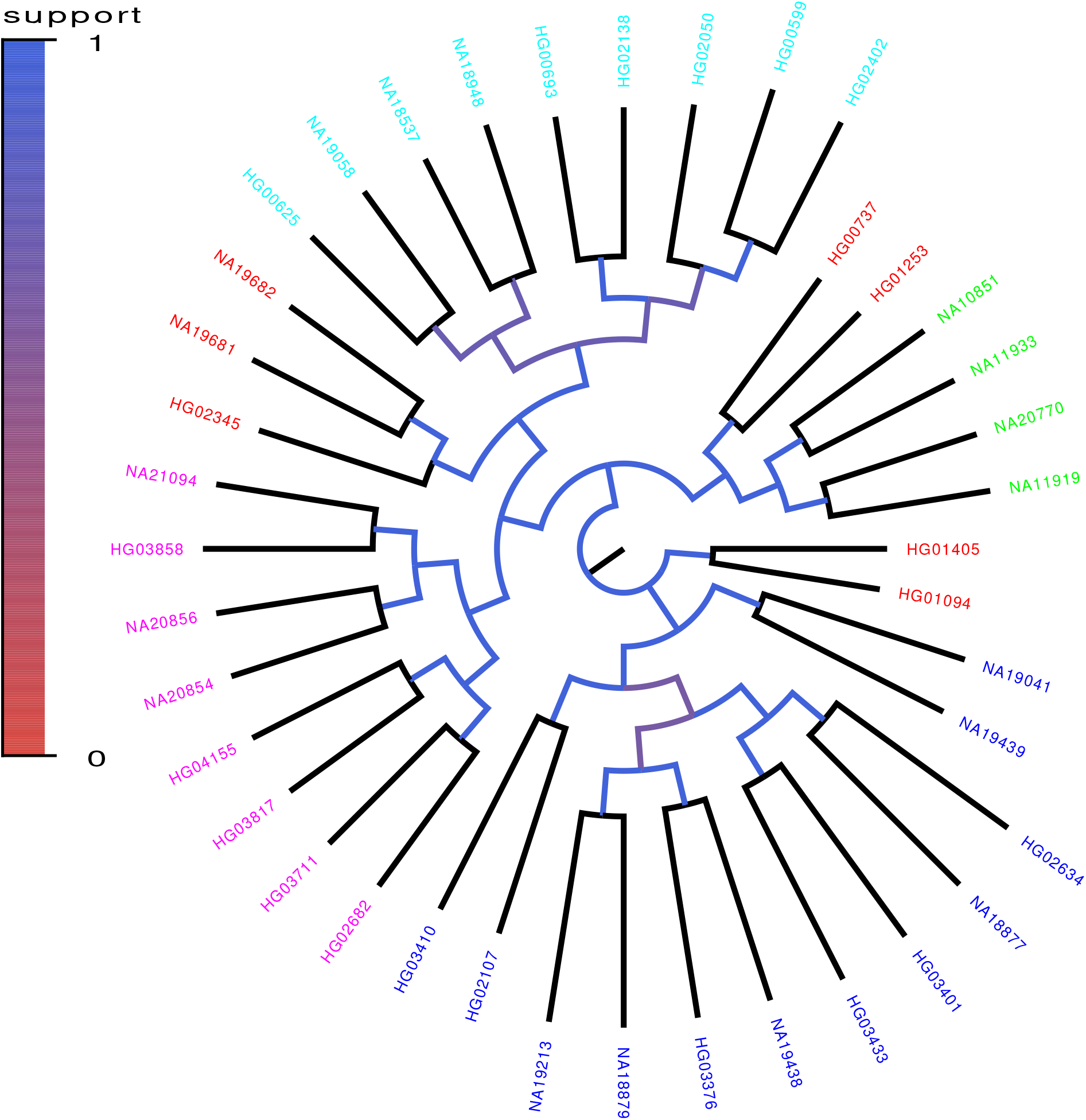
Core tree build by PhyML from the sequences of chromosomes 1 — 12 of 40 randomly chosen genomes of the 1000 Genome project. The colors and numbers indicate what fraction of the time each split in the core tree occurred in trees build from a random subsets of half of the genomic loci. The colors on the leafs indicate the annotated ancestry of the individuals, with blue corresponding to African ancestry, green to European ancestry, red to (South) American ancestry, cyan to East Asian ancestry, and magenta to South Asian ancestry. Individuals from the same geographic area reliably form clades in the tree with the exception of South Americans, two of which form an outgroup of the Europeans, three an outgroup of East Asians, and two an outgroup of Africans.

**Figure S19.**
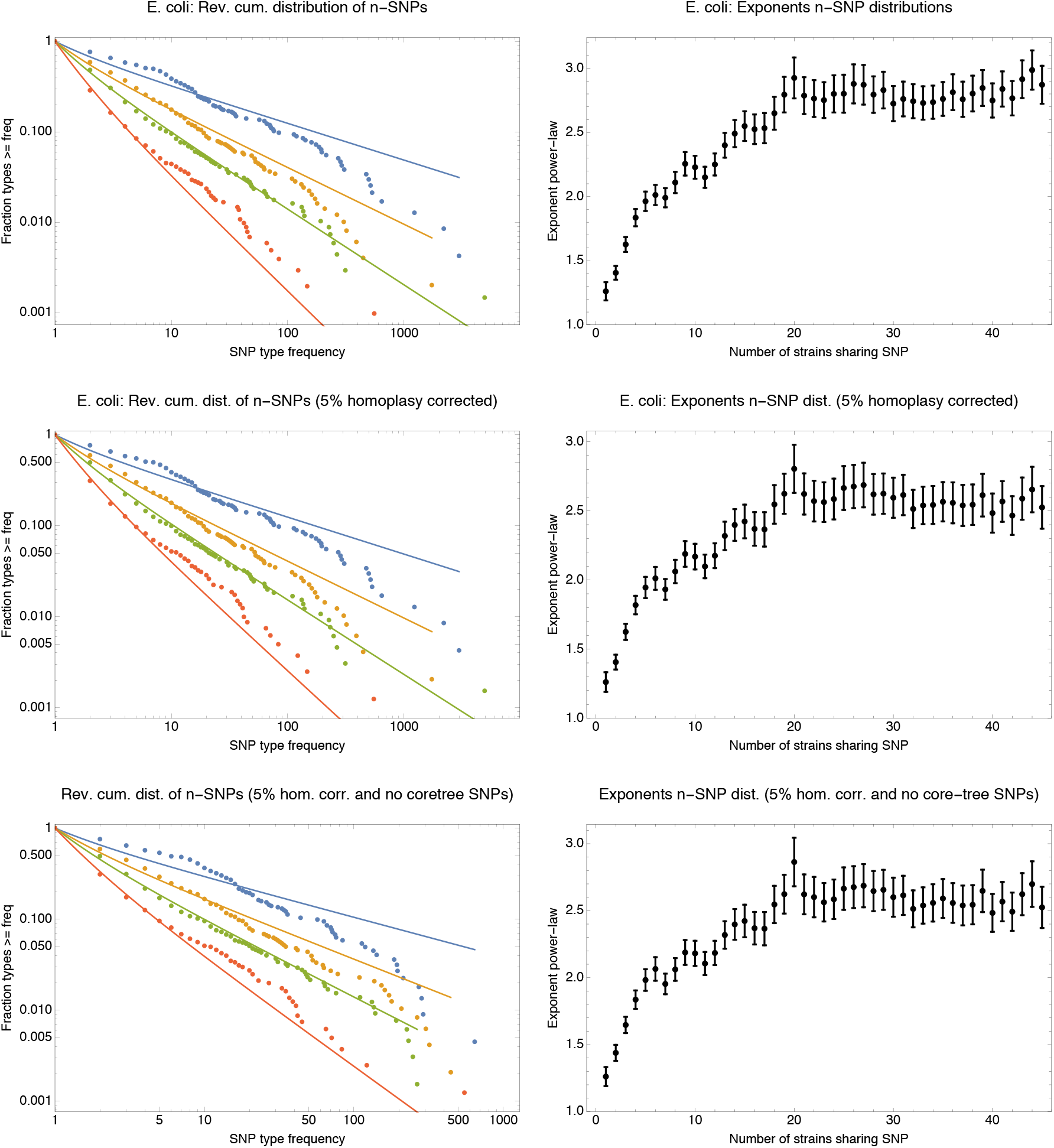
Distributions of *n*-SNP frequencies (left panels) and exponents of the power-law fits (right panels) for original *E. coli* core genome alignment (top row), the core genome alignment from which all *n*-SNPs that correspond to branches of the core tree have been removed (middle row), and the 5% homoplasy-corrected core genome alignment (bottom row). The observed *n*-SNP distributions and exponents are very similar for all three datasets.

**Figure S20.**
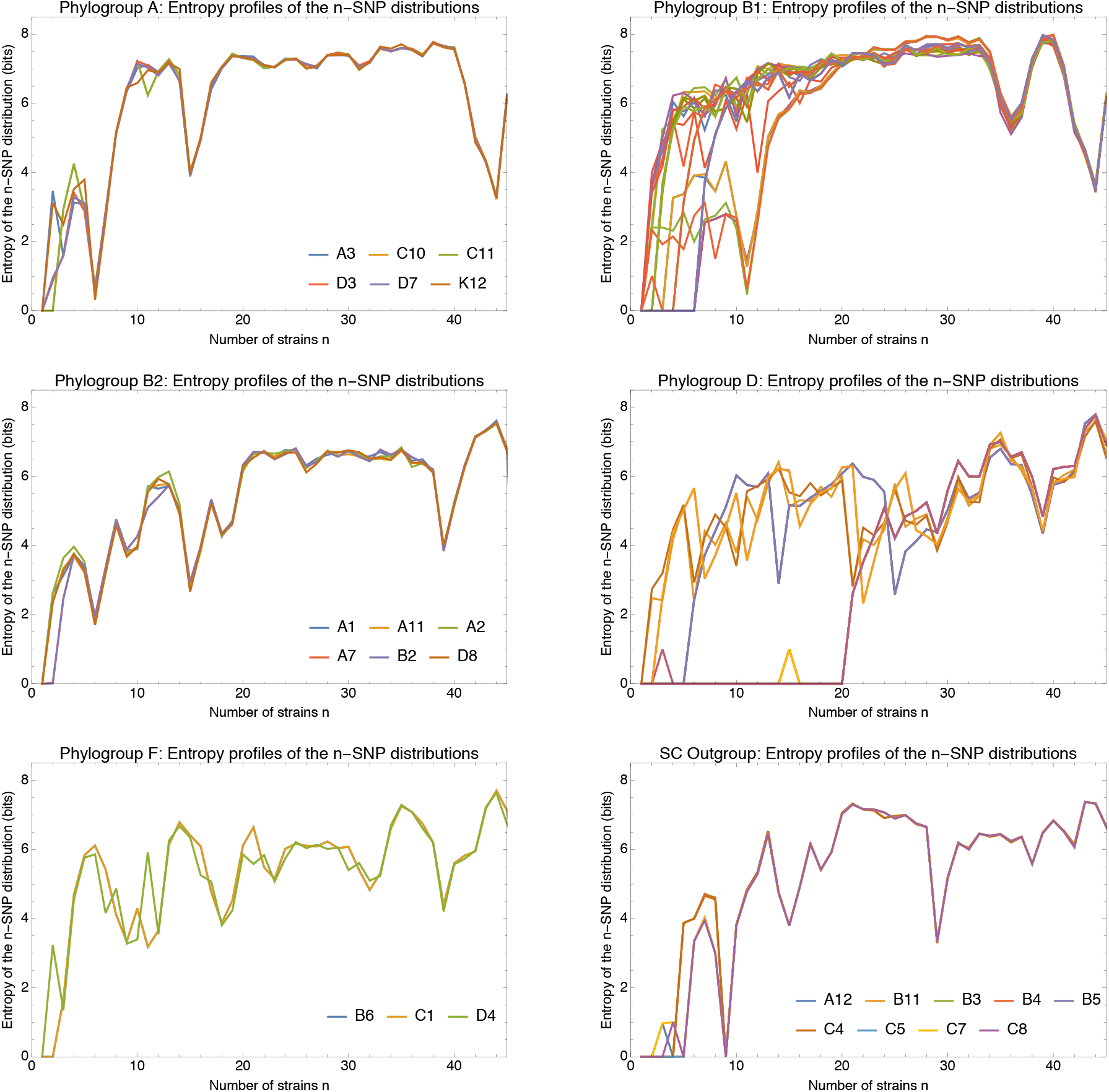
Entropy profiles of the *n*-SNP distributions for each of the *E. coli* phylogroups. Each panel shows the entropy *H_n_*(*s*) of the *n*-SNP distribution (vertical axis) as a function of *n* for each strain *s* (differently colored lines) within a phylogroup (indicated at the top of the panel). All entropies were calculated on the 5% homoplasy-corrected alignments. For phylogroups with less than 10 strains, the names of the strains are indicated in the legend of the plot. Note that the bottom right panel corresponds to the outgroup of 9 strains that are highly diverged from all other strains. Note also that entropy profiles are perfectly overlapping for strains within groups that are so close that they had a recent clonal ancestor. For example, for the pairs (C10,C11) and (D3,D7) in phylogroup A, the pair (A7,B2) in phylogroup B2, and pair (B6,C1) in phylogroup F, only the two profiles are identical and only one curve is visible.

**Figure S21.**
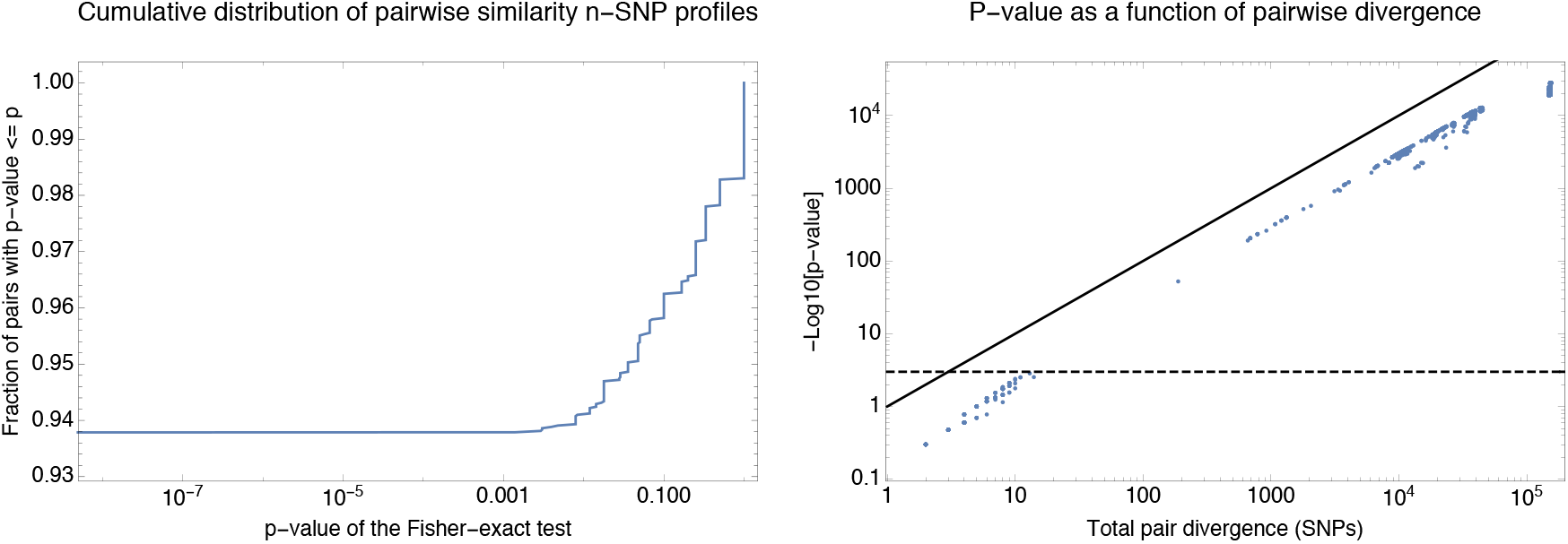
Statistical significance of the difference in *n*-SNP statistics of all pairs of strains. **Left panel**: Cumulative distribution of the p-values of the Fisher exact test (Materials and Methods) for the *n*-SNP statistics across all pairs of strains. About 94% of all pairs have significantly different *n*-SNP distributions. **Right panel**: p-values of the Fisher-exact test as a function of the divergence of each pair of strains. Each dot corresponds to one pair with the horizontal axis showing the total number of single-nucleotide differences between the pair of core genomes and the vertical axis showing minus the logarithm of the p-value (base 10). The diagonal line corresponds to the line *y* = *x* and the horizontal dashed line corresponds to a *p*-value of 0.001. Note that the significance correlates very well with the divergence of the pair and that only pairs of genomes that differ in less than 10 nucleotides genome-wide have indistinguishable *n*-SNP statistics.

**Figure S22.**
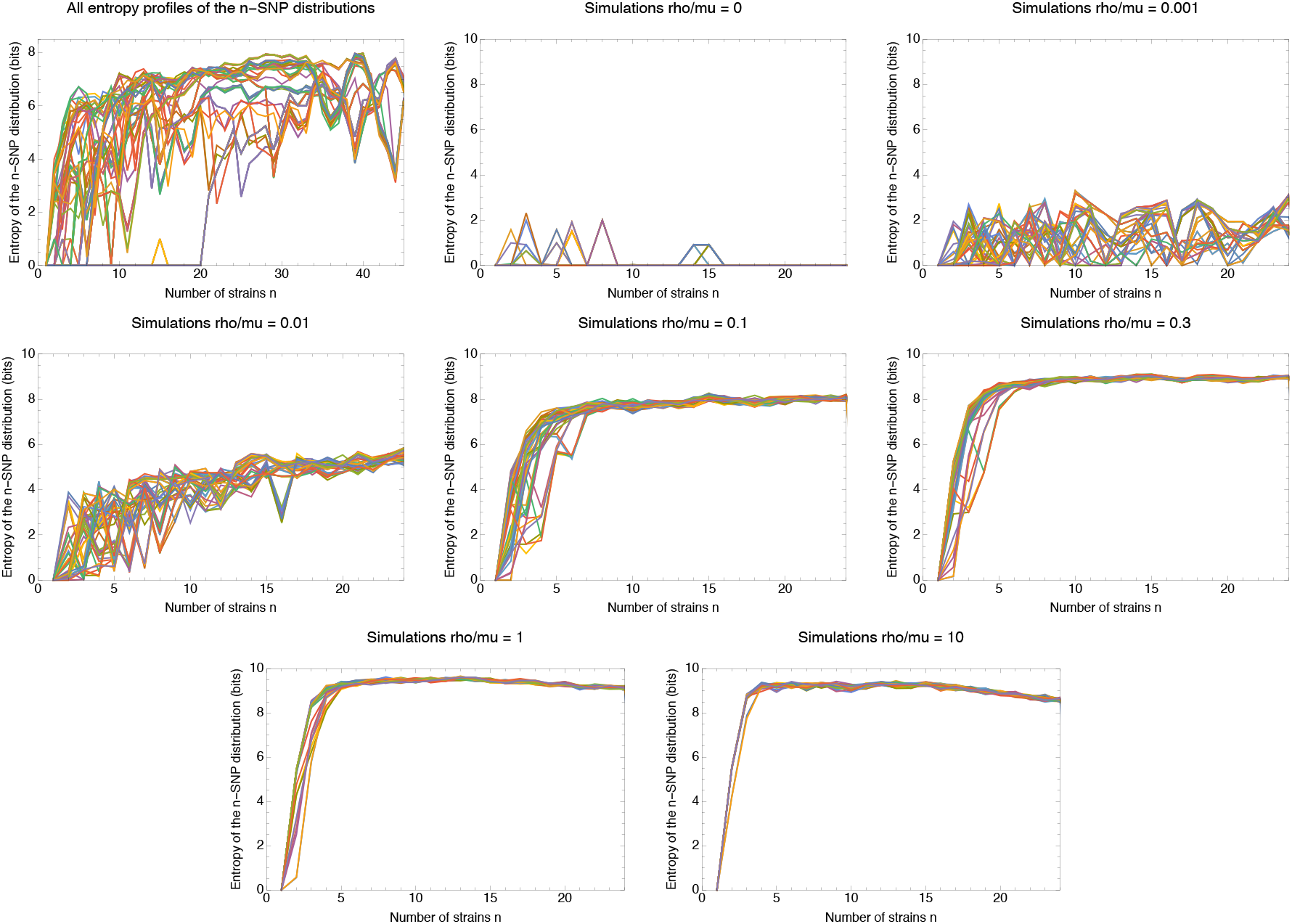
Entropy profiles of the *n*-SNP distributions for the *E. coli* data (top left panel) and the data from the simulations (other panels, with the recombination rate *ρ/μ* indicated in the title of each panel. Each panel shows the entropy *H_n_*(*s*) of the *n*-SNP distribution (vertical axis) as a function of *n* for each strain *s* (differently colored lines). All entropies were calculated on the 5% homoplasy-corrected alignments. Note that the small number of nonzero entropies for the simulations without recombination are due to the small number of remaining homoplasies.

**Figure S23.**
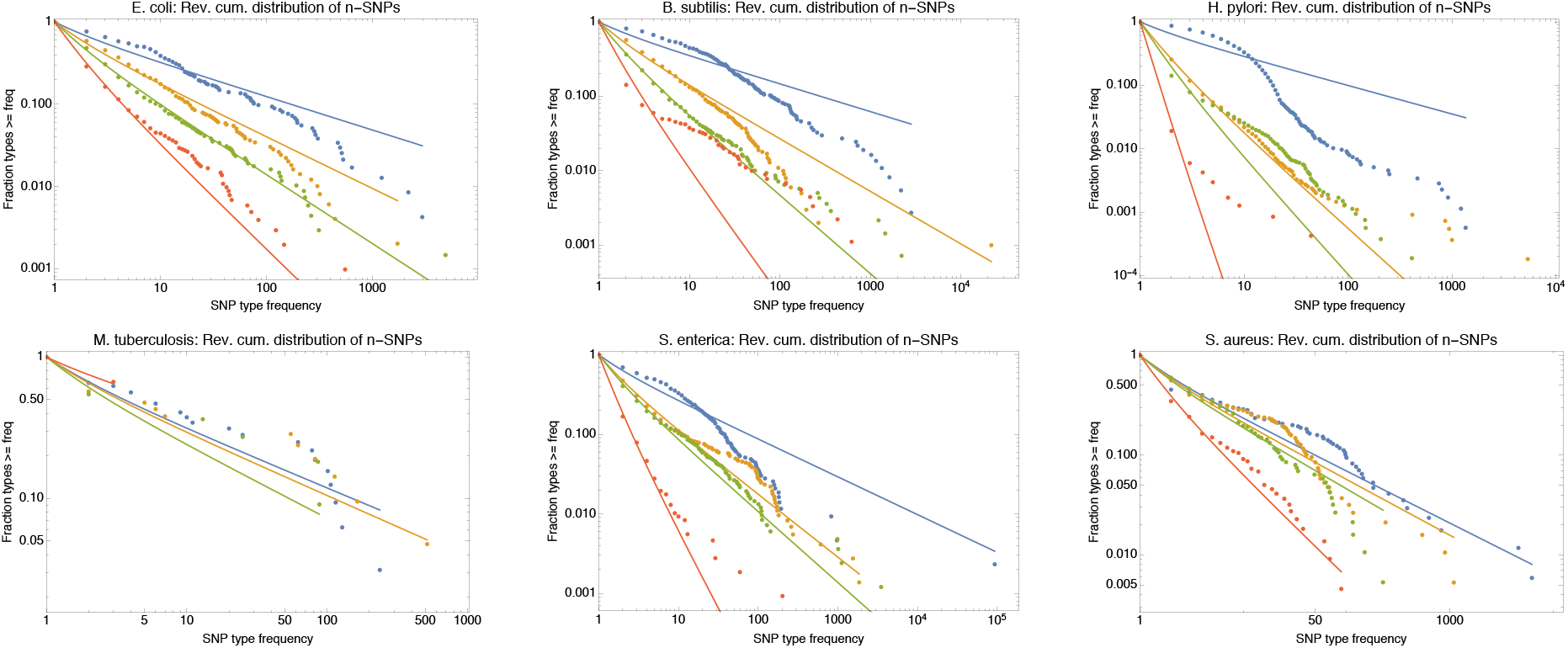
Power-law fits of the *n*-SNP distributions for all 6 species. Each panel shows the reverse cumulative distributions of the frequencies of all observed 2-SNPs (blue dots), 3-SNPs (orange dots), 4-SNPs (green dots), and 12-SNPs (red dots), with the solid lines in corresponding colors showing power-law fits. The species is indicated at the top of each panel.

**Figure S24.**
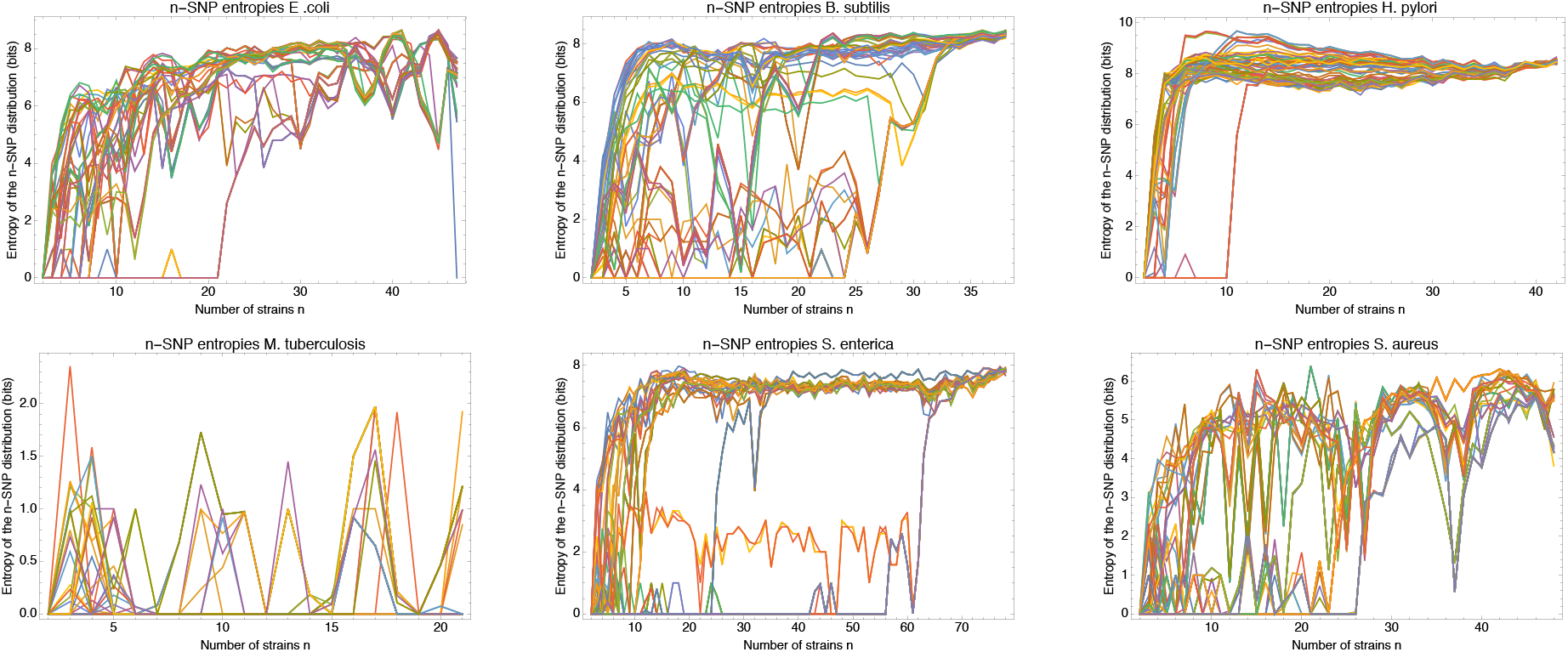
Entropy profiles of all the strains for each of the 6 species. Each panel corresponds to one species (indicated at the top) and shows the entropy profiles *H_s_*(*n*) of the distribution of *n*-SNPs in which a particular strain occurs as a function of the number of strains *n* for each strain *s* (different colors).

**Figure S25.**
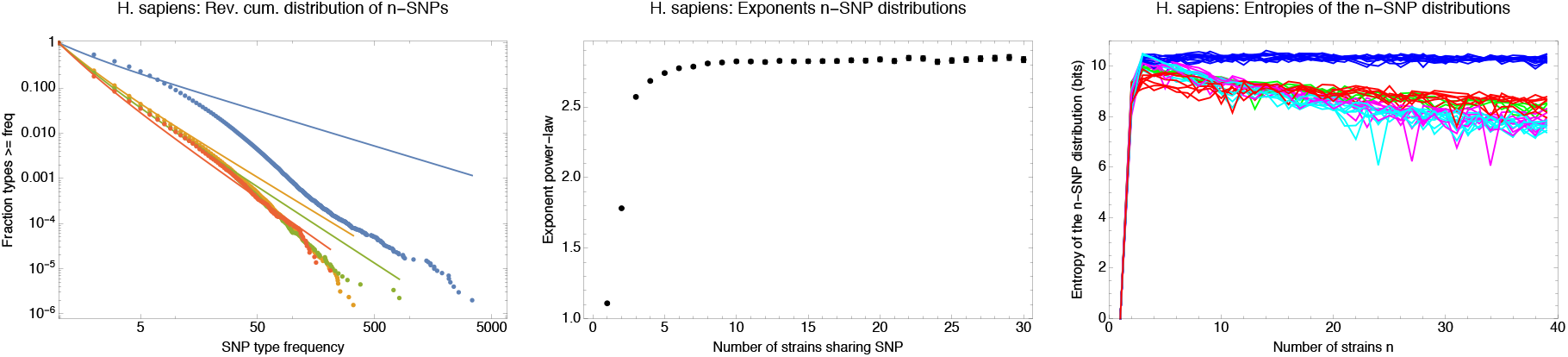
Human *n*-SNPs distributions and entropy profiles. **Left panel**: Reverse cumulative distributions of the frequencies of all observed 2-SNPs (blue dots), 3-SNPs (orange dots), 4-SNPs (green dots), and 12-SNPs (red dots), with the solid lines in corresponding colors showing power-law fits, for the human data. Both axes are shown on a logarithmic scale. **Middle panel**: Fitted exponents for the power-law *n*-SNP distributions on the human data for *n* ranging from 1 to 30. The error bars correspond to 95% posterior probability intervals. **Right panel**: Entropy profiles of the *n*-SNP distributions for the individuals from the 1000 Genomes project. Individuals with African ancestry are in blue, European in green, South American in red, East Asian in cyan, and South Asian in magenta.

**Figure S26.**
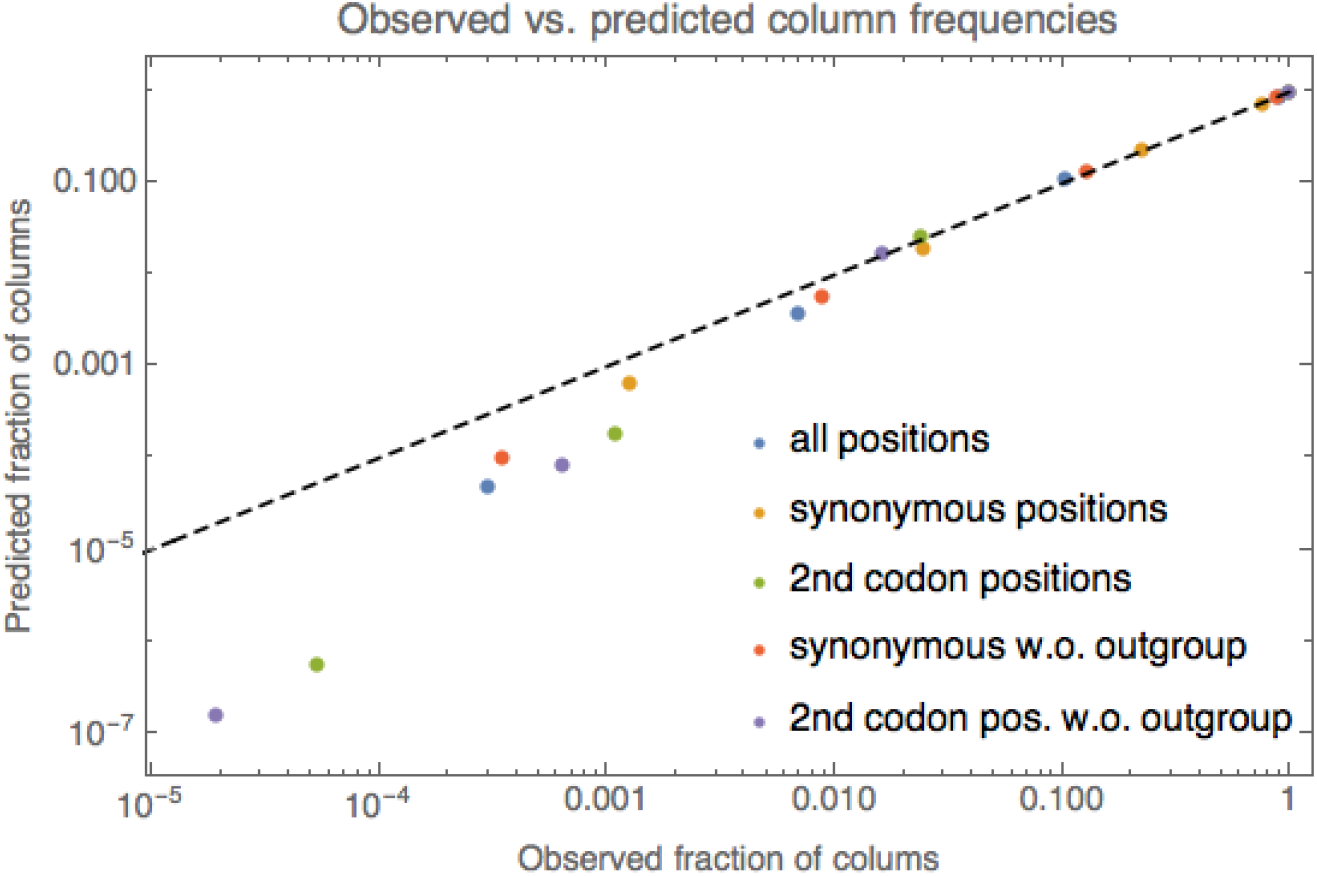
Comparison of the observed frequencies of columns with 1, 2, 3, and 4 different nucleotides under the simple model described in the methods. Different colored dots correspond to different subsets of columns, as indicated in the legend. For each color, 4 dots are shown corresponding to the observed frequencies of columns with 1, 2, 3, and 4 nucleotides (horizontal axis) and the predicted frequencies according to the simple model (vertical axis). The dashed line shows the identity *y* = *x*. Both axes are shown on logarithmic scales.

**Table S1.**
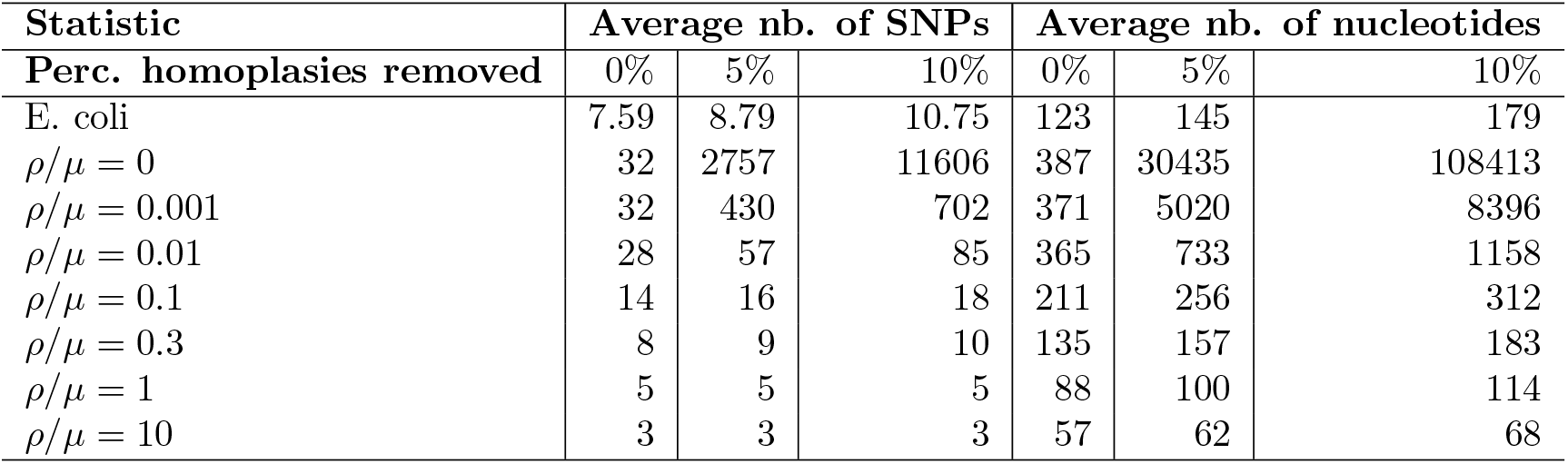
Average length of tree compatible segments, both in terms of number of consecutive SNPs and number of consecutive nucleotides, for the full *E. coli* and simulation data, as well as for alignments from which 5% or 10% of potentially homoplasic sites were removed.

**Table S2.** Number of substitutions and phylogeny changes within sub-alignments corresponding to known phylogroups. Starting from the full 5% homoplasy-corrected core genome alignment we extracted, for each phylogroup, the sub-alignment of all strains belonging to that phylogroup and counted the number of bi-allelic SNPs *M* and number of phylogeny changes *C*. Each row of the table corresponds to one of the known phylogroups and shows the number of our strains in that phylogroup, the SNP rate within the clade *M/L*, the lower bound *C* on the number of phylogeny changes, the ratio *C/M*, and the estimated average number of times each position in the genome has been overwritten by recombination Test. Note that the phylogroup ‘O’ stands for the outgroup. See Suppl. Fig. S1 for the phylogroup annotation of our strains.

| Phylogroup | Nb. of strains | SNP rate $M/L$ | Phyl. changes $C$ | $C/M$ | $T_{\text{est}}$ |
| --- | --- | --- | --- | --- | --- |
| A | 6 | 0.0024 | 178 | 0.027 | 0.65 |
| B1 | 35 | 0.013 | 4540 | 0.130 | 16.5 |
| B2 | 6 | 0.011 | 2664 | 0.088 | 9.7 |
| D | 29 | 0.017 | 5426 | 0.114 | 19.7 |
| E | 1 | - | - | - | - |
| F | 3 | 0.007 | - | - | - |
| O | 9 | 0.003 | 2 | 0.0002 | 0.007 |

**Table S3.** Summary statistics of the core genome alignments of the different bacterial species. For each species, the number of strains, the median genome size, the size of the core genome alignment, and the number of informative SNPs in the core alignment are listed.

| Species | Strains | Genome size | Core size | Informative SNPs |
| --- | --- | --- | --- | --- |
| Escherichia coli | 92 | 4929299 | 2756541 (56%) | 247822 |
| Bacillus subtilis | 75 | 4155843 | 2341553 (56%) | 182535 |
| Helicobacter pylori | 83 | 1655288 | 850827 (51%) | 114993 |
| Mycobacterium tuberculosis | 40 | 4465985 | 4150139 (93%) | 3502 |
| Salmonella enterica | 155 | 4810980 | 2846634 (59%) | 192117 |
| Staphylococcus aureus | 95 | 2881899 | 2002833 (69%) | 73756 |

**Table S4.** SNP rates (SNPs per alignment columns) *M/L*, lower bound on the number of phylogeny changes *C*, and lower bound on the ratio of phylogeny changes to mutations *C/M* for each of the six species.

| Species | SNP rate $M/L$ | Phyl. changes $C$ | $C/M$ |
| --- | --- | --- | --- |
| Escherichia coli | 0.101 | 43575 | 0.156 |
| Bacillus subtilis | 0.113 | 40811 | 0.155 |
| Helicobacter pylori | 0.202 | 50743 | 0.295 |
| Mycobacterium tuberculosis | 0.002 | 755 | 0.078 |
| Salmonella enterica | 0.085 | 31598 | 0.131 |
| Staphylococcus aureus | 0.053 | 13215 | 0.124 |

**Table S5.**
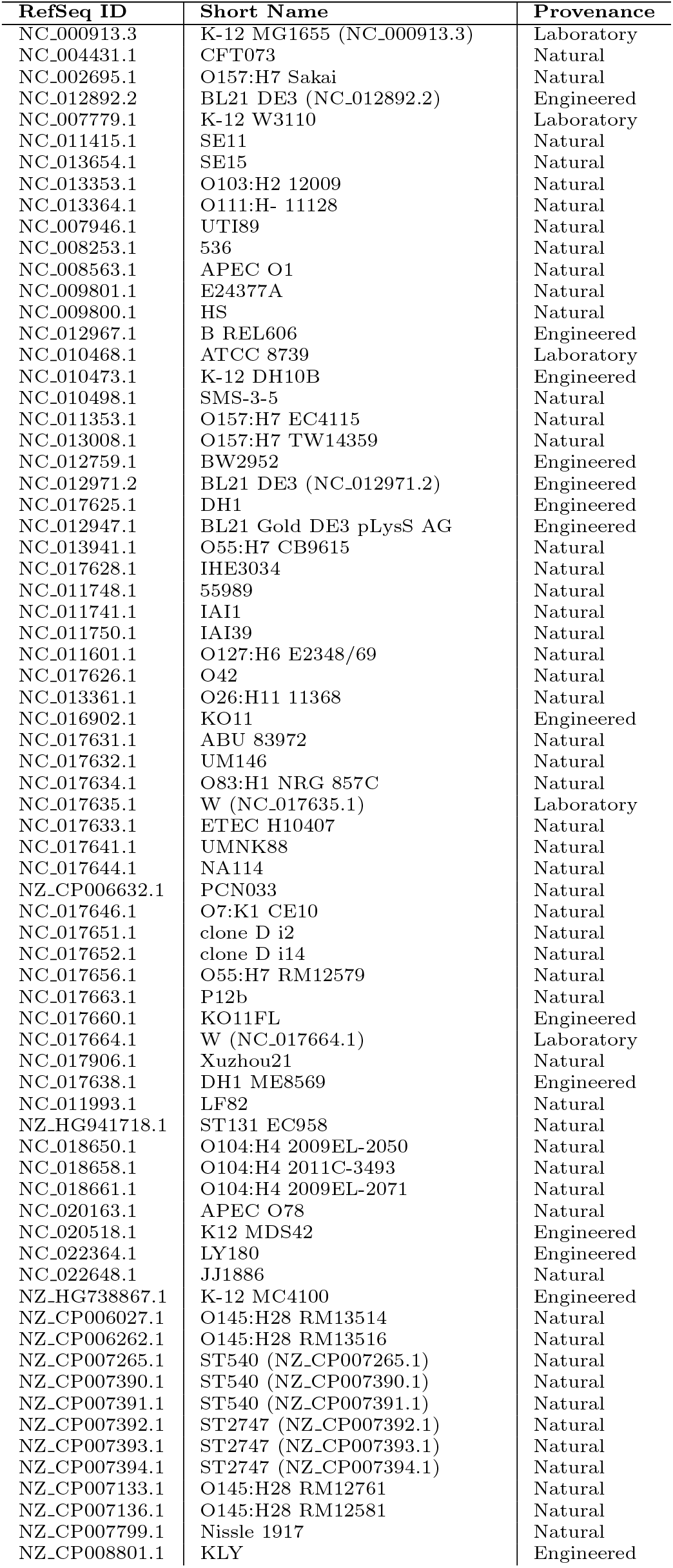
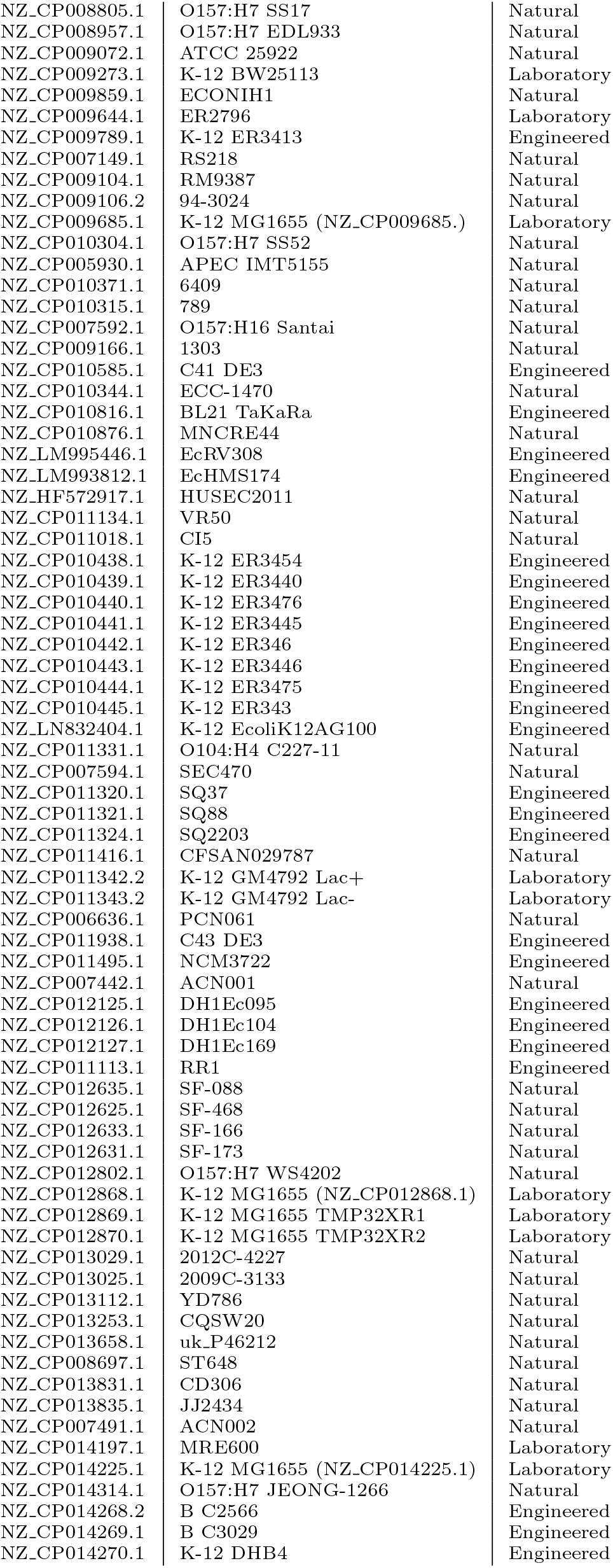
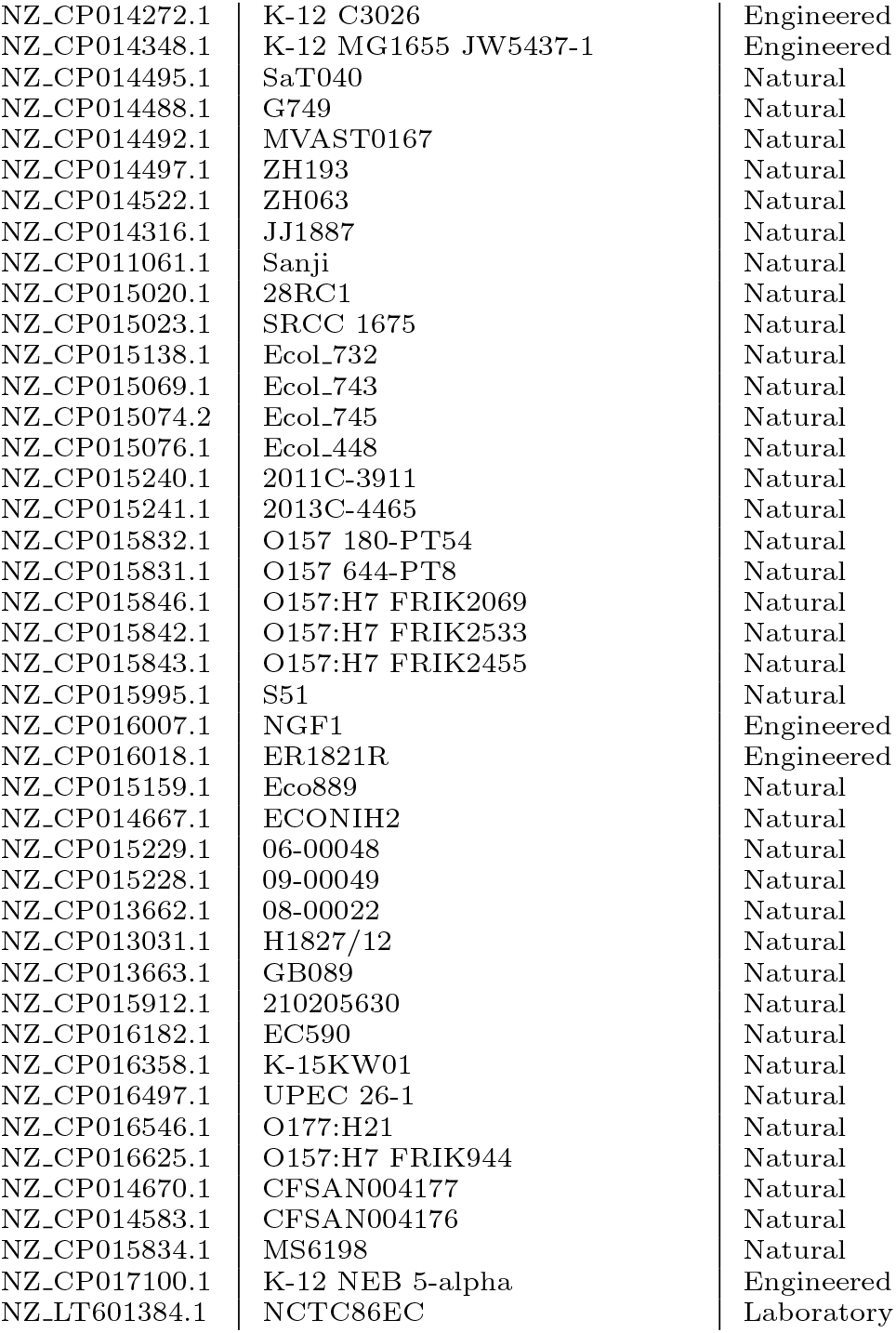
The details of the reference strains used in this work. RefSeq IDs and short names are from the NCBI nucleotide database. Provenance refers to where the strain came from: a natural strain was sequenced directly after isolation, whereas a laboratory strain has passed many generations in artificial conditions, and engineered strains have had specific changes deliberately introduced to their genomes.

**Table S6.**
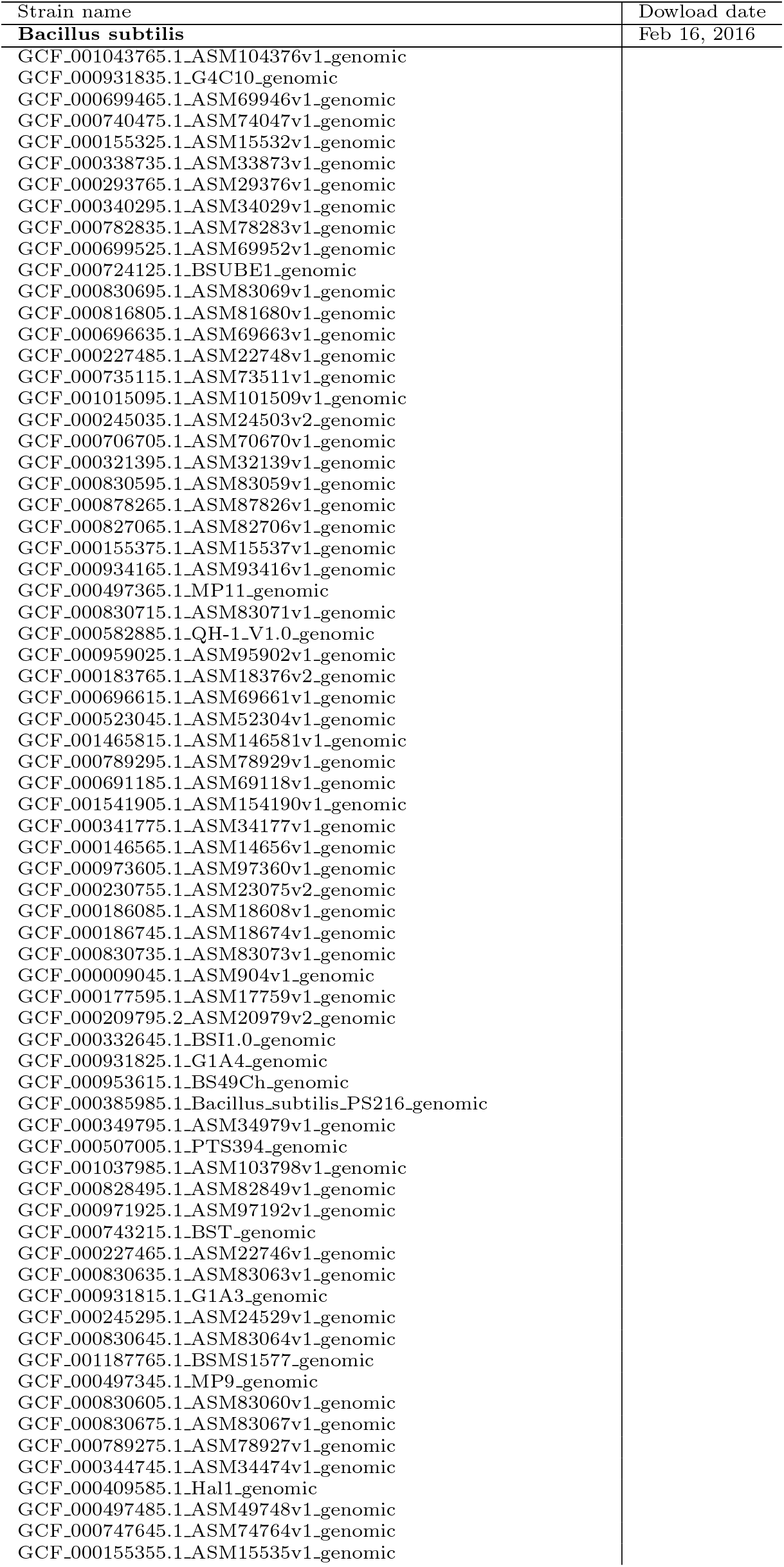
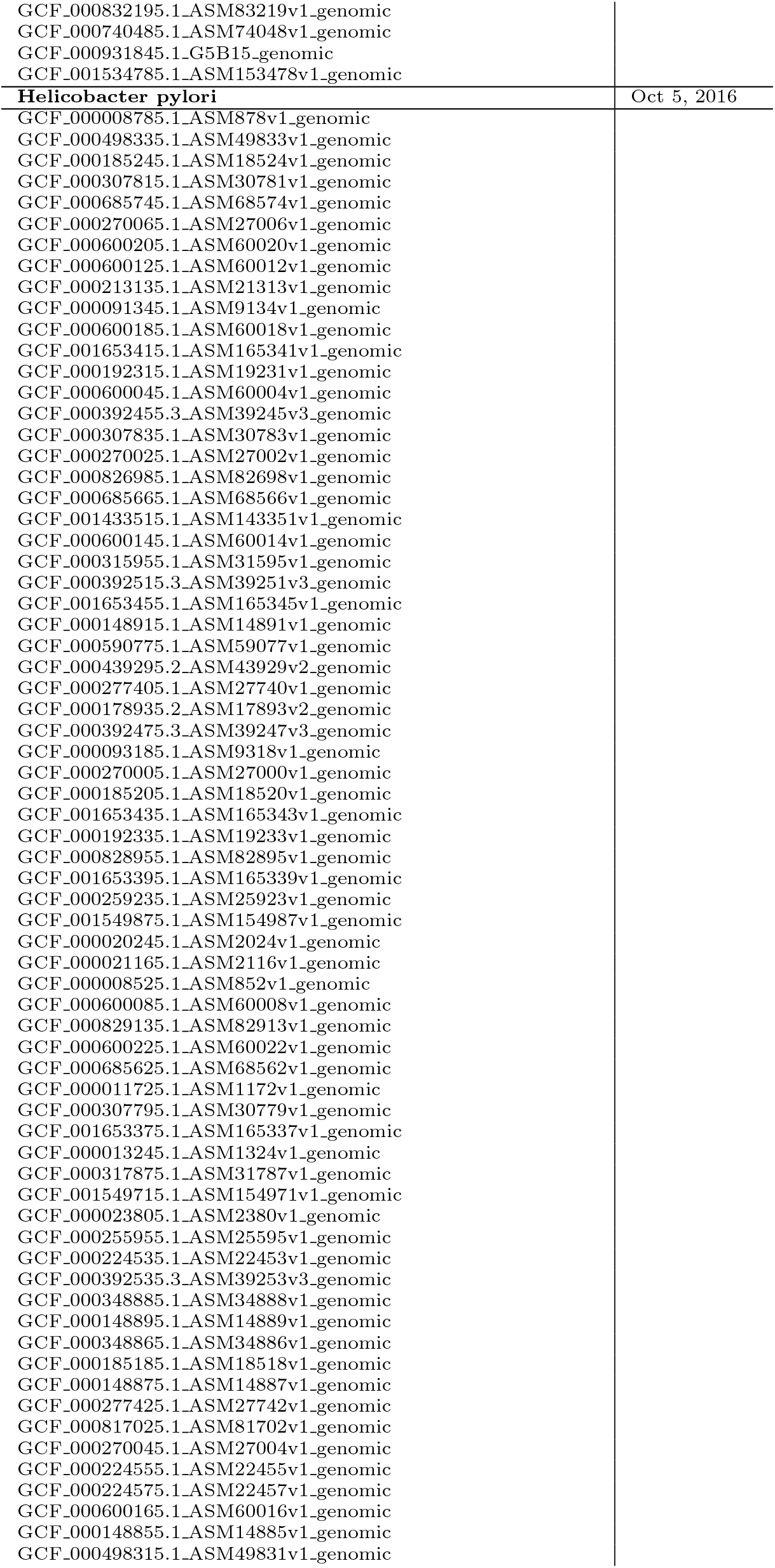
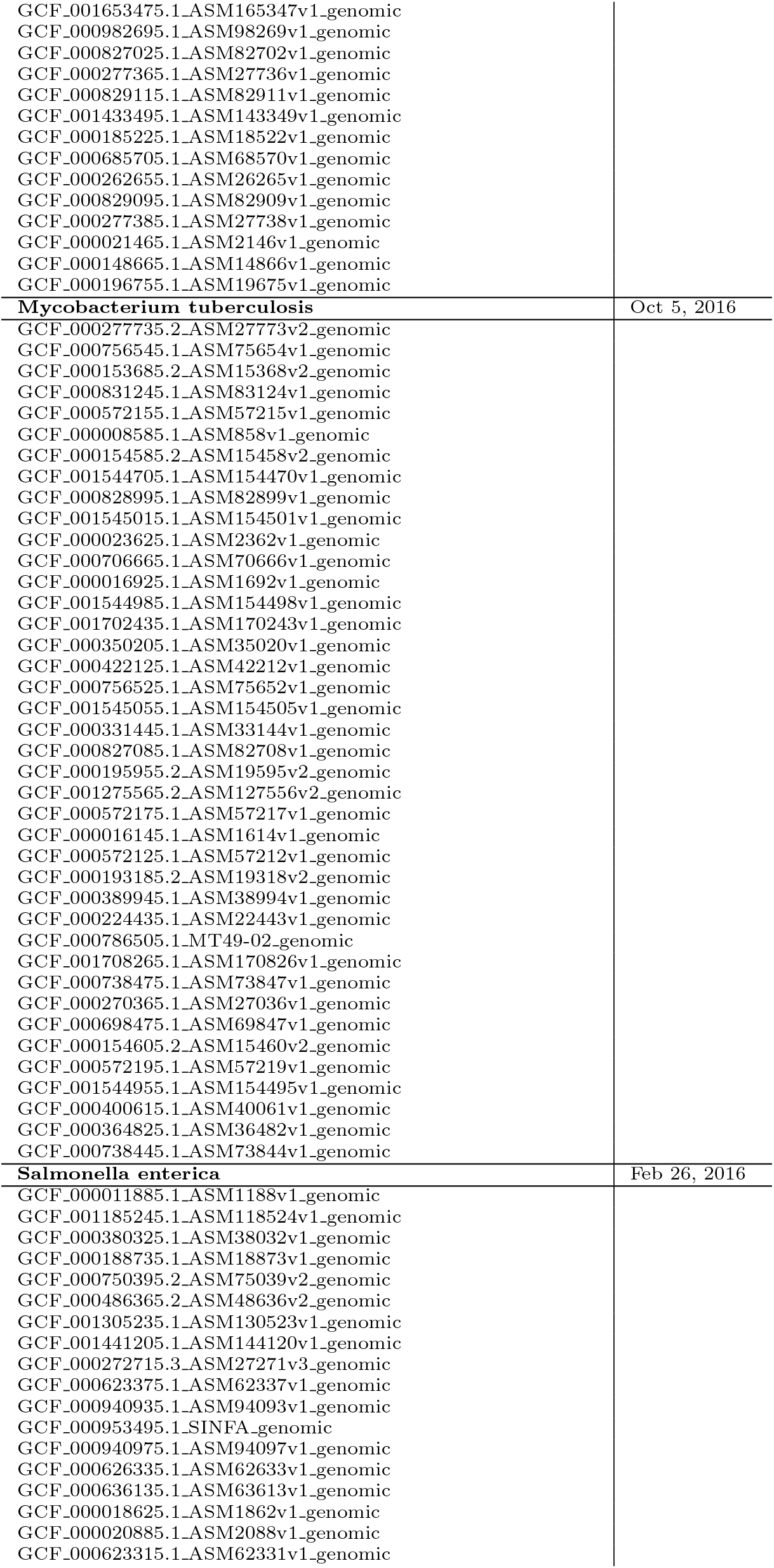
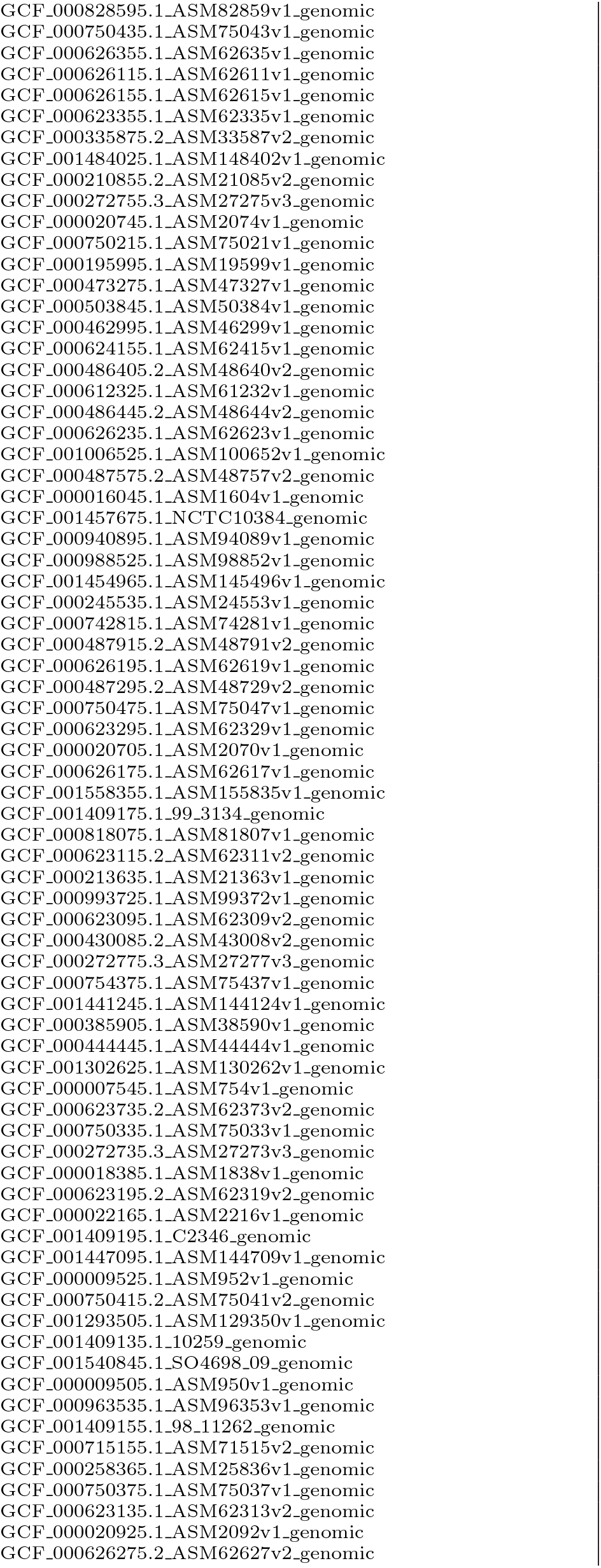
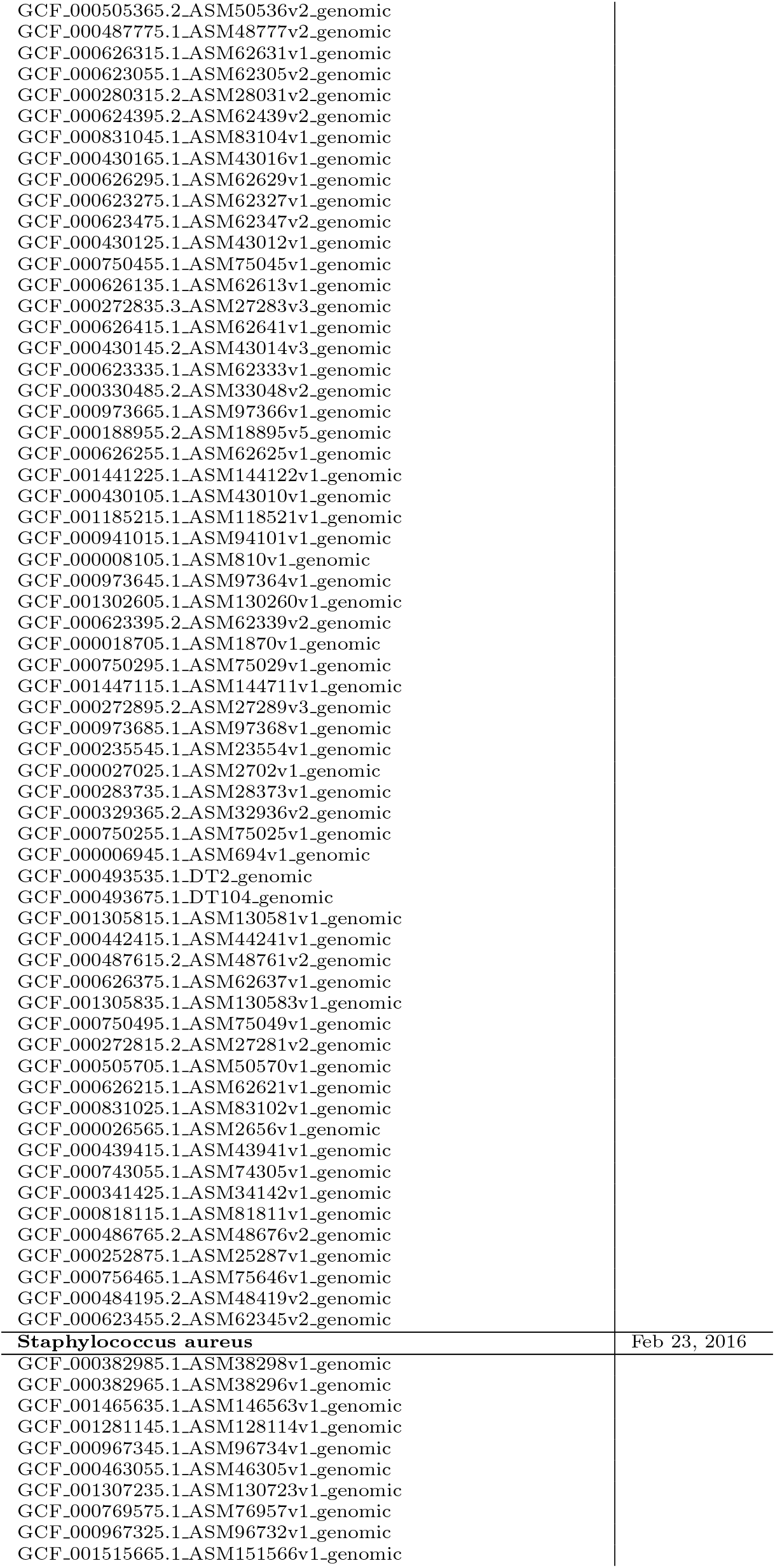
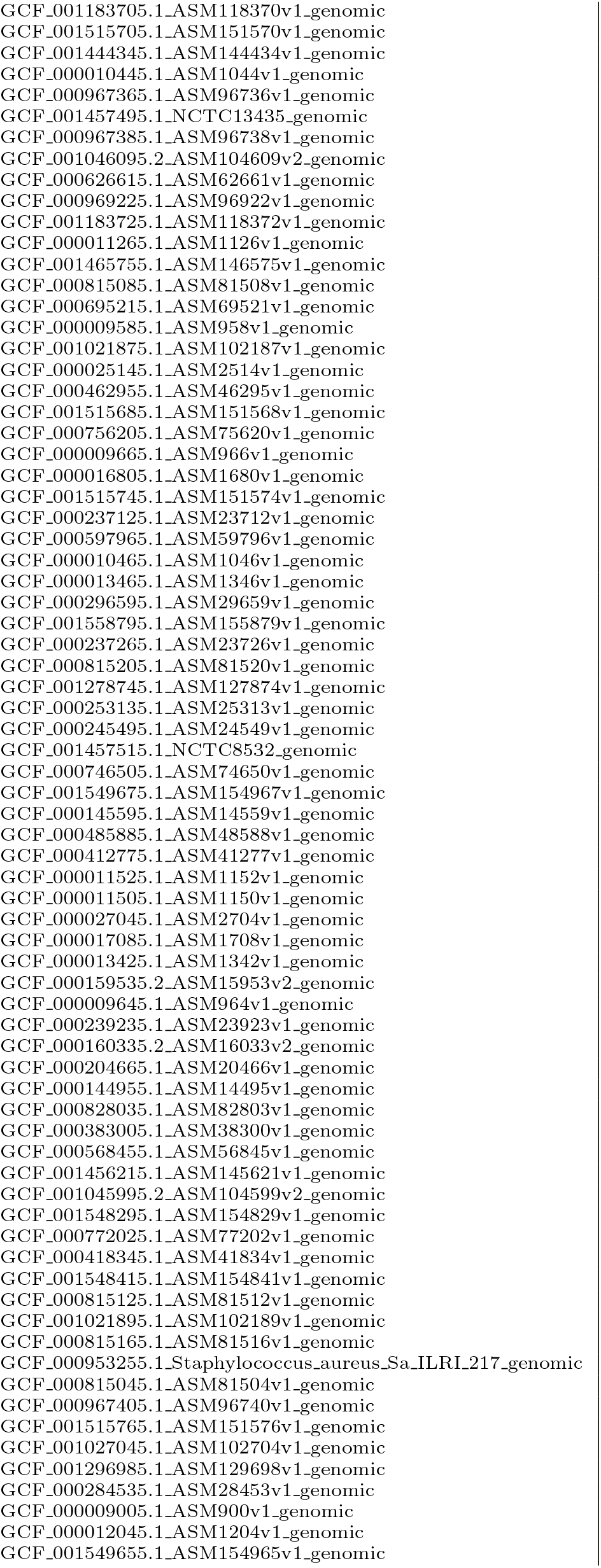
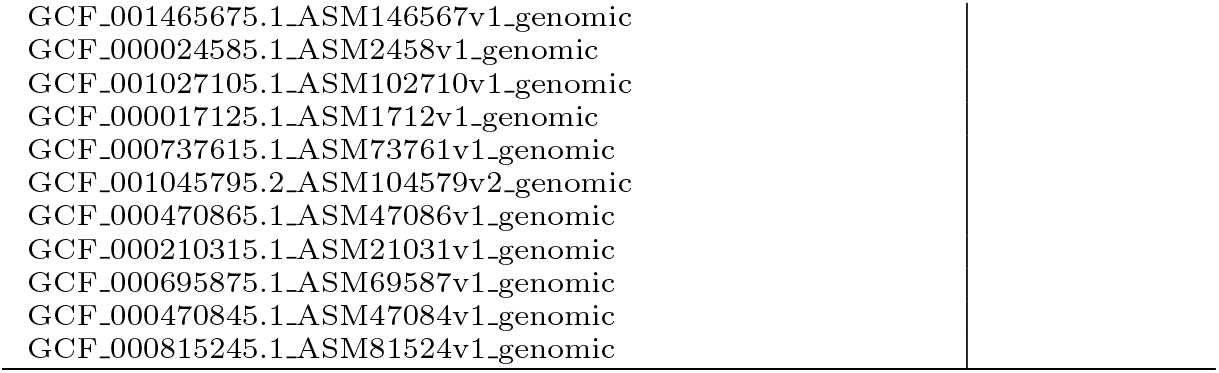
For each of the other 5 species, the table lists the strain names and the date on which the sequences were donwloaded from the database.

### S1.1 E. coli’s *n*-SNP statistics differ from *n*-SNP statistics of simulations with free recombination

In this section we systematically compare the *n*-SNP statistics observed for the *E. coli* data with *n*-SNP statistics observed for the simulations with different rates of recombination. We show that the *n*-SNP statistics for *E. coli* differ in several fundamental ways from those observed for simulations of population of freely recombining individuals.

A often considered statistic in the population genetic analysis of sexually reproducing species is the so-called site frequency spectrum, which in our terminology corresponds to the total number of *n*-SNPs *T*(*n*), i.e the total number of SNPs shared by *n* strains, as a function of *n*. Supplementary Figure S27 shows the site frequency spectrum *T*(*n*) for the *E. coli* data, the simulations without recombination, and the simulations with recombination to mutation rates ranging from *ρ/μ* = 0.001 to 10, using the 5% homoplasy-corrected alignments. To distinguish the contribution of potentially clonal SNPs falling on the core tree, the solid lines show *T*(*n*) for all SNPs

**Figure S27.**
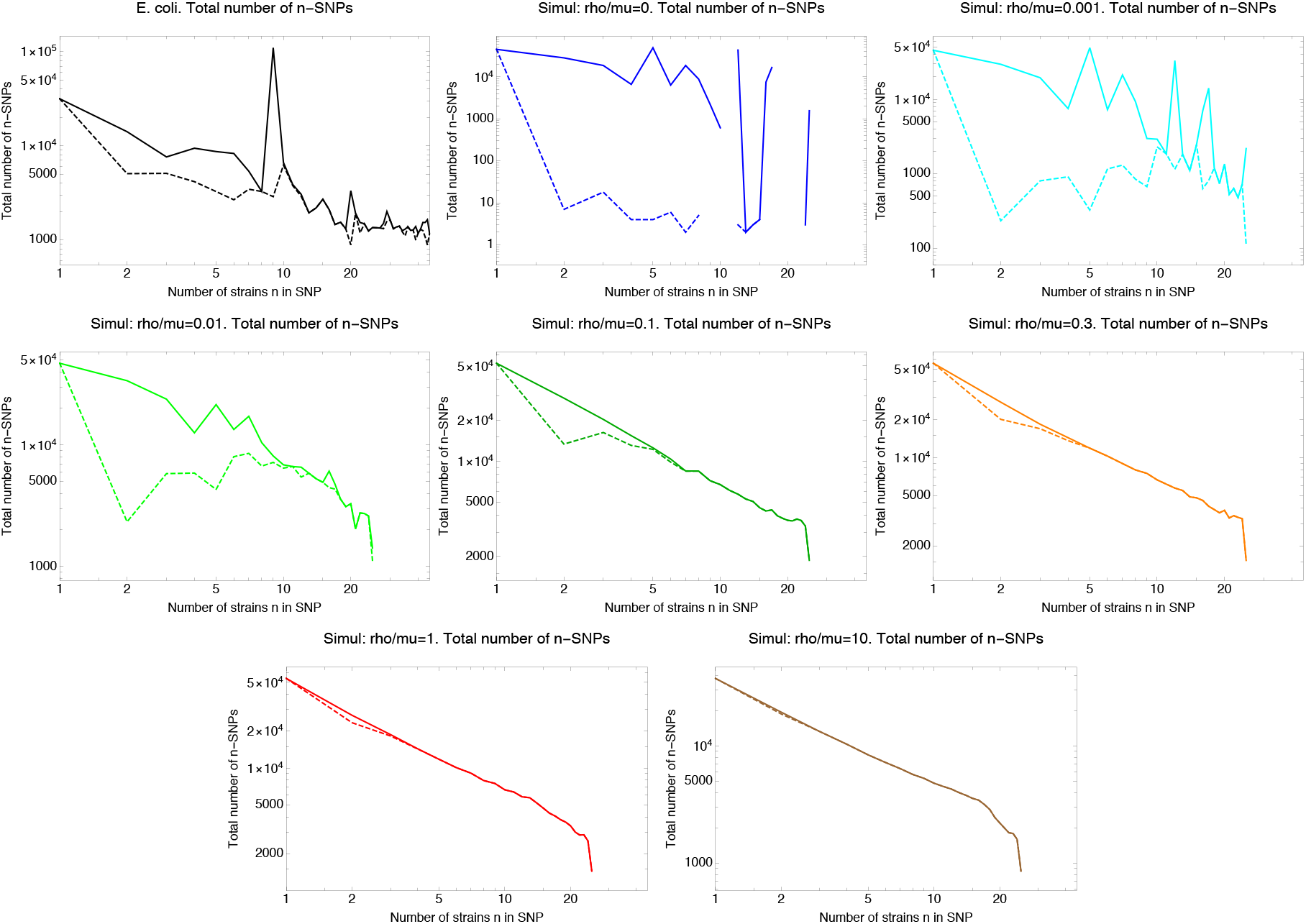
SNP frequency spectra. Total number of occurrences of *n*-SNPs, i.e. SNPs shared by *n* strains (vertical axes) as a function of *n* (horizontal axes) for the *E. coli* data (top left panel) and all simulated data with different recombination rates, where the ratio of recombination to mutation rate *ρ/μ* is indicated above each panel. The solid lines correspond to all *n*-SNPs in the 5% homoplasy-corrected alignment, whereas for the dashed lines all *n*-SNPs corresponding to branches in the core tree have been removed. Note that both axes are shown on a log-scale and for high recombination rates *ρ/μ* ≥ 0.1, the *n*-SNP frequencies are approximately proportional to 1/*n*.

We first focus on the simulation data. It is well-known that, for populations evolving under a Kingman coalescent, *T*(*n*) ∝ 1/*n*, i.e. the total frequency of SNPs shared by *n* strains falls as 1/*n* when averaged over many instantiations of the coalescent process. Indeed, for recombination rates sufficiently high that essentially all branches in the clonal history are dominated by recombination, i.e. *ρ/μ* ≥ 0.3, the observed site frequency spectrum falls exactly as 1/*n*, and removing the SNPs that fall on the core tree has virtually no effect on the *n*-SNP frequencies *T*(*n*). In contrast, for the simulations without recombination (top center panel in Suppl. Fig. S27) virtually all SNPs fall on the clonal tree, i.e. after removal of the core tree SNPs only a tiny number of SNPs remain that derive from homoplasies that escaped the homoplasy correction. Moreover, because the counts *T*(*n*) for the simulation without recombination derive from a single tree, we observe significant deviations from the average trend *T*(*n*) *x* 1/*n*, e.g. the clonal tree happens to have particularly long branches towards clades with *n* = 5, *n* = 7, and *n* = 12 strains.

For *ρ/μ* = 0.001 all loci are still predominantly clonally inherited across all branches of the tree (see Suppl. Fig. S3), which is reflected in the fact that the number of SNPs drops dramatically when the clonal SNPs are removed, and that we clearly see the peaks in *T*(*n*) at *n* = 5, *n* = 7, and *n* =12 that derive from the clonal tree. The recombination rate *ρ/μ* = 0.01 is an interesting intermediate case. While most branches of the tree are predominantly clonally inherited, a few longer branches are already mostly recombined (Suppl. Fig. S3). Reflecting this, we see that for *ρ/μ* = 0.01 clonal SNPs dominate the *n*-SNP counts for small *n*, whereas from about *n* =10 onward, the *n*-SNP counts derive mostly from recombination. At recombination rate *ρ/μ* = 0.1 clonal SNPs only affect *n*-SNP counts for *n* < 5, and for higher recombination rates the site frequencies *T*(*n*) perfectly follow the function 1/*n*.

For the *E. coli* data, we see that core tree SNPs make a significant contribution to the *n*-SNP counts for *n* ≤ 9. In particular, the peak at *n* = 9, corresponding to the extremely common SNPs toward the outgroup, as well as a smaller peak at *n* = 20, corresponding to a group of 20 extremely close strains, both disappear after removal of the core tree SNPs. From *n* =10 onward the core tree SNPs do not contribute significantly to the *n*-SNP counts *T*(*n*). Interestingly, while the counts *T*(*n*) roughly follow the 1/*n* trend until *n* = 20, for *n* > 20 the counts *T*(*n*) do not further decrease but appear virtually constant.

Next, we investigated how many different *types* of *n*-SNPs occur as a function of *n* for the different datasets, as well as the average number of occurrence of the each of the *n*-SNP types. Since the exponents of the power-law fits are determined by the geometric average of the *n*-SNP type occurrences (see Materials and Methods) we calculated the geometric average of the counts per *n*-SNP type as a function of *n* for each dataset. To focus on the *n*-SNPs associated with recombination, we used the SNPs from the 5% homoplasy-corrected alignment and removed all SNPs corresponding to branches of the core tree. The results are shown in Suppl. Fig. S28.

We first note that the number of 1-SNP types is 50 for all the simulated data (which consisted of a sample of *n* = 50) genomes. That is 1-SNPs are observed that are exclusive to each of the 50 genomes. However, the number of 1-SNP types observed for the *E. coli* data is 55, i.e. much less than the 92 possible types given that there are 92 strains. This reflect the fact that, in our collection of wild *E. coli* isolates, there are some groups of extremely closely-related strains with identical core genomes. Note that such closely related strains are unlikely to occur for random samples from a Kingman coalescent.

Second, we see that, for the *E. coli* data, the number of *n*-SNP types increases with *n*, saturating at around 500 — 800 unique *n*-SNP types for each *n* at *n* ≥ 5. In the right panel of Suppl. Fig. S28 we see that, as the number of *n*-SNP types increases, the geometric mean frequency of the *n*-SNP types decreases smoothly. This smooth decrease of the geometric mean mirrors the increasing exponents of the *n*-SNP power-laws. None of the simulation data show patterns that mick what is observed for the *E. coli* data. First, at the lowest recombination rate *ρ/μ* = 0.001 the number of *n*-SNP types is low and decreasing with *n*. Second, for recombination rate *ρ/μ* = 0.01, the number of SNP types increase more gradually with *n*, saturating at about 200 *n*-SNP types for large *n*. However, in contrast to what we observe for the *E. coli* data, the geometric mean number of occurrences for the simulations with *ρ/μ* = 0.01 is much larger and virtually constant for *n* ≥ 2. At higher recombination rates *ρ/μ* ≥ 0.1, the number of *n*-SNP types rises quickly with *n*, reaching a maximum of thousands of unique *n*-SNP types between *n* = 3 and *n* = 5, and then decreases with larger *n*. At the same time, the geometric mean of the number of occurrences per type decreases quickly with *n* and stays at low values of about 3 for *ρ/μ* = 0.1 and close to 1 for the highest recombination rates. That is, when recombination completely dominates, and there is no population structure by construction, almost all SNP types are distinct, occurring only once or a few times.

**Figure S28.**
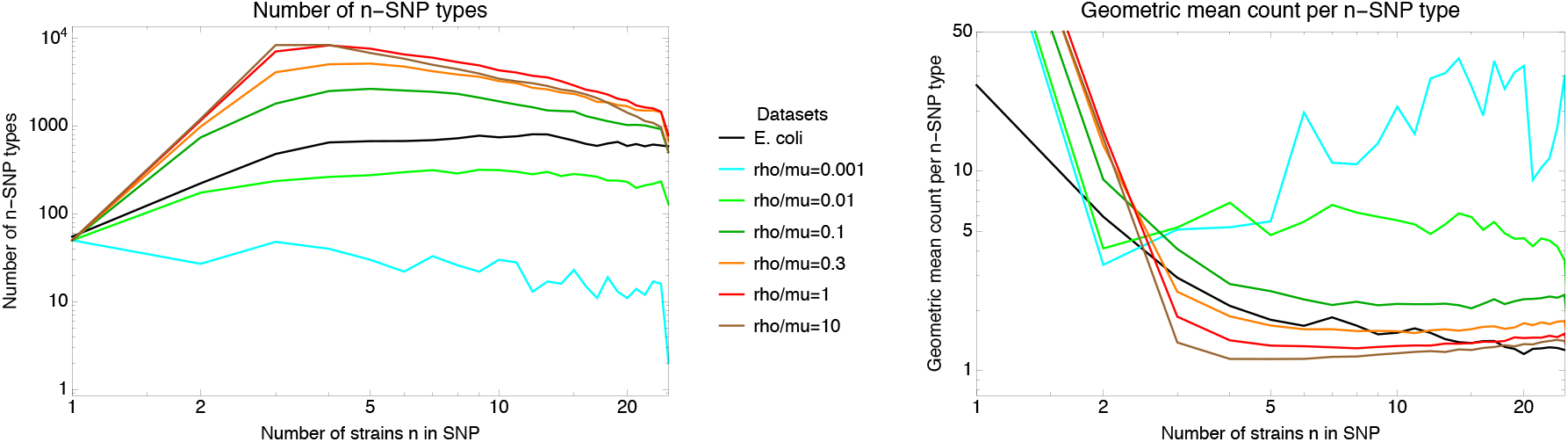
Diversity of *n*-SNPs. **Left panel**: Number of *n*-SNP types (vertical axis) as a function of *n* (horizontal axis) for the *E. coli* data (black line) and for simulations with different recombination to mutation rates *ρ/μ* (colored lines, see legend). Both axes are shown on a logarithmic scale. **Right panel**: Geometric mean of the number of occurrences of *n*-SNP types (vertical axis) as a function of *n* for the *E. coli* data (black line) and for simulations with different recombination to mutation rates *ρ/μ* (colored lines, see legend). Both axes are shown on a logarithmic scale. Note that, in order to better show the heights of the curves for the different datasets the vertical axis is clipped at 50. For the simulated datasets the geometric mean of the 1-SNP counts runs from 400 to 1000.

Note that our previous analysis has shown that the number of phylogeny changes per SNP column *C/M* for *E. coli* is similar to that observed for *ρ/μ* = 1. However, as Suppl. Fig. S28 shows, at such high recombination rates one would expect much higher *n*-SNP diversity still than is observed for the *E. coli* data. The fact that the *E. coli* data shows evidence of high recombination on the one hand, in combination with lower SNP diversity, is consistent with SNP diversity being constrained by population structure.

We next compared the approximately power-law *n*-SNP frequency distributions that we observed for *E. coli*, with the *n*-SNP frequency distributions observed for the simulation data. Supplementary Figure S29 shows the *n*-SNP distributions for 6 different values of *n* ranging from *n* = 2 to *n* =12.

Whereas, for the *E. coli* data, all distributions are approximately power-law, almost all of the distributions for the simulated data are clearly bending downwards in the log-log plot, often severely so. Long tailed distributions are only observed for the very low recombination rates *ρ/μ* = 0.001 and *ρ/μ* = 0.01, for which we have seen that *n*-SNPs are still largely dominated by clonal SNPs at small values of *n*. However, as we have seen above, such low recombination rates are not consistent with the *E. coli* data. In addition, as we will see in more detail below, the exponents of the *n*-SNP distributions at recombination rates *ρ/μ* = 0.001 and *ρ/μ* = 0.01 are much lower than is observed for *E. coli*, and the average number of occurrence per *n*-SNP type are significantly higher than observed for *E. coli* (Suppl Fig. S28, right panel).

All *n*-SNP distributions at recombination rates *ρ/μ* ≥ 0.1 differ clearly from power-laws. That is, instead of long-tailed distributions most of these *n*-SNP frequencies fall within a fairly narrow range. For example, at *ρ/μ* = 1 most 2-SNPs have frequencies between 10 — 50, most 3-SNPs have frequencies between 2 — 20, and 4-SNPS that occur more than 10 times are very rare.

Although the *n*-SNP distributions at mutation rates *ρ/μ* ≥ 0.1 cannot reasonably be fitted to powerlaws, we can still determine the exponents of the maximum likelihood fits since these only depend on the geometric average number of occurrences of the *n*-SNP types (see Materials and Methods). Supplementary Figure S30 shows the resulting exponents for the simulated data, as well as for the *E. coli* data for reference.

**Figure S29.**
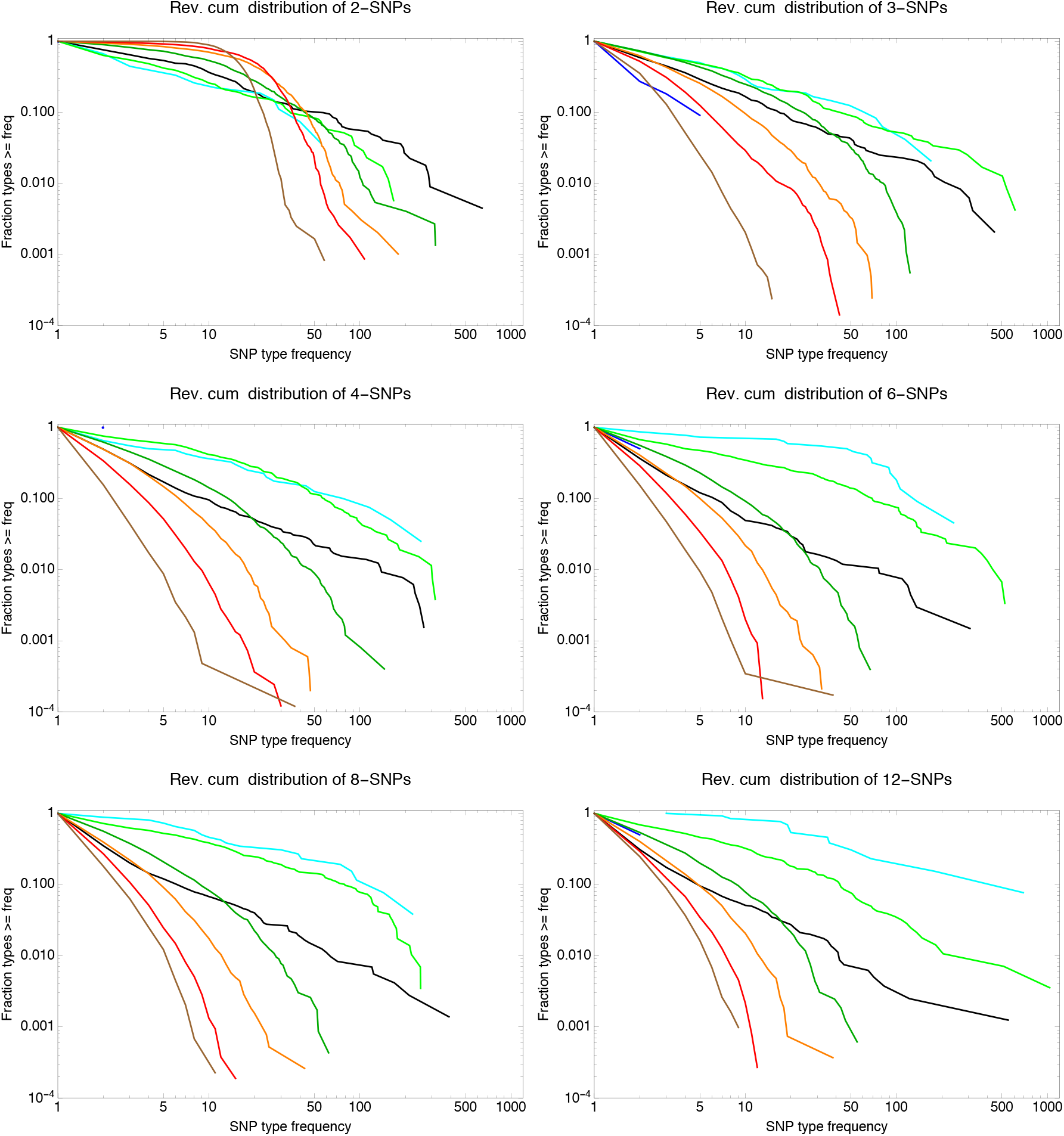
Example *n*-SNP distributions for *E. coli* (black lines) as well as for the simulations with different rates of recombination with *ρ/μ* = 0 shown in blue, *ρ/μ* = 0.001 in cyan, *ρ/μ* = 0.01 in light green, *ρ/μ* = 0.1 in dark green, *ρ/μ* = 0.3 in orange, *ρ/μ* = 1 in red, and *ρ/μ* = 10 in brown. Each panel corresponds to the observed *n*-SNP distributions with the value of *n* indicated at the top of each panel. All axes are shown on logarithmic scales.

**Figure S30.**
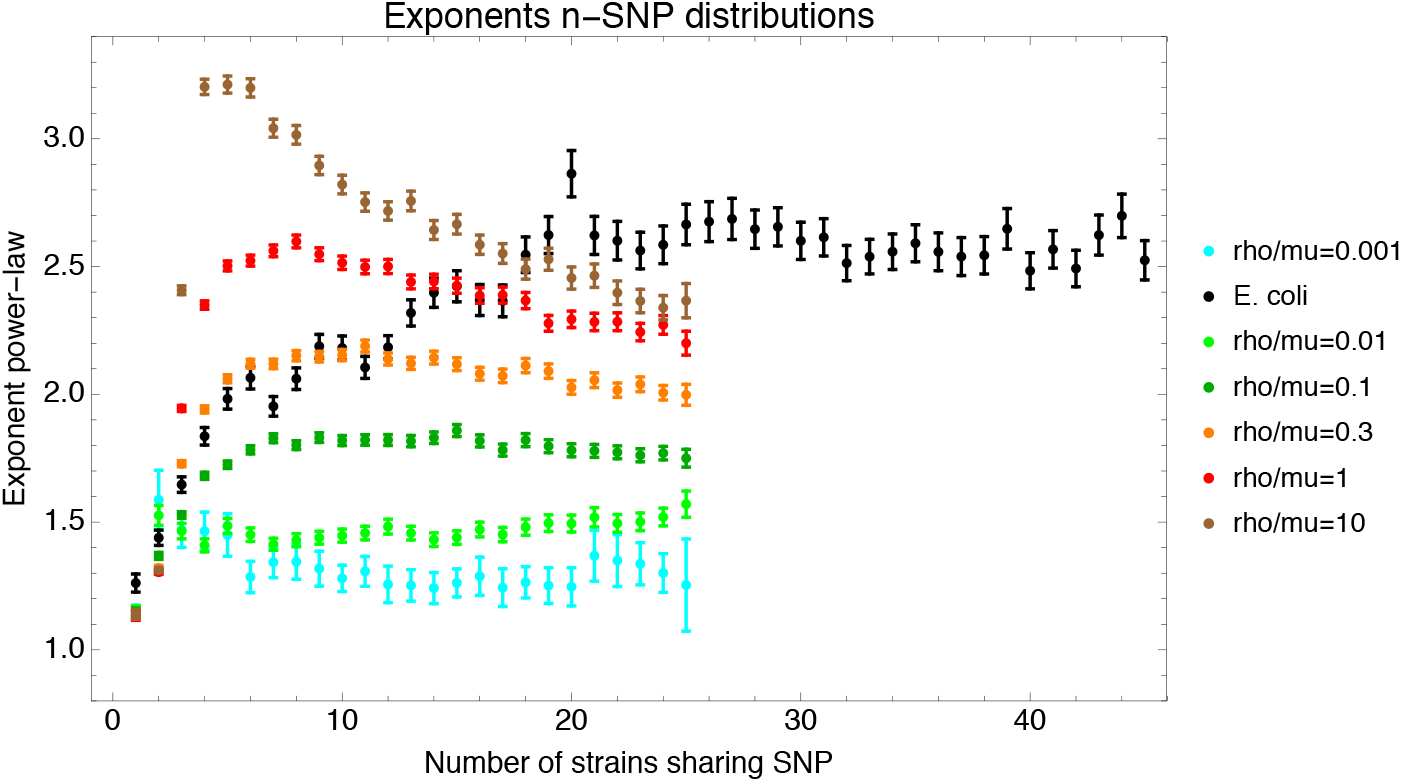
Fitted exponents of the *n*-SNP distributions for the *E. coli* data (black) and the data from the simulations with different recombination rates (colors, see legend). The bars show the fitted exponent plus and minus one standard-deviation of the posterior distribution.

We see that, for the simulation data, exponents increase systematically with recombination rate. For low recombination rates *ρ/μ* = 0.001 and *ρ/μ* = 0.01 all exponents are below 1.5, whereas for recombination rate *ρ/μ* = 1 the exponents lie in the range 2.3 — 2.6. In contrast to the *E. coli* data, for which exponents increase smoothly from about 1.25 to 2.7 as *n* increases, for the simulations the exponents tend to be more constant as a function of *n*.

In summary, none of the *n*-SNP statistics of the data from simulations matches the *n*-SNP statistics observed for the *E. coli* data. For example, while fairly long-tailed distributions of *n*-SNPs are observed at very low recombination rates, the exponents of these distributions are much lower than found for the *E. coli* data, and the *n*-SNP diversity is lower than observed for the *E. coli* data. In addition, such low recombination rates are inconsistent with all of our other analyses of the *E. coli* data. In contrast, while high *n*-SNP diversity is observed at higher recombination rates, there the *n*-SNP distributions are not long tailed, but instead *n*-SNP frequencies vary only over a relatively narrow range.

### S1.2 Detailed SNP statistics for Phylogroup B2

In the main text, we illustrated 2-SNPs using strain A1 (Fig. 8), which is part of the phylogroup B2. The phylogroup B2, which is represented by the 6 strains A1, A2, A7, A11, B2, and D8 in our dataset, is small enough to allow for an illustrative discussion of what our various analyses shows us about the role of recombination in the evolutionary history of these strains.

#### S1.2.1 Pairwise analysis for the sextet of strains (A1,A2,A7,A11,B2,D8)

Supplementary table S7 shows the pairwise divergences between strain A1 and the other strains of this phylogroup.

**Table S7.** Pairwise divergence strain A1 with other strains of phylogroup B2.

| Strain | divergence |
| --- | --- |
| B2 | 0.00626 |
| A7 | 0.00627 |
| A11 | 0.00702 |
| D8 | 0.00729 |
| A2 | 0.00778 |

Note that these divergences are in the regime where, in general, most of the pairwise alignment has already been overwritten by recombination (see Fig. 2G and note that the critical divergence at which 50% of the genome has recombined is at 0.0032). Indeed, if one looks at the pairwise alignments of A1 with these strains (Fig. S31), one finds that for each of these 5 strains there is very little ancestrally inherited DNA left (Suppl. Fig. S31).

**Figure S31.**
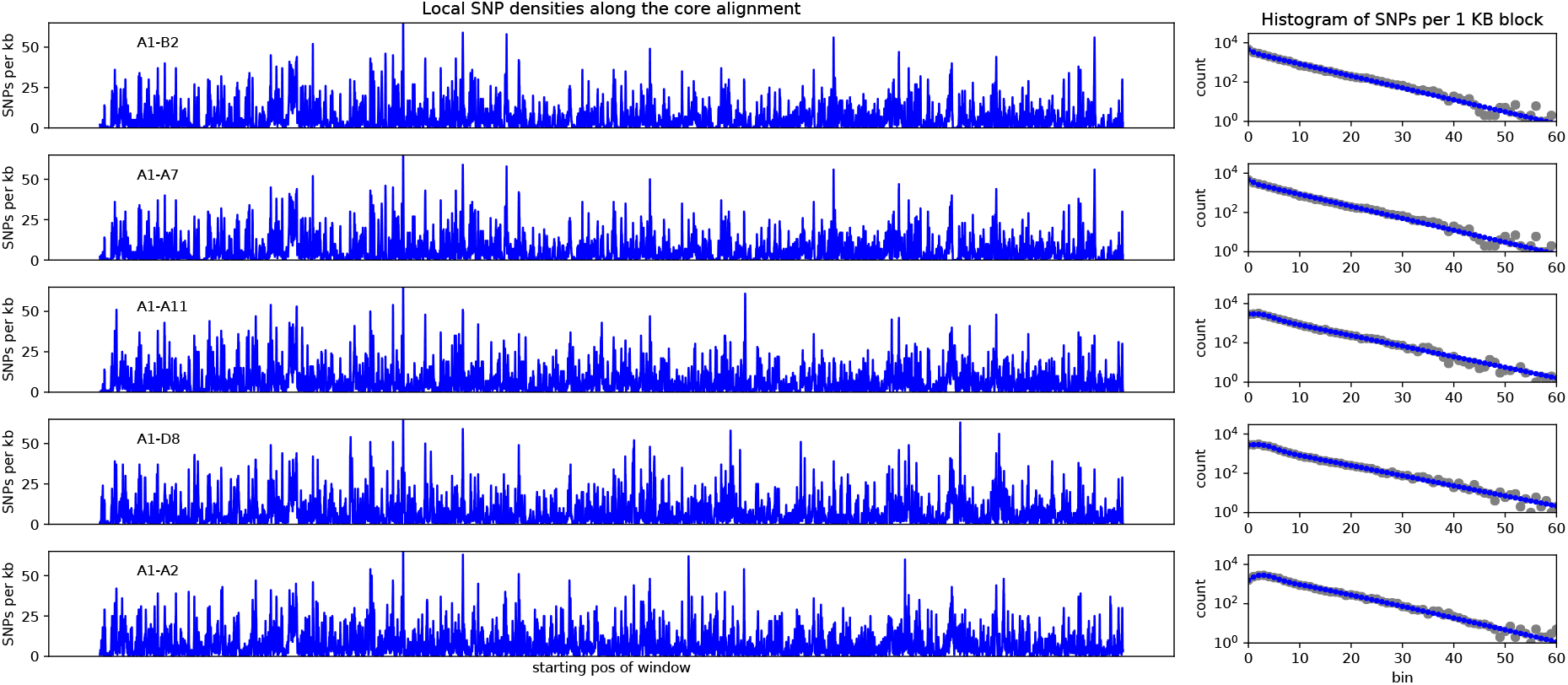
**Left panels**: SNP densities (SNPs per kilobase) along the core genome for the 5 pairs of strains (A1-B2), (A1-A7), (A1-A11), (A1-D8), and (A1-A2). **Right panels**: Corresponding histograms for the number of SNPs per kilobase (dots) together with fits of the mixture model. Note the vertical axis is on a logarithmic scale.

This is in fact true for *all* pairs of the phylogroup B2, with the exception of the pair (A7,B2). The divergence of (A7,B2) is less than 10^-4^ and their alignment is fully clonal, i.e. A7 and B2 share a recent common ancestor. However, according to the analysis of SNP densities along the pairwise alignments, all other pairs in this clade are close to fully recombined. Note that this also means that the pairwise divergences are dominated by mutations in recombined regions, not by ancestrally inherited mutations (see Fig. 2H). Thus, this pairwise analysis already rejects that the core genome phylogeny for this phylogroup corresponds to the clonal phylogeny.

#### S1.2.2 Phylogeny changes along the core alignment of phylogroup B2

That recombination is pervasive within this set of strains with relative low divergence is confirmed by the phylogenetic inconsistencies along the alignment made from the core genomes of just this sextet of strains (A1,A2,A7,A11,B2,D8). In particular, in the 5% homoplasy-corrected alignment there are *M* = 30’196 SNPs and at least *C* = 2664 changes in phylogeny along this core genome alignment. This means that the lower bound on the ratio of the number of phylogeny changes to mutation events is *C/M* = 0.088, i.e. a break on average every 11 SNPs. Supplementary Fig. S32 shows the distribution of segment length between phylogeny breaks (either counting the number of consecutive SNPs or total length of the segments) which shows that breaks typically occur within 10 SNPs and within a few hundred base pairs.

**Figure S32.**
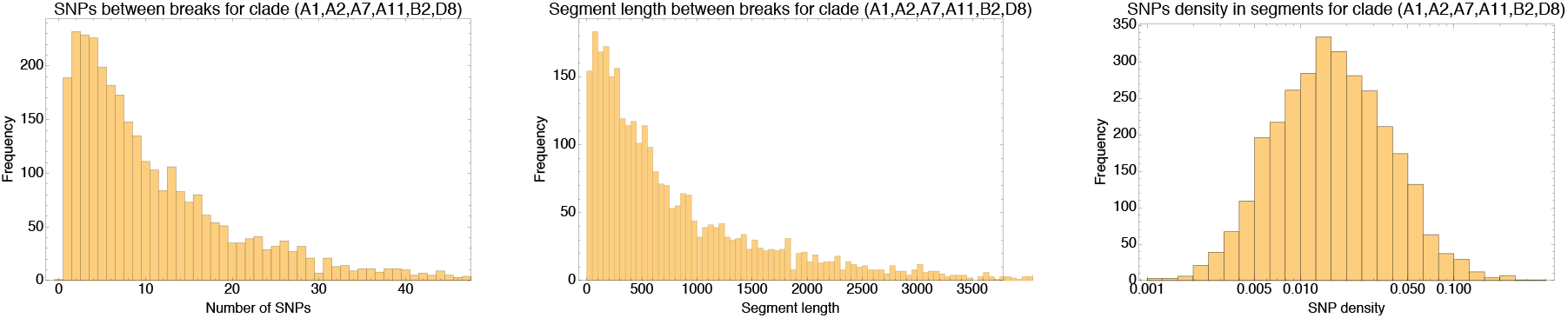
**Left:** Histogram of the number of consecutive SNPs before an inconsistency in the core genome alignment of the sextet of strains (A1,A2,A7,A11,B2,D8) of phylogroup B2. Note that the vertical axis corresponds to the number of segments with the corresponding number of consecutive SNPs. **Middle:** Histogram of the length of segments without phylogeny breaks. **Right:** Histogram of the SNP density across the segments that are consistent with a single phylogeny (shown on a logarithmic axis). Note that the SNP densities vary by two orders of magnitude.

Supplementary Fig. S32 also shows (right panel) that the SNP density varies by as much as hundredfold across the 2664 segments, consistent with the fact that the transfered fragments are themselves mosaics of previous recombination events.

#### S1.2.3 SNP types for the strains in phylogroup B2

Another indication that these single-phylogeny segments are the result of recombination comes from looking at what the actual SNP types are that occur in the segments. Supplementary table S8 shows, for the ten longest segments, the lengths, the total number of SNPs, and SNP-types that occur in each segment.

Note that the SNP-types that occur in these longest unbroken segments are not only almost all inconsistent with the core genome phylogeny, they are also almost all inconsistent with each other. That is, each of these segments suggests a different phylogeny. Note also that, in each segment, there are typically multiple SNPs of the same type.

Finally, the fact that the alignment of this sextet is a mixture of segments that follow different phylogenies is also confirmed by the overall relative frequencies of the SNP-types that are observed for these strains. Supplementary table S9 shows the most common 2-SNPs, 3-SNPs and 4-SNPS together with their number of occurrences, for this sextet of strains. Note that these are SNP-types obtained from the entire alignment. That is, a 4-SNP (A1 A11 A7 B2) means that these strains share a nucleotide that differs from the nucleotide that all other 88 strains have.

**Table S8.** Lengths, total SNP count, and SNP types (i.e. the strains that carry the minority allele) for the ten longest segments (in terms of number of consecutive SNPs) along the core genome of the sextet (A1,A2,A7,A11,B2,D8) of phylogroup B2.

| Length segment | Number of SNPs | SNP types |
| --- | --- | --- |
| 9698 | 109 | (A7, B2) (A1, A7, B2) (A2, D8) |
| 5672 | 100 | (A7,B2) (A1,A2) (A11,D8) |
| 2068 | 100 | (A2,D8) (A11,A7,B2) |
| 2255 | 95 | (A7,B2) (A11,A2) (A1,A11,A2) |
| 11726 | 95 | (A7,B2) (A2,A7,B2) (A1,D8) |
| 11790 | 93 | (A7,B2) (A2,D8) |
| 3614 | 86 | (A11,A2) |
| 4564 | 86 | (A11,A7,B2) (A1,A2) |
| 2890 | 75 | (A7,B2) (A1,A7,B2) |
| 2390 | 71 | (A1,A11) |

**Table S9.**
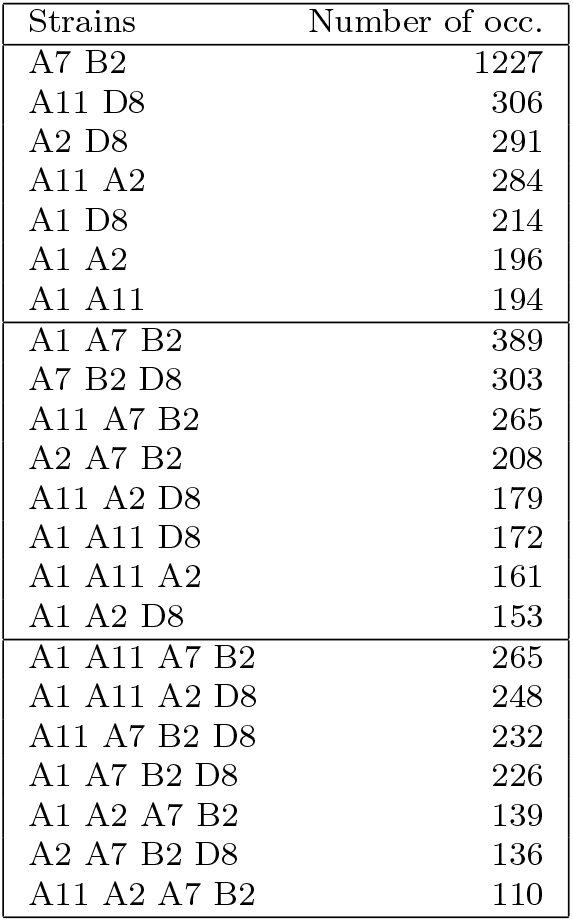
All 2-SNPs, 3-SNPs, and 4-SNPs involving strains from the sextet (A1 A2 A7 A11 B2 D8), that occur at least 100 times, sorted by their frequency of occurrence.

Supplementary table S9 shows that, apart from the clonal pair (A7 B2), which is consistently supported by the observed SNPs, the others strains occur at similar frequencies in different mutually inconsistent combinations. e.g. the 4 most common 3-SNPs all consist of the pair (A7 B2) with a different third strain. This diversity of phylogenetic partners for each strain is well summarized by the entropy profiles of these 6 strains, as shown in Suppl. Fig. S33.

**Figure S33.**
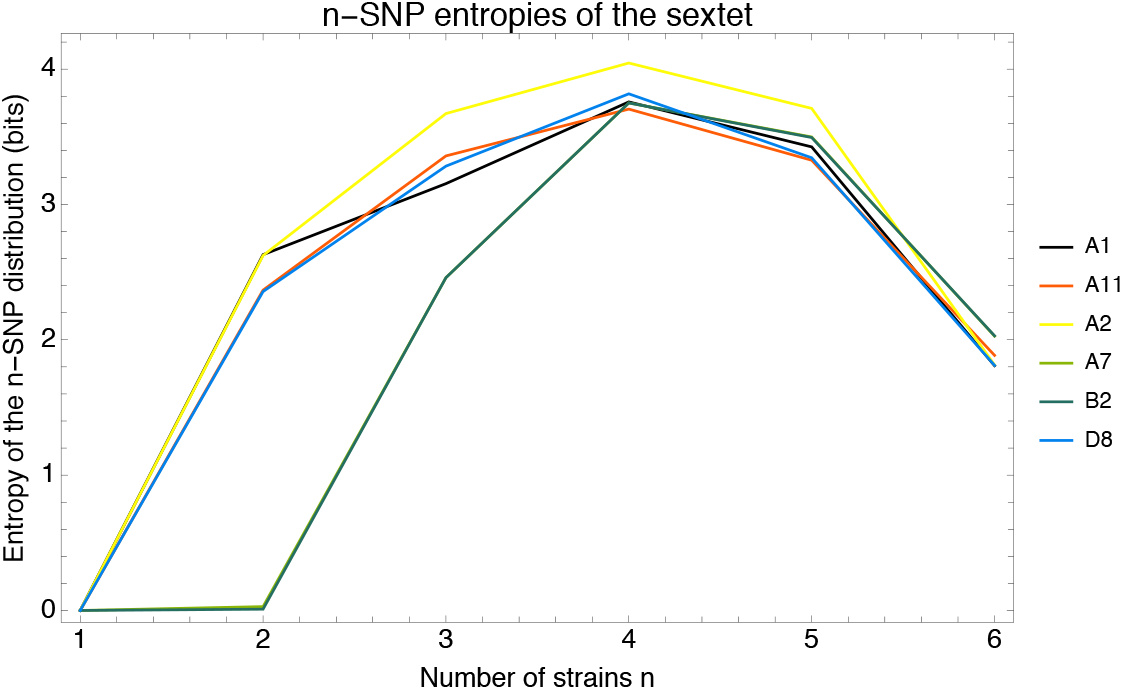
Entropy profiles for the 6 strains (A1 A2 A7 A11 B2 D8).

First, note that the strains of the pair (A7 B2) have virtually identical entropy profiles, indicating that they had a clonal ancestor before much recombination took place, i.e. their entropy profiles correspond to the entropy profile of their common ancestor. They also have zero pair entropy because they only occur in pairs with each other. However, all other strains have pair entropies around 2.5, which is equivalent to occurring with equal frequency in 5 — 6 different pairs. For quartets the entropies range from 3.6 to 4, equivalent to 12 — 16 equally frequent quartets. That is, there is a diverse collection of phylogenetic relationships in which these 6 strains occur.

#### S1.2.4 Summary: The phylogeny of strains (A1,A2,A7,A11,B2,D8)

In summary, all statistics show that even for the relative close sextet of strains (A1 A2 A7 A11 B2 D8), the only clear clonal signal left is the close pair (A7 B2). The alignments between all other pairs have been mostly overwritten by recombination, the phylogeny changes thousands of times along the core genome alignment, and each strain occurs in a diverse collection of multiply inconsistent *n*-SNPs with the other strains. However, as shown in the bottom panel of Fig. 1 of the main text, the branches in the core tree of this sextet are well supported when a tree is constructed from sufficiently many loci. That is, the core genome phylogeny is just the best compromise capturing the statistics with which different phylogenetic patterns appear, and one reliably converges to this best compromise when sufficiently many loci are taken into account.

